# Calibrating and documenting host-switching and evolution of incompatibility loci for two closely related *Wolbachia* clades

**DOI:** 10.64898/2026.02.13.705778

**Authors:** J. Dylan Shropshire, William R. Conner, Dan Vanderpool, Ary A. Hoffmann, Michael Turelli, Brandon S. Cooper

## Abstract

Maternally inherited *Wolbachia* alphaproteobacteria are the most common arthropod endosymbionts.

Often *Wolbachia* spread to high frequencies through cytoplasmic incompatibility, in which *cif* loci act through sperm to kill embryos lacking *Wolbachia*. Closely related *Wolbachia* with diverse *cif* loci often associate with anciently diverged hosts, but the timescale of associations remains uncertain. We produce new calibrations based on filarial nematodes with vertically inherited *Wolbachia* that codiverge with their hosts. Applying these calibrations to *Wolbachia* variants closely related to pathogen-blocking *w*Mel from *Drosophila melanogaster*, we demonstrate that over a timescale of 1–2 million years, a core set of single-copy *Wolbachia* loci evolve largely through bifurcation rather than by gene exchange with distant *Wolbachia*. Dating bifurcating core genomes, we show that “*w*Mel-like” *Wolbachia* diverged 2.1×10^5^–2.4×10^6^ years inhabit dipteran and hymenopteran hosts diverged more than 10^8^ years. Previous published analysis of variants related to *w*Ri from *D. simulans*, the first *Wolbachia* found in a drosophilid, concluded that “*w*Ri-like” *Wolbachia* spread among different *Drosophila* in tens of thousands of years.

However, our new calibrations suggest these estimates from a mutation-based calibration underestimated *w*Ri-like spread by about a factor of seven. In addition, *cif* exchanges between *w*Mel-like and *w*Ri-like *Wolbachia* genomes have occurred over ∼10^4^–10^6^ years. Comparing intact *cif* loci found in various *Wolbachia*, we find function-preserving selection in their evolution. We discuss these results in light of theoretical predictions concerning selection on cytoplasmic incompatibility phenotypes within and among host lineages. The *w*Mel variants analyzed may offer new options for *Wolbachia*-based biocontrol efforts.

**Article Summary:** Maternally transmitted *Wolbachia* endosymbionts associate with many arthropods. Closely related *Wolbachia* are known to occupy anciently diverged hosts, but the timescale of associations remains uncertain. Using genomic data from obligate *Wolbachia* codiverging with filarial nematodes, we calibrate *Wolbachia* molecular evolution. *w*Mel-like variants that diverged about 10^4^–10^6^ years ago occur in hosts that diverged more than 10^8^ years ago. Incompatibility loci move even more rapidly among these *Wolbachia* genomes. As these variants diverge, selection occurred on loci involved in both inducing and protecting against cytoplasmic incompatibility. These results elucidate *Wolbachia* host-range and genome dynamics and provide potential material for applications.

## Introduction

Maternally inherited *Wolbachia* bacteria were first discovered in the ovaries of the mosquito *Culex pipiens* (Hertig and Wolbach 1924). *Wolbachia* are now estimated to occur in about half of all terrestrial arthropod species (Weinert *et al*. 2015), as well as some nematodes (Ferri *et al*. 2011). Some clades of filarial nematodes require *Wolbachia* for normal development and fertility (Bandi *et al*. 1998, reviewed in Manoj *et al*. 2021); in contrast, variants associated with arthropods tend to be facultative and not required by hosts (Hoffmann and Cooper 2025). Most facultative *Wolbachia* show phylogenetic distributions consistent with horizontal host switching, with closely related variants observed in distantly related hosts (*e.g.,* O’Neill et al. 1992; Turelli *et al*. 2018; Vancaester and Blaxter 2023). Using clades defined by their affinity to the first two *Wolbachia* variants found in *Drosophila* species – *w*Mel in *D. melanogaster* (Hoffmann 1988) and *w*Ri in *D. simulans* (Hoffmann *et al*. 1986) – our goal is to describe the tempo and patterns of their distribution among diverse hosts and their genome evolution, focusing on pairs of “cytoplasmic incompatibility factor” (*cif*) genes.

Both *w*Ri and *w*Mel (and many *Wolbachia* in our “*w*Mel-like” and “*w*Ri-like” samples) cause cytoplasmic incompatibility (CI), the most common reproductive phenotype associated with *Wolbachia* (Shropshire *et al*. 2020a). Embryos from females lacking *Wolbachia* fertilized by sperm from males with *Wolbachia* suffer increased mortality, the defining feature of CI. The genes that cause CI (*cifB* and occasionally *cifA*) and rescue CI-induced lethality (*cifA*) are associated with prophage (*Wovirus*) regions of *Wolbachia* genomes (LePage *et al*. 2017; Beckmann *et al*. 2017; Shropshire *et al*. 2018, 2021a; Shropshire and Bordenstein 2019; Adams *et al*. 2021; Sun *et al*. 2022). *Woviruses* are diverse and show patterns consistent with horizontal movements among *Wolbachia* genomes (Masui *et al*. 2000; Bordenstein and Wernegreen 2004; Gavotte *et al*. 2007; Chafee *et al*. 2010; Bordenstein and Bordenstein 2022), analogous to *Wolbachia* host switching (O’Neill *et al*. 1992; Boyle *et al*. 1993; Werren *et al*. 1995; Heath *et al*. 1999; Van Meer *et al*. 1999; Dyson *et al*. 2002; Russell *et al*. 2009; Ahmed *et al*. 2016; Turelli *et al*. 2018; Martinez *et al*. 2021; Vancaester and Blaxter 2023). *cif* diversity also reflects horizontal movements (Lindsey *et al*. 2018; Shropshire *et al*. 2020; Martinez *et al*. 2021; Vancaester and Blaxter 2023), including patterns consistent with *cif* movements between *Woviruses* (*e.g.,* LePage *et al*. 2017; Cooper *et al*. 2019). Transposable elements are associated with *cif* genes within *Wovirus* regions (*e.g.,* Amoros *et al*. 2025a), plausibly mediating these exchanges (Gillespie *et al*. 2018; Cooper *et al*. 2019; Madhav *et al*. 2020). Hence, *Wolbachia* move among hosts, *Woviruses* move among *Wolbachia* genomes, and *cifs* move with and among *Woviruses*. The timescales of these movements are uncertain.

Understanding the timescale of *Wolbachia* acquisitions and genomic evolution requires accurate calibrations of *Wolbachia* divergence times. Turelli *et al*. (2018) estimated divergence times for *w*Ri-like *Wolbachia* using a mutation-based calibration of *Wolbachia* molecular evolution from Richardson *et al*. (2012). They concluded that variants which diverged only about 30,000 years ago occupy *Drosophila* hosts that diverged over 40 million years ago. However, mutation rates may overestimate the substitution rates that underlie chronograms if mutation rates do not accurately reflect substitution rates (Richardson *et al*. 2012). Fossil-based estimates of divergence times provide an alternative framework for estimating rates of genome evolution. When *Wolbachia* codiverge with their hosts, fossil-based host divergence times match divergence times for their concordantly diverging *Wolbachia*, allowing estimation of *Wolbachia* molecular evolution rates (*cf.* Raychoudhury *et al*. 2009; Gerth and Bleidorn 2016). However, *Wolbachia*-host codivergence is rarely documented by comparing molecular divergence of nuclear and mtDNA genomes of hosts with that of their *Wolbachia* (*cf.* Hamm *et al*. 2014; Conner *et al*. 2017).

Hence, it is often difficult to establish *Wolbachia*-host codivergence for facultative variants with uncertain evolutionary histories. In addition, chronograms, which are essential to conclusions, assume bifurcating evolution; yet prior analyses suggest that *Wolbachia* can have a mosaic genome structure produced by recombination and horizontal genomic exchanges between closely and distantly related variants (Werren and Bartos 2001; Jiggins *et al*. 2001; Baldo *et al*. 2006; Ellegaard *et al*. 2013; Wang *et al*. 2020). Prior attempts to resolve the timescales of *Wolbachia* movements (*e.g.,* Raychoudhury *et al*. 2009; Gerth and Bleidorn 2016; Turelli *et al*. 2018) have mostly ignored the potential for recombination to bias divergence-time estimates.

To produce robust calibrations, we first assess support for bifurcating evolution of regions of *Wolbachia* genomes. We provide evidence that over the timescales of divergence of the *w*Mel-like and *w*Ri-like clades, large portions of their genomes, specifically most single-copy protein-coding genes, evolve as bifurcating lineages, with little evidence of horizontal acquisition of alleles from distantly related *Wolbachia*. We address how recent bifurcation of these genomic regions can be reconciled with the evidence for recombination among *Wolbachia*. We use these bifurcating regions of genomes as a temporal standard to calibrate rates of *Wolbachia* molecular evolution using new data from trios of filarial nematodes (Qing *et al*. 2025) that carry obligate *Wolbachia* (Hoerauf *et al*. 2000; Taylor *et al*. 2005; Fenn and Blaxter 2004). Extensive evidence supports codivergence of filarial nematodes and their *Wolbachia* in particular clades (Moran 2007; Lefoulon *et al*. 2016; Vancaester *et al*. 2025). The case for codivergence is strengthened by data supporting obligate symbioses (see Manoj *et al*. 2021).

We provide new estimates of *w*Ri-like divergence and also estimate divergence times for closely related *w*Mel-like *Wolbachia* from widely divergent hosts, spanning holometabolous insects. Although the nematode *Wolbachia* lineages we use for calibration are distantly related to *w*Mel-like and *w*Ri-like *Wolbachia*, the molecular evolution rates they imply are consistent with some previous estimates derived from the facultative and plausibly codiverging *Wolbachia* in *Nasonia* wasps (Raychoudhury *et al*. 2009) and *Nomada* bees (Gerth and Bleidorn 2016) that are much more similar to *w*Mel and *w*Ri than the nematode *Wolbachia* lineages. With our calibrations, we also show that *Woviruses* and *cif* genes are rapidly gained and lost outside of the bifurcating *w*Ri-like and *w*Mel-like genomic regions. Analyzing selection on intact *cif* genes from *Wolbachia* from different host species, we find evidence that selection acts on both *cifA* and *cifB* sequence evolution. We discuss these results in light of faster *cifB* mutation accumulation (*e.g.,* Meany *et al*. 2019; Martinez *et al*. 2021), and theoretical predictions concerning how selection acts on CI phenotypes within host species (Prout 1994; Turelli 1994; Haygood and Turelli 2009) and in the proliferation of CI phenotypes among *Wolbachia* across host species (Hurst and McVean 1996; Turelli *et al*. 2022). Overall, our analyses provide insights into *Wolbachia* genome evolution and shed light on the timescale of host switching and the evolution of incompatibility loci for two closely related and model *Wolbachia* clades.

## Results

### *w*Mel-like *Wolbachia* and their hosts

Our study includes 19 closely related *Wolbachia* plus model pathogen-blocking *w*Mel found in *D. melanogaster.* Fig. 1A presents our cladogram for these variants, which we refer to as “*w*Mel-like” *Wolbachia* (Hague *et al*. 2020). Hosts that carry *w*Mel-like *Wolbachia* are diverse taxonomically and in their life histories (Tables 1 & S1; Supplemental Information). They include species from the clades Diptera and Hymenoptera, which diverged approximately 350 million years ago (MYA), and span the Holometabola (Wang *et al*. 2016; Fig. 1B; Supplemental Information). These patterns imply phylogenetic discordance between *w*Mel-like *Wolbachia* and their hosts, as observed between many *Wolbachia* and host species (*e.g.,* Fig. 2B in Vancaester and Blaxter 2023; O’Neill *et al*. 1992), including *w*Ri-like *Wolbachia* found in hosts that span the *Drosophila* genus (Turelli *et al*. 2018).

**Figure 1.**
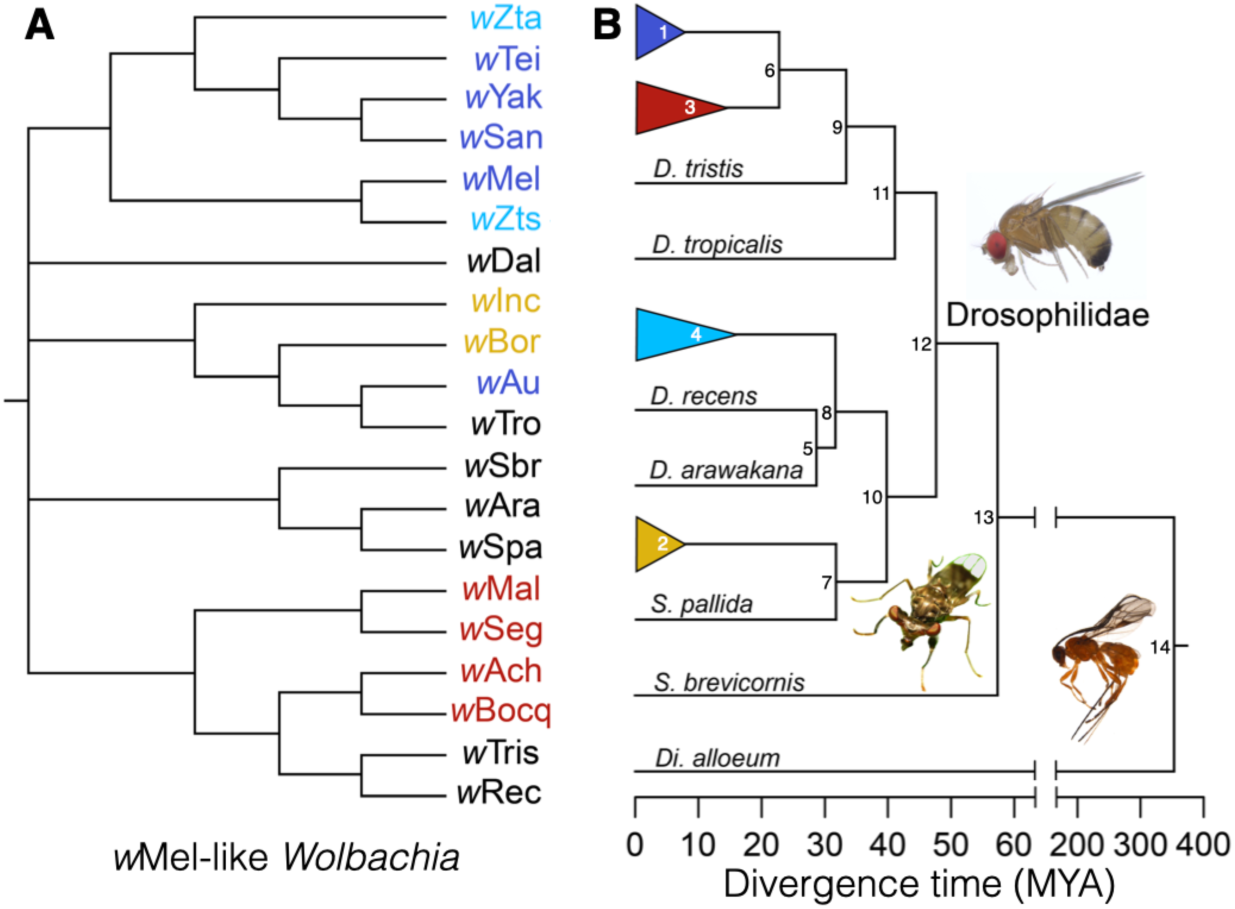
Closely related *w*Mel-like *Wolbachia* occupy divergent hosts. **(A)** A cladogram for the *w*Mel-like *Wolbachia* included in our study (Tables 1 & S1). Note that several of the basal nodes are not resolved, reflecting rapid diversification of these *w*Mel-like variants. The *Wolbachia* labels are colored to match their hosts in panel B. **(B)** An approximate chronogram for the hosts of the *w*Mel-like *Wolbachia* from panel A. Node numbers correspond to the node numbers in Table 2, which provides node ages and approximate credible intervals. For the drosophilids, the dark blue clade includes *melanogaster* subgroup hosts, the red clade includes *montium* subgroup hosts, the light blue clade includes two *Zaprionus* hosts, and the gold clade includes *D. borealis* and *D. incompta*. The species within focal clades and their relationships are also presented in Table 2, and additional information on host divergence is presented in the Supplemental Information. Open-source images taken by CBD (*Diachasma alloeum*), K. Schulz (*S. brevicornis*) and T. Wheeler (*D. teissieri*).

**Figure. 2.**
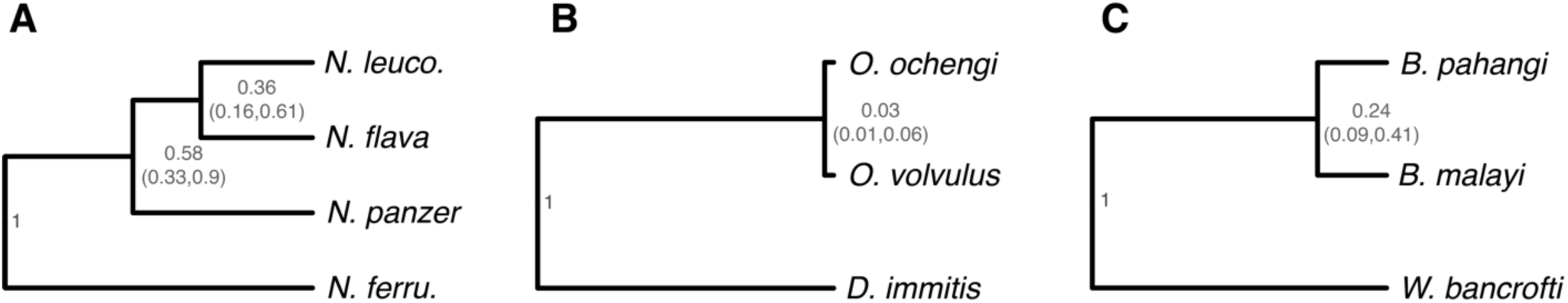
Relative chronogram for the four-species *Nomada* clade and estimates of divergence times of nematode congeners relative to their respective outgroup. (A) A relative chronogram for the four-species *Nomada* clade: (*N. ferruginata*, (*N. panzer*, (*N. flava*, *N. leucophthalma*))). Following Gerth and Bleidorn (2016, p. 4), the crown age of the clade is approximately 2.14 MYA. The internal nodes have relative ages 0.58 (with 95% HPD support interval 0.33–90) and 0.36 (with 95% HPD support interval 0.16–0.61). Thus, the MRCA ages (and support intervals) for (*N. panzer*, (*N. flava*, *N. leucophthalma*)) and (*N. flava*, *N. leucophthalma*) are roughly 1.24 (0.71–1.93) MYA and 0.77 (0.34–1.31) MYA. Estimates for the divergence times of (B) *O. volvulus* and *O. ochengi* and of (C) *B. malayi* and *B. pahangi*, each relative to their respective outgroup. The relative age of the MRCA for *O. volvulus* and *O. ochengi* is 0.031, with 95% HPD 0.012–0.059, corresponding to about 2.01 MYA; for *B. malayi* and *B. pahangi*, the relative divergence time is 0.23, with 95% HPD 0.09–0.41, corresponding to about 6.66 MYA.

**Table 1.** Our study includes 20 *w*Mel-like and the 8 *w*Ri-like *Wolbachia* (*N* = 37 *w*Ri-like genomic variants) described in Turelli *et al*. (2018). All variants are members of the larger *Wolbachia* supergroup A. We also include the supergroup A *Nomada* (*N* = 4) and the supergroups C (*N* = 3) and D (*N* = 4) nematode-associated *Wolbachia* variants used for alternative calibrations. We present taxonomic information for both *Wolbachia* and the hosts that carry them. In Table S1, we provide all genome accessions and metrics of *Wolbachia* genome quality.

| *Wolbachia* | Group | Host Order | Host Genus | Host Species |
| --- | --- | --- | --- | --- |
| wSan | wMel-like | Diptera | *Drosophila* | *santomea* |
| wYak | wMel-like | Diptera | *Drosophila* | *yakuba* |
| wTeis | wMel-like | Diptera | *Drosophila* | *teissieri* |
| wZta | wMel-like | Diptera | *Zaprionus* | *taronus* |
| wZts | wMel-like | Diptera | *Zaprionus* | *tsacasi* |
| wMel | wMel-like | Diptera | *Drosophila* | *melanogaster* |
| wAu | wMel-like | Diptera | *Drosophila* | *simulans* |
| wTro | wMel-like | Diptera | *Drosophila* | *tropicalis* |
| wBor | wMel-like | Diptera | *Drosophila* | *borealis* |
| wInc | wMel-like | Diptera | *Drosophila* | *incompta* |
| wAch | wMel-like | Diptera | *Drosophila* | *sp. aff. chauvacae* |
| wBocq | wMel-like | Diptera | *Drosophila* | *bocqueti* |
| wTris | wMel-like | Diptera | *Drosophila* | *tristis* |
| wRec | wMel-like | Diptera | *Drosophila* | *recens* |
| wMal | wMel-like | Diptera | *Drosophila* | *malagassya* |
| wSeg | wMel-like | Diptera | *Drosophila* | *seguyi* |
| wSpa | wMel-like | Diptera | *Scaptomyza* | *pallida* |
| wAra | wMel-like | Diptera | *Drosophila* | *arawakana* |
| wSbr | wMel-like | Diptera | *Sphyracephala* | *brevicornis* |
| wDal | wMel-like | Hymenoptera | *Diachasma* | *alloeum* |
| wRi | wRi-like | Diptera | *Drosophila* | *simulans* |
| wAno | wRi-like | Diptera | *Drosophila* | *anomalata* |
| wPan | wRi-like | Diptera | *Drosophila* | *pandora* |
| wAna | wRi-like | Diptera | *Drosophila* | *ananassae* |
| wSuz | wRi-like | Diptera | *Drosophila* | *suzukii* |
| wSpc | wRi-like | Diptera | *Drosophila* | *subpulchrella* |
| wAur | wRi-like | Diptera | *Drosophila* | *auraria* |
| wTri | wRi-like | Diptera | *Drosophila* | *triauraria* |
| wNfla | Nomada | Hymenoptera | *Nomada* | *flava* |
| wNfe | Nomada | Hymenoptera | *Nomada* | *ferruginata* |
| wNleu | Nomada | Hymenoptera | *Nomada* | *leucophthalma* |
| wNpa | Nomada | Hymenoptera | *Nomada* | *panzeri* |
| wOo | supergroup C | Rhabditida | *Onchocerca* | *ochengi* |
| wOv | supergroup C | Rhabditida | *Onchocerca* | *volvulus* |
| wDimm | supergroup C | Rhabditida | *Dirofilaria* | *immitis* |
| wBm | supergroup D | Rhabditida | *Brugia* | *malayi* |
| wBp | supergroup D | Rhabditida | *Brugia* | *pahangi* |
| wWb | supergroup D | Rhabditida | *Wuchereria* | *bancrofti* |
| wLs | supergroup D | Rhabditida | *Litomosoides* | *sigmodontis* |

**Table 2.**
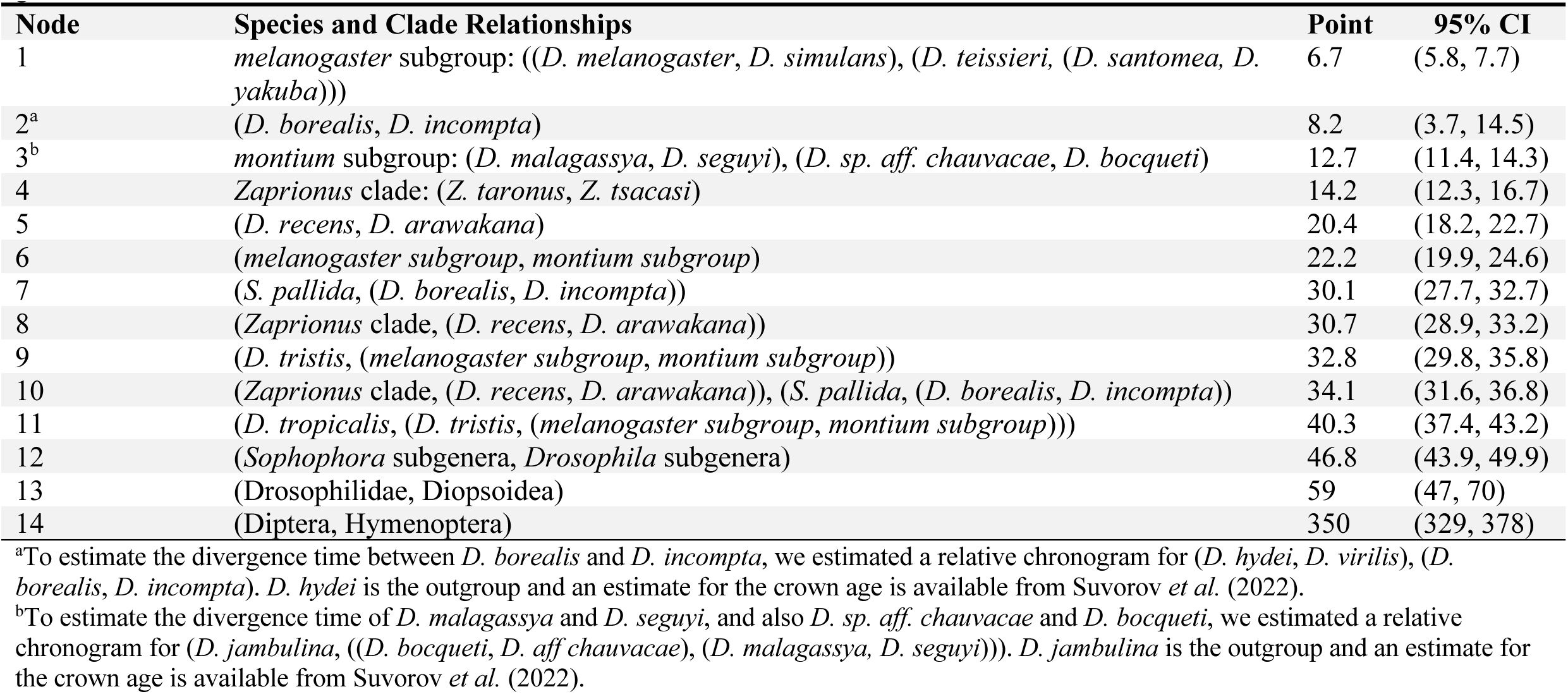
Ages and approximate confidence intervals in millions of years for hosts that carry *w*Mel-like *Wolbachia*. Node numbers correspond to those presented in Fig. 1B and are presented in order from the youngest to the oldest nodes. Table S1 provides the genome accessions.

| Node | Species and Clade Relationships | Point | 95% CI |
| --- | --- | --- | --- |
| 1 | *melanogaster* subgroup: ((*D. melanogaster*, *D. simulans*), (*D. teissieri*, (*D. santomea*, *D. yakuba*))) | 6.7 | (5.8, 7.7) |
| 2<sup>a</sup> | (*D. borealis*, *D. incompta*) | 8.2 | (3.7, 14.5) |
| 3<sup>b</sup> | *montium* subgroup: (*D. malagassya*, *D. seguyi*), (*D. sp. aff. chauvacae*, *D. bocqueti*) | 12.7 | (11.4, 14.3) |
| 4 | *Zaprionus* clade: (*Z. taronus*, *Z. tsacasi*) | 14.2 | (12.3, 16.7) |
| 5 | (*D. recens*, *D. arawakana*) | 20.4 | (18.2, 22.7) |
| 6 | (*melanogaster* subgroup, *montium* subgroup) | 22.2 | (19.9, 24.6) |
| 7 | (*S. pallida*, (*D. borealis*, *D. incompta*)) | 30.1 | (27.7, 32.7) |
| 8 | (*Zaprionus* clade, (*D. recens*, *D. arawakana*)) | 30.7 | (28.9, 33.2) |
| 9 | (*D. tristis*, (*melanogaster* subgroup, *montium* subgroup)) | 32.8 | (29.8, 35.8) |
| 10 | (*Zaprionus* clade, (*D. recens*, *D. arawakana*)), (*S. pallida*, (*D. borealis*, *D. incompta*)) | 34.1 | (31.6, 36.8) |
| 11 | (*D. tropicalis*, (*D. tristis*, (*melanogaster* subgroup, *montium* subgroup))) | 40.3 | (37.4, 43.2) |
| 12 | (*Sophophora* subgenera, *Drosophila* subgenera) | 46.8 | (43.9, 49.9) |
| 13 | (*Drosophilidae*, *Diopsoidea*) | 59 | (47, 70) |
| 14 | (*Diptera*, *Hymenoptera*) | 350 | (329, 378) |

<sup>a</sup>To estimate the divergence time between *D. borealis* and *D. incompta*, we estimated a relative chronogram for (*D. hydei*, *D. virilis*), (*D. borealis*, *D. incompta*). *D. hydei* is the outgroup and an estimate for the crown age is available from Suvorov *et al.* (2022).
<sup>b</sup>To estimate the divergence time of *D. malagassya* and *D. seguyi*, and also *D. sp. aff. chauvacae* and *D. bocqueti*, we estimated a relative chronogram for (*D. jambulina*, ((*D. bocqueti*, *D. aff. chauvacae*), (*D. malagassya*, *D. seguyi*))). *D. jambulina* is the outgroup and an estimate for the crown age is available from Suvorov *et al.* (2022).

After introducing host chronograms for *Wolbachia* calibrations and establishing that major portions of *Wolbachia* genomes show patterns consistent with a bifurcating evolutionary history (*i.e.*, a history of vertical descent), we use new data on filarial nematodes with vertically inherited *Wolbachia* to resolve the timescale of *w*Mel-like *Wolbachia* host switching and to revise our prior *w*Ri-like estimates. We then describe the tempo and patterns of *w*Mel-like and *w*Ri-like *Wolbachia* genome evolution outside of bifurcating regions, focusing on pairs of *cif* genes involved in protecting from and inducing CI. Our study does not include all known variants, *e.g.,* variants closely related to *w*Au (Miller and Riegler 2006), but our conclusions are robust to incomplete sampling of these clades as we discuss below.

### Host chronograms for calibrating *Wolbachia* divergence

*Wolbachia* chronograms rely on molecular clock approximations that can be obtained when *Wolbachia* codiverge with their hosts and fossil-based estimates of host divergence times are available. We first consider host chronograms for three cases in which *Wolbachia*-host codivergence has been proposed: the four-species *Nomada* bee clade with facultative *Wolbachia* analyzed by Gerth and Bleidorn (2016) and two trios of filarial nematodes with obligate *Wolbachia*, for which both *Wolbachia* and host genomic data have recently become available.

Fig. 2A presents our relative chronogram for the four-species *Nomada* clade: (*N. ferruginata*, (*N. panzeri*, (*N. flava*, *N. leucophthalma*))). Following Gerth and Bleidorn (2016, p. 4), the crown age of the clade is approximately 2.14 MYA. The internal nodes have relative ages 0.58 (with 95% HPD support interval 0.33–90) and 0.36 (with 95% HPD support interval 0.16–0.61). Thus, the MRCA ages (and support intervals) for (*N. panzeri*, (*N. flava*, *N. leucophthalma*)) and (*N. flava*, *N. leucophthalma*) are roughly 1.24 (0.71–1.93) MYA and 0.77 (0.34–1.31) MYA. These estimates of internal node ages are consistent with those that Gerth *et al*. (2013, Fig. 3) obtained from more limited host nuclear data. In contrast, when Gerth and Bleidorn (2016, Fig. 3) estimated these node ages using *Wolbachia* genomes, assuming codivergence with all four hosts, their point estimates for these branch splits, 0.43 MYA and 0.25 MYA, respectively, fall outside the support intervals obtained from our more extensive nuclear-gene analyses. We argue that these comparative genomic data, including *Wolbachia* and host nuclear and mtDNA, suggest horizontal transmission of *Wolbachia* within the ingroup clade (*N. panzeri*, (*N. flava*, *N. leucophthalma*)) explain the discordant chronograms based on host versus *Wolbachia* genomes (Supplemental Information).

**Fig. 3.**
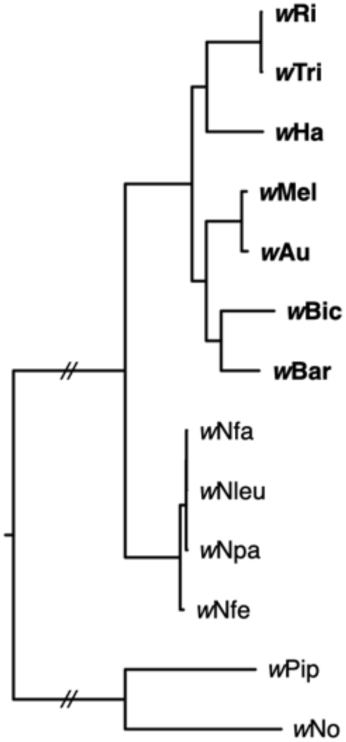
Phylogram of seven focal supergroup A *Wolbachia*, four supergroup A *Nomada*-derived *Wolbachia*, and *w*No and *w*Pip from supergroup B. Phylogram based on 438 concatenated loci. The *Wolbachia* include four supergroup A variants from *Nomada* species, (*w*Nfla, *w*Nleu, *w*Nfe, and *w*Npa), and *w*No and *w*Pip from supergroup B. These six *Wolbachia* were used to root the seven focal supergroup A genomes (bold) that span the divergence of the *w*Mel-like and *w*Ri-like clades. The resulting “consensus phylogeny,” denoted C, for the seven supergroup A variants is: (((*w*Mel, *w*Au), (*w*Bic, *w*Bar)), ((*w*Ri, *w*Tri), *w*Ha)). This phylogeny has posterior probabilities of 1 for all nodes and 100% bootstrap support over 1000 replicates. Branches leading to supergroups A and B are shortened for presentation (//).

We also considered two trios of filarial nematodes, each with a pair of congeners and an outgroup containing a *Wolbachia* variant belonging to the same supergroup as those in the congeners: (*Dirofilaria immitis*, (*Onchocerca volvulus, O. ochengi*)) that contain supergroup C *Wolbachia*, and (*Wuchereria bancrofti*, (*Brugia malayi, B. pahangi*)) that contain supergroup D *Wolbachia*. Unlike prior calibrations from *Nasonia* and *Nomada* that carry facultative *Wolbachia*, these hosts carry obligate and vertically inherited *Wolbachia* known to codiverge with their hosts. For both trios, Qing *et al*. (2025) estimated the time to the MRCA of the three species. The estimate for *Dirofilaria* and *Onchocerca* is 64.73 MYA with 95% HPD 37.56–91.96 MYA; the estimate for *Wuchereria* and *Brugia* is 28.97 MYA, with 95% HPD 12.8–48.59 MYA. Figures 2B and 2C provide estimates for the divergence times of the congeners relative to their respective outgroup. The relative age of the MRCA for *O. volvulus* and *O. ochengi* is 0.031, with 95% HPD 0.012–0.059, corresponding to about 2.01 MYA; for *B. malayi* and *B. pahangi*, the relative divergence time is 0.23, with 95% HPD 0.09–0.41, corresponding to about 6.66 MYA.

### Assessing support for bifurcating evolution of major portions of *Wolbachia* genomes across two orders of magnitude of genomic divergence

Robust calibrations rely on bifurcating phylogenies, which assume that differences among tip sequences arose from independent evolutionary changes along the branches, without introgressions from genomes with distinct evolutionary histories. Recombination between *Wolbachia* genomes has been repeatedly demonstrated (Werren and Bartos 2001; Jiggins *et al*. 2001; Baldo *et al*. 2006; Ellegaard *et al*. 2013), so it is important to assess evidence for recombination and the extent to which horizontal acquisition of sequence elements may preclude meaningful analyses of divergence times. Prior attempts to resolve the timescales of *Wolbachia* movements (including our own) have mostly ignored the potential for recombination to bias divergence-time estimates.

To understand the phylogenetic implications of recombination, it is critical to recognize that recombination events in bacteria are asymmetric and heterogeneous. For *Wolbachia*, “recombination” can result in: (1) replacing a genome fragment in a recipient genome with an homologous fragment from a donor genome, (2) integrating donor genome elements, such as transposable elements (TEs) or prophages, into recipient positions that need not be homologous to positions in the donor genome, or (3) recombination-mediated rearrangements within a single genome, such as inversions, phage or TE movements, or duplications. The recombination studies cited above found portions of individual loci with distinct phylogenies, indicating horizontal acquisition of fragments from relatively distantly related donor genomes. The resulting mosaic structure is clearly inconsistent with a single bifurcating history of the recombinant locus. Similarly, horizontal acquisition of larger fragments, involving complete genes with alleles that are distantly related to the alleles replaced, are inconsistent with bifurcation. In contrast, exchange of genome fragments between essentially identical *w*Mel-like or *w*Ri-like genomes that involve few or no SNPs would have little or no effect on recreating an essentially bifurcating history. Because it is well known that TEs and prophage move within genomes and among *Wolbachia* genomes (Masui *et al*. 2000; Bordenstein and Wernegreen 2004; Riegler *et al*. 2005; Cerveau *et al*. 2015) and that rearrangements, such as inversions, are common (Wu *et al*. 2004; Klasson *et al*. 2009; Cerveau et al. 2011; Ellegaard et al. 2013; Baião *et al*. 2021), we focus on determining whether closely related genomes, such as our *w*Ri-like and *w*Mel-like samples, exhibit a bifurcating history for single-copy, protein-coding *Wolbachia* loci (*i.e.*, loci not within TEs or prophages). Intragenome rearrangements, such as inversions, do not alter the bifurcating history of individual genes.

To better understand the dynamics of *Wolbachia* genome evolution in our system, we assessed evidence for bifurcating evolution in sets of variants with increasing levels of sequence divergence: (1) a set of 8 *w*Ri-like *Wolbachia* (Turelli *et al*. 2018), (2) the set of 20 *w*Mel-like *Wolbachia* introduced here, and (3) a broader set of seven supergroup A variants. These seven include two *w*Mel-like variants (*w*Mel and *w*Au), two *w*Ri-like variants (*w*Ri and *w*Tri), *w*Ha from *D. simulans*, *w*Bic from *D. bicornuta*, and *w*Bar from *D. barbarae.* As noted in Fig. 3, the clade defined by these seven spans our *w*Ri-like and *w*Mel-like clades. In every case, our analyses included single-copy genes extracted from each set, using criteria described in our Materials and Methods. We focused on these genes because they are used in our downstream calibrations. In total, we extracted 525 genes (506,307 bp) from *w*Ri-like genomes, 438 genes (409,848 bp) from *w*Mel-like genomes, and 442 genes (413,115 bp) from the 7 supergroup A variants. We also assessed whether the nematode *Wolbachia* genomes evolved primarily through bifurcation. In the following subsections, we describe the results of several tests demonstrating that these single-copy genes comprise bifurcating genomic regions, from which we can robustly estimate *Wolbachia* divergence times.

### Analyzing intragenic recombination

We first assessed evidence for intralocus recombination using a genetic algorithm for recombination detection (GARD) (Kosakovsky Pond *et al*. 2006), plus all three statistical methods implemented in PhiPack (Bruen *et al*. 2006). We found no evidence for intragenic recombination in our least-diverged set of *w*Ri-like *Wolbachia*. It is worth noting that given the extreme similarity of the *w*Ri-like genomes, recombination between variants within this clade would be essentially impossible to detect, and our ability to detect recombination increases with the divergence of our sets of variants. However, we found almost no evidence for intragenic recombination in our set of slightly more diverged *w*Mel-like *Wolbachia* (GARD: 0/411 genes; PHI test: 0/411, Max χ2 test: 1/411, Neighbor Similarity Score: 0/411, FDR-adjusted *P* < 0.01), and in our most diverged set that includes four samples of supergroup D *Wolbachia*: GARD: 0/356 genes; PHI test: 0/356, Max χ2 test: 1/356, Neighbor Similarity Score: 0/356, FDR-adjusted *P* < 0.01). Analysis of supergroup C *Wolbachia* was not possible due to an insufficient number of available sequences. In contrast, we found evidence of intragenic recombination within many of the genes that met our criteria for the intermediately diverged set of seven supergroup A variants (GARD: 187/401 genes; PHI test: 40/401, Max χ2 test: 110/401, Neighbor Similarity Score: 63/401, FDR-adjusted *P* < 0.01). In the Supplemental Information, we describe that when sufficient variation exists for GARD to produce a tree, its recombination calls are error prone if there is insufficient variation to accurately resolve trees for putative partitions. Thus, we completed several alternative tests that illustrate the bifurcating evolution of major portions of *Wolbachia* genomes across levels of divergence relevant to our analyses of *w*Mel-like and *w*Ri-like variants.

### Analyzing of single-gene phylogenies

To determine the prevalence of horizontal exchanges and departures from purely bifurcating evolution, we next assessed single-gene tree discordance. We estimated single-gene phylogenies from 442 individual genes in the group of seven supergroup A *Wolbachia*, of which 52 genes produced fully resolved topologies with posterior probabilities > 0.95 at each node. Of the 52 phylogenetically informative loci, all supported the pairs (*w*Mel, *w*Au) and (*w*Ri, *w*Tri) as closest relatives. Using IQ-TREE (Wong *et al*. 2025), we observed gene concordance factors (gCFs) > 0.99 for both (*w*Au, *w*Mel) and (*w*Ri, *w*Tri) across bootstrap thresholds (50%, 60%, 80% and 95%) used to collapse gene-tree nodes prior to scoring decisiveness (Table S2). These results suggest that essentially no horizontal transmission disrupts the bifurcating evolution of these genomic regions within the *w*Mel-like and *w*Ri-like clades. This is supported by our polymorphism analyses in the following subsection, which assessed the 52 loci described above and those in the unresolved class.

Departures from pervasive bifurcation seem more plausible when we consider all seven supergroup A genomes that span the *w*Mel-like and *w*Ri-like clades. We estimated a consensus phylogeny by concatenating all 442 loci, using the four *Nomada*-derived *Wolbachia* as outgroups from supergroup A and *w*No and *w*Pip from supergroup B to root it. This “consensus phylogeny,” denoted C, is: (((*w*Mel, *w*Au), (*w*Bic, *w*Bar)), ((*w*Ri, *w*Tri), *w*Ha)), with posterior probabilities of 1 for all nodes and 100% bootstrap support over 1000 replicates (Fig. 3). We also estimated an unrooted species tree from our set of 442 gene trees using weighted ASTRAL (Zhang and Mirarab 2022). The species tree agreed with C, with local posterior support of 1 for all internal branches. ASTRAL quartet frequencies were skewed toward the reference resolution (median q1 = 0.70, range: 0.44–0.97; Table S3), consistent with short but positive internode lengths under the multi-species coalescent. While gCFs were over 0.99 for (*w*Au, *w*Mel) and (*w*Ri, *w*Tri), they were ∼0.56 for the other nodes that were again relatively insensitive to the collapse threshold (see below; Table S2). It is notable that the stem branches leading to the three clades (*w*Bic, *w*Bar), ((*w*Mel, *w*Au), (*w*Bic, *w*Bar)), and ((*w*Ri, *w*Tri), *w*Ha) are significantly shorter than the terminal branches leading to *w*Ha, *w*Bic and *w*Bar and the internal branches leading to the *w*Mel-like and *w*Ri-like pairs. Table S4 indicates the number of loci that support each of the 15 possible unrooted trees of the five entities *w*Ha, *w*Bic, *w*Bar, (*w*Mel, *w*Au) and (*w*Ri, *w*Tri). Each topology was supported by at least one of the 52 phylogenetically informative loci, with 14 loci supporting the consensus phylogeny. Each of the alternative unrooted topologies, denoted A1–A14 in Table S4, are supported by 6 or fewer loci.

The loci supporting the alternative topologies, A1–A14, are candidates for belonging to horizontally acquired genomic fragments. However, alternative topologies may also arise if multiple *Wolbachia* genomes that differ slightly are horizontally transmitted between host species. This could produce discordant gene trees by a process analogous to incomplete lineage sorting (Gillespie and Langley 1979; Maddison 1997; Edwards 2009). Indeed, the resolved ASTRAL species tree and the observed gCF values together point to short-branch ILS or inference errors associated with model mis-specification as plausible sources of discordance, rather than pervasive horizontal exchange. In contrast, if the “discordant” loci derive from recombination events involving integration of genomic fragments acquired via transformation or transduction from distantly related *Wolbachia* genomes, the loci might be expected to show atypically high levels of pairwise differences relative to loci that did not experience horizontal transfer (*cf.* Wang *et al*. 2020). In general, the phylogenetically informative loci might show relatively high levels of sequence divergence, given that we selected them based on their ability to produce a fully resolved tree. Moreover, if horizontal transfer involves large genomic segments, we might expect the discordant loci to be clustered. Table S5 shows the positions of the 52 phylogenetically informative loci with respect to 4 reference genomes, *w*Mel, *w*Au, *w*Ri and *w*Ha. The genes are listed in their order of occurrence relative to the origin of the *w*Mel reference genome. Under each alternative reference genome, we present in parentheses their order of appearance in that reference genome. The gene-order differences illustrate the pervasiveness of rearrangements, especially inversions, as documented by Ellegaard *et al*. (2013). The last two columns give two measures of gene-specific divergence among the seven genomes: the fraction of polymorphic sites within each gene and the total number of substitutions across the entire tree (estimated from the single-gene phylogram).

As expected, the loci that supported non-consensus topologies versus the consensus topology have more polymorphic sites (means ± standard deviations: 0.051 ± 0.020 versus 0.040 ± 0.026) and higher estimated numbers of substitutions (0.062 ± 0.026 versus 0.050 ± 0.022), but the differences are neither large nor statistically significant (*t*-tests, *P* = 0.085 and 0.105, respectively). This suggests that these loci supporting non-consensus topologies did not arise by horizontal transmission from deeply divergent *Wolbachia*. Our allele-divergence analyses in the next subsection further suggest relatively few examples of horizontally acquired alleles among our filtered single-copy transcribed loci that comprise bifurcating genomic regions.

From the seven *Wolbachia* from filarial nematodes, we extracted 143 single-copy genes (133,677 bp). Of these, 83 produced fully resolved single gene trees with posterior probabilities >0.95 at each node. All 83 genes produced the same phylogeny, which recapitulated the host phylogeny: ((((*B. malayi, B. pahangi*), *W. bancrofti*), *L. sigmodontis*), ((*O. volvulus, O. ochengi)*, *D. immitis*)). Like our recombination analyses above, this analysis supported bifurcating evolution of filarial nematode *Wolbachia* over tens of millions of years.

### Analyzing polymorphism and allelic differences at individual loci

If horizontal acquisition of single-copy genes from distantly related *Wolbachia* occurs frequently (*e.g.,* on a timescale of hundreds or a few thousand years), outlier alleles should be most apparent in sets of closely related genomes. To assess the timing and pervasiveness of horizontal gene transfer among *Wolbachia* genomes, we examined in turn genome sets showing three levels of differentiation. First, we considered representatives of the *w*Ri-like *Wolbachia* from eight *Drosophila* hosts (Turelli *et al*. 2018). For the *w*Ri-like set, the 525 single-copy genes showed a maximum observed pairwise third-position divergence (estimated number of substitutions per site) of 2.4 × 10^-4^ (*w*Ana versus *w*Tri) after Jukes-Cantor correction. Over the 438 single-copy genes, the *w*Mel-like variants differed by at most 2.1 × 10^-3^ (*w*Rec versus *w*Zts). Finally, over 442 single-copy genes, the seven supergroup A genomes showed a maximum third-position divergence of 3.1 x 10^-2^ (*w*Bic versus *w*Tri). Hence, our three sets span two orders of magnitude in their differentiation.

Based on these pairwise differences, we applied a heuristic criterion for identifying plausible examples of horizontally acquired alleles. For individual loci within a specific set of *Wolbachia*, we categorized an allele as a “two-fold outlier” if it differed from all other alleles in the set at twice as many nucleotides as any of the remaining alleles differed from one another. This heuristic criterion is obviously most easily satisfied for the least diverged *w*Ri-like clade. However, across the 525 single-copy gene set, only one relatively long single-copy gene (1,915 bp) showed four variable nucleotide sites; the remainder showed three or fewer. Thus, there were no two-fold outlier loci. As noted above, recombination between *w*Ri-like variants would have been essentially impossible to detect; but even if it occurred, it would not have significantly distorted the bifurcating divergence of genomic regions composed of single-copy loci on which we based our chronograms (see the following Results sections). However, our criteria for selecting loci to include in these analyses, namely that they be single-copy and equal length for at least *n* – 1 of the *n* alleles compared, may have screened out some loci with evidence of horizontally acquired alleles.

To identify outlier alleles in the *w*Mel-like and seven supergroup A gene sets, we ranked loci in each set by the fraction of variable nucleotides, under the assumption that the most variable loci are the most promising candidates for horizontal transmission. Among the 20 *w*Mel-like loci with the largest fraction of variable nucleotides, only 4 showed a two-fold outlier allele (Table S6), and only the second most-variable gene initially appeared as a plausible candidate for horizontal acquisition of distantly related alleles (Supplemental Information). This gene is homologous to RefSeq_WP_010962738.1, peptidase M48, with 43 variable sites across the 1,282 bp (3.35%). The *w*Bor allele differs from the other 19 alleles at 34–36 sites (2.7–2.8%), while the other alleles differ from one another at fewer than 5 sites (≤0.39%). The *w*Bor allele appears intact and ends with an in-frame stop codon. A 55 bp segment at the 3′ end of the *w*Bor allele matches a *w*Mel ISWpi1 insertion sequence element (EU714520.1), with 100% identity on the reverse strand. There are many ISWpi1 elements in the *w*Bor genome, and one outside the peptidase M48 (on a different contig) is an exact match of the one within peptidase M48. All other 19 alleles are identical across this 55 bp segment, each differs from the *w*Bor allele at 34 sites. At the 5’ of this gene in *w*Bor, the next single-copy gene is homologous to C4-dihydroxy-2-butanone-4-phosphate synthase, which shows less than 1% divergence between *w*Bor and *w*Mel. On the 3’ end, there are two transposases followed by a membrane protein in *w*Bor, while the transposases are absent and the membrane protein is present in *w*Mel.

BLAST found that the *w*Bor allele shares 99.9% similarity with an allele in *w*Inn from *D. innubila*. Both 5’ and 3’ flanking regions of the allele in *w*Inn match those in *w*Bor, including the two transposases at the 5’ end. The transposases closer to peptidase M48 in *w*Bor and *w*Inn are identical. Taken together these data are consistent with IS activity within the genome of the *w*Inn-*w*Bor MRCA generating the 55 bp fragment that distinguishes them from the 19 other alleles in our analysis. While transposon insertions from the same genome affect terminal branch lengths in a phylogram, they do not affect topology in the case of our 20 alleles at this gene, given that the 19 other alleles are essentially identical.

In contrast to the *w*Mel-like gene set, we did find limited evidence for horizontal transfer from a distantly related *Wolbachia* in the seven supergroup A set, with the most variable locus meeting our two-fold outlier criteria: the gene homologous to RefSeq_WP_010962975.1, a membrane protein. This locus showed about 19% polymorphic sites, with 95 variable sites out of 498 bp. The *w*Ha allele shows 92–94 bp differences from the homologous alleles in the other 6 *Wolbachia* (18.5–19.9%), whereas the other 6 alleles have a maximum of 3 pairwise differences (≤ 0.6%). BLAST revealed that the *w*Ha copy is identical to copies in *Wolbachia* from diverse host species, including sawflies, hoverflies, moths, carpet beetles, and Tachinidae flies (*e.g.,* Accessions: OZ034713.1, OX366332.1, AP028948.1, OX366359.1, OZ034764.1). The genes on either side did not seem exceptional. tRNA-Leu is on the 5’ side and identical in all seven genomes. On the 3’ side is uroporphyrinogen III methyltransferase (708 bp), where the *w*Ha allele has up to 27 pairwise differences over 708 bp from the other six, but the other alleles differed from one another at up to 18 bp (less than two-fold). No other locus met our two-fold outlier criteria. As noted previously, the 38 loci that supported non-consensus phylogenies did not show significantly more pairwise differences among the alleles than did the loci that supported the consensus phylogeny.

Our gene sets contained most of the single-copy genes in these genomes. In the 1.45 Mb *w*Ri reference genome, Prokka found 1,397 genes, of which 726 had a single copy. Removing what are clearly phage genes and transposases, the number of single-copy genes dropped to 694. Hence, ignoring our alignment and gap criteria, 694 genes (706,146 bp) set the theoretical maximum. The 525 genes we analyzed for “outliers” represent 75% of this theoretical maximum and roughly a third of the entire *w*Ri genome. In the 1.27 Mb *w*Mel reference genome, Prokka found 1,271 genes, of which 710 were single-copy genes and not phage genes or transposases. The *w*Mel-like and seven supergroup A gene sets we analyzed included more than half of these and represented about a third of the entire *w*Mel genome.

It appears that a majority of single-copy loci in our analyses did not acquire exogenous alleles over the timescales of differentiation for our *w*Ri-like and *w*Mel-like variants. Hence, these genomes contain substantial genomic regions composed of single-copy loci that undergo bifurcating evolution on which we can estimate chronograms, once we establish calibrations of *Wolbachia* evolution. In our Supplemental Information, we provide additional details and support for bifurcation.

### *Wolbachia* substitution-rate calibrations

While multiple approaches for calibrating *Wolbachia* divergence times have been developed, they may be unreliable for different reasons. For example, the earliest calibration probably underestimated divergence times because of relatively slow *ftsZ* evolution (Werren *et al*. 1995; Table S7; Supplemental Information), calibrations based on facultative associations may be unreliable if they do not actually involve codivergence (Raychoudhury *et al*. 2009), and mutation-based calibrations may be biased if mutation rates do not accurately reflect substitution rates (Richardson *et al*. 2012). We calibrated rates of *Wolbachia* molecular evolution using new data from trios of filarial nematodes (Qing *et al*. 2025) that are known to codiverge with their obligate *Wolbachia* that they require for survival (Comandatore *et al*. 2013; Manoj *et al*. 2021). Table 4 provides four estimates for the relative rates of divergence for the *Wolbachia* versus the host nuclear and mitochondrial genomes, derived from two trios: ((*O. volvulus, O. ochengi)*, *D. immitis*) and ((*B. malayi, B. pahangi*), *W. bancrofti*). Notably, the relative rates of *Wolbachia* and host nuclear divergence are quite consistent within each trio. However, the ratios for *Wolbachia*/nuclear divergence in the trio ((*B. malayi, B. pahangi*), *W. bancrofti*) is only about half that observed in ((*O. volvulus, O. ochengi)*, *D. immitis*). This underlies the differences in estimated *Wolbachia* substitution rates derived from these alternative trios.

**Table 3.** Absolute and relative divergence of *Wolbachia* versus host nuclear and mitochondrial (mtDNA) genomes for cases of putative cladogenic *Wolbachia* transmission. For each entry in the body of the table, the first value is the estimated percent substitutions (after Jukes-Cantor correction) per third-position codon site per year, the second value is the estimated percentage of synonymous substitutions across the first and third positions. The values in parentheses are total number of nucleotides in the exons used in the divergence estimates.

| Host Species | *Wolbachia* | Nuclear | mtDNA | *Wolbachia*/nuclear | mtDNA/*Wolbachia* | mtDNA/nuclear |
| --- | --- | --- | --- | --- | --- | --- |
| *Nasonia longicornis*(wNlonB1) / *N. giraulti*(wNgirB)<sup>1</sup> | 0.2, 0.3<br>(4486 bp) | NA, 1.12<br>(4135 bp) | NA, 45.24<br>(2241 bp) | NA, 0.27 | NA, 150.8 | NA, 40.4 |
| *Onchocerca ochengi*/*O. volvulus* | 0.55, 0.84<br>(379047 bp)<sup>2,3</sup> | 0.53, 0.77<br>(33099 bp)<sup>4</sup> | 7.60, 11.23<br>(10373 bp)<sup>4</sup> | 1.04, 1.09 | 13.8, 13.4 | 14.3, 14.6 |
| *Dirofilaria immitis*<sup>5</sup>/*O. ochengi* | 23.53, 35.27<br>(379047 bp)<sup>3</sup> | 21.93, 33.42<br>(33099 bp)<sup>4,5</sup> |  | 1.07, 1.06 |  |  |
| *D. immitis*/*O. volvulus* | 23.56, 35.33<br>(379047 bp)<sup>5</sup> | 22.02, 33.59<br>(33099 bp)<sup>4,5</sup> |  | 1.07, 1.05 |  |  |
| *Brugia malayi*<sup>2</sup>/*B. pahangi*<sup>3</sup> | 0.88, 1.24<br>(499230 bp)<sup>8</sup> | 1.94, 2.87<br>(33099 bp)<sup>6,7</sup> | 28.2, 43.7<br>(10361 bp)<sup>6,7</sup> | 0.45, 0.43 | 32.0, 35.2 | 14.5, 15.2 |
| *Wuchereria bancrofti*<sup>9</sup>/*B. malayi* | 4.30, 6.10<br>(499230 bp)<sup>8</sup> | 7.94, 11.66<br>(33099 bp) |  | 0.54, 0.54 |  |  |
| *W. bancrofti*/*B. pahangi* | 4.30, 6.10<br>(499230 bp)<sup>8</sup> | 7.93, 11.48<br>(33099 bp) |  | 0.54, 0.53 |  |  |
| ***Nomada* clade: (*N. ferruginata*, (*N. panzeri*, (*N. flava*, *N. leucophthalma*)))<sup>10</sup>** |  |  |  |  |  |  |
| *N. ferruginata*/*N. panzeri* | 0.29, 0.46<br>(613605 bp) | 1.73, 2.39<br>(36402 bp) | 2.53, 4.95<br>(9249 bp) | 0.17, 0.19 | 8.72, 10.76 | 1.46, 2.07 |
| *N. ferruginata*/*N. flava* | 0.27, 0.43<br>(613605 bp) | 1.43, 2.15<br>(36402 bp) | 2.61, 5.27<br>(9249 bp) | 0.19, 0.20 | 9.67, 12.26 | 1.83, 2.45 |
| *N. ferruginata*/*N. leucophthalma* | 0.27, 0.43<br>(613605 bp) | 1.47, 2.13<br>(36402 bp) | 2.22, 4.45<br>(9249 bp) | 0.18, 0.20 | 8.22, 10.35 | 1.51, 2.09 |
| *N. panzeri*/*N. flava* | 0.032, 0.051<br>(613605 bp) | 0.99, 1.29<br>(36402 bp) | 2.42, 4.94<br>(9249 bp) | 0.032, 0.040 | 75.6, 96.9 | 2.44, 3.83 |
| *N. panzeri*/*N. leucophthalma* | 0.033, 0.051<br>(613605 bp) | 1.00, 1.27<br>(36402 bp) | 2.25, 4.52<br>(9249 bp) | 0.033, 0.040 | 68.2, 88.6 | 2.25, 3.56 |
| *N. flava*/*N. leucophthalma* | 0.0088, 0.011<br>(613605 bp) | 0.65, 1.06<br>(36402 bp) | 1.34, 2.62<br>(9249 bp) | 0.014, 0.010 | 152.3, 238.2 | 2.06, 2.47 |

**Data sources and notes:** 1. Raychoudhury *et al.* 2009. 2. Darby *et al.* 2012. 3. Data from 387 protein-coding loci. For the *Onchocerca* pair, 572,142 bp of *Wolbachia* data are available. For consistency, the data were trimmed to the loci for which *Wolbachia* sequences from *Dirofilaria immitis* sequences were also available. 4. Nuclear and mtDNA genome data for the *Onchocerca* species and *Wolbachia* data for *Dirofilaria immitis* from the 50 Helminth Genomes Project: https://www.sanger.ac.uk/collaboration/50hgp/. 5. Gomes-de-Sa *et al.* 2022. 6. Ghedin *et al.* 2009. 7. Lau *et al.* 2015. 8. Data from 511 transcribed loci. For the *Brugia* pair, 598,257 bp of *Wolbachia* data are available. For consistency, the data were trimmed to the loci for which *Wolbachia* sequences from *Wuchereria bancrofti* were also available. 9. Small *et al.* 2019. 10. Gerth and Bleidorn 2016.

**Table 4.** Substitution rate estimates used to calibrate *Wolbachia* chronograms.

| Method | Substitutions/site/year |  | Data | Time calibration/validation |
| --- | --- | --- | --- | --- |
|  | third sites | synonymous sites |  |  |
| “Universal” estimate of the average substitution rate for coding DNA in bacteria applied to *Wolbachia ftsZ*<sup>1</sup> | NA | 7–8×10<sup>−9</sup> (ref. 2) | *ftsZ* differences among *Wolbachia*<sup>1</sup> | Divergence-time estimates based on 21 kb of coding DNA from bacteria *Salmonella typhimurium* and *Escherichia coli*<sup>2</sup> supplemented with data indicating similar synonymous-site substitution rates for *ftsZ* in *Wolbachia*<sup>1</sup>. |
| Divergence of mitochondrial vs. *Wolbachia* genomes among *Drosophila melanogaster* isofemale lines<sup>3</sup> | 6.87×10<sup>−9</sup> | NA | mtDNA and *Wolbachia* genomes | Compare sequence differences among isofemale lines for mtDNA and *Wolbachia*. Calibrate *Wolbachia* substitution rates using the per-generation mtDNA mutation rate as a prior for “short term” mtDNA (and *Wolbachia*) evolution<sup>3</sup>. |
| **Plausible example of codivergence of specific *Wolbachia* variants in two *Nasonia* species** |  |  |  |  |
| Updated universal synonymous-site substitution rate<sup>4</sup> for bacteria cross-validated with a host nuclear clock<sup>5</sup> for synonymous sites in *Wolbachia* (*wNlonB1* vs. *wNgirB*) putatively cladogenically transmitted between *Nasonia longicornis* and *N. giraulti* | 2.2×10<sup>−9</sup> | 3.3×10<sup>−9</sup> | Fragments of *Wolbachia* and host nuclear loci | Concordance of divergence times estimated from molecular clocks applied to *Wolbachia* and host nuclear DNA<sup>5</sup>. Our substitution-rate estimates assume MRCA for host and *Wolbachia* 0.46 MYA, averaging the bacterial and eukaryotic clock-based estimates from ref. 5. |
| **Codivergence of obligate symbiotic *Wolbachia* with filarial nematodes** |  |  |  |  |
| *Onchocerca volvulus*/*O. ochengi* | 1.38×10<sup>−9</sup><br>(0.73–3.44)×10<sup>−9</sup> | 2.11×10<sup>−9</sup><br>(1.11–5.25)×10<sup>−9</sup> | Draft *Wolbachia* and host nuclear genomes | Time back to MRCA of hosts (and 95% credible interval), 1.99 (0.80–3.79) MY, estimated from our relaxed-clock relative chronogram (Fig. 2B) for *Onchocerca* and *Dirofilaria immitis*, assuming 64.73 MYA as age of the MRCA of *Onchocerca volvulus* and *Dirofilaria immitis*. |
| *Dirofilaria immitis*/*Onchocerca* | 1.82×10<sup>−9</sup><br>(1.28–3.13)×10<sup>−9</sup> | 2.73×10<sup>−9</sup><br>(1.92–4.70)×10<sup>−9</sup> | Draft *Wolbachia* and host nuclear genomes | The fossil-calibrated estimate (and support interval) for the age of the MRCA for *Onchocerca volvulus* and *Dirofilaria immitis* is 64.73 (37.56–91.96) MYA<sup>6</sup>. Rates estimated by averaging the *Wolbachia* divergences from Table 3 between *Dirofilaria immitis* and the two *Onchocerca* species. |
| *Brugia malayi*/*B. pahangi* | 6.47×10<sup>−10</sup><br>(3.71–16.4)×10<sup>−10</sup> | 9.12×10<sup>−10</sup><br>(5.23–23.1)×10<sup>−10</sup> | Draft *Wolbachia* and host nuclear genomes | Time back to MRCA of *Brugia* hosts (and 95% credible interval), 6.80 (2.68–11.85) MY, from our relaxed-clock relative chronogram (Fig. 2C) for *Brugia* and *Wuchereria bancrofti*, assuming 28.97 MYA as age of the MRCA of *Brugia malayi* and *Wuchereria bancrofti* <sup>6</sup>. Support intervals for rates from support intervals for MRCA age. |
| *Wuchereria bancrofti*/*Brugia* | $7.42 \times 10^{-10}$<br>(4.42–16.8) $\times 10^{-10}$ | $1.05 \times 10^{-9}$<br>(6.28–23.8) $\times 10^{-10}$ | Draft *Wolbachia* and host nuclear genomes | The fossil-calibrated estimate (and support interval) for the age of the MRCA for *Brugia malayi* and *Wuchereria bancrofti* is 28.97 (12.8–48.59) MYA<sup>6</sup>. Rates estimated by averaging *Wolbachia* divergences from Table 3 between *W. bancrofti* and the two *Brugia* species. |
| **Three alternative *Wolbachia* calibrations based on the *Nomada* bee clade: (*N. ferruginata*, (*N. panzeri*, (*N. flava*, *N. leucophtalma*)))** |  |  |  |  |
| *Wolbachia* in *N. ferruginata* vs. three-species ingroup | $6.46 \times 10^{-10}$ | $1.03 \times 10^{-9}$ | Draft *Wolbachia* and host nuclear genomes<sup>7</sup> | Estimated time back to MRCA of hosts, 2.14 MY (p. 4, ref. 6). Rates estimated by averaging the three sequence divergence estimates in Table 3. |
| *Wolbachia* in *N. panzeri* vs. (*N. flava*, *N. leucophtalma*) (95% credible intervals based on approximate credible range of divergence times) | $1.31 \times 10^{-10}$<br>(0.84–2.29) $\times 10^{-10}$ | $2.06 \times 10^{-10}$<br>(1.32–3.59) $\times 10^{-10}$ | Draft *Wolbachia* and host nuclear genomes | Time back to MRCA of hosts and 95% credible interval, 1.24 (0.71–1.93) MY, obtained from our relative chronogram (Fig. 2A), assuming a crown age of 2.14 MY for the four-species clade. The rates were estimated by averaging the two sequence divergence estimates in Table 3. |
| *Wolbachia* in *N. flava* vs. *N. leucophtalma* (95% credible intervals based on approximate credible range of divergence times) | $5.71 \times 10^{-11}$<br>(3.36–12.9) $\times 10^{-11}$ | $7.14 \times 10^{-11}$<br>(4.20–16.2) $\times 10^{-11}$ | Draft *Wolbachia* and host nuclear genomes | Time back to MRCA of hosts and 95% credible interval, 0.77 (0.34–1.31) MY, obtained from our relative chronogram (Fig. 2A), assuming a crown age of 2.14 MY for the four-species clade. |

**References.** 1. Werren *et al.* 1995. 2. Ochman and Wilson 1987. 3. Richardson *et al.* 2012. 4. Ochman *et al.* 1999. 5. Raychoudhury *et al.* 2009. 6. Qing *et al.* 2025. 7. Gerth and Bleidorn 2016.

We used the support intervals for the *Dirofilaria* and *Onchocerca* divergence time (95% HPD CI of 37.56–91.96 MYA) and for the *Brugia* and *Wuchereria* (95% HPD CI of 12.8–48.59 MYA) MRCAs estimated by Qing *et al*. (2025) to approximate support intervals for the substitution rates of the codiverging *Wolbachia*, averaging the differences given in Table 3 between each congener and its respective outgroup. Substitution rates were estimated by dividing the estimated number of substitutions between lineages (after Jukes-Cantor correction) by these divergence time-estimates. The *Dirofilaria*-*Onchocerca* point estimate for the substitution rate per third site per year, 1.82×10^-9^ [with support interval (1.28–3.13)×10^-9^], is about 2.5 times the *Brugia*-*Wuchereria* point estimate, 7.42×10^-10^ [with support interval (4.42–16.8)×10^-10^]. However, the support intervals overlap, and the *Dirofilaria*-*Onchocerca* estimate is consistent with our revised estimate from the Raychoudhury *et al*. (2009) *Nasonia* example (see below).

Two additional rate estimates were available from the pairs of congeners. As noted above, we used relative chronograms for each host trio to estimate the MRCA ages for each pair. We used the point estimates for the MRCA ages of each trio, together with the support intervals for the relative ages of the congeners, to approximate support intervals for the MRCA ages of the congeners. The resulting point estimates and support intervals for the *Wolbachia* substitution rates appear in Table 4. In both cases, the congener-based rate estimates are consistent with the respective estimates based on comparisons to the outgroup. The range of the four point estimates is 6.47×10^-10^ to 1.82×10^-9^. It is notable that according to Qing *et al*. (2025), the age of the MRCA of these nematode trios is over 40 MYA, yet the *Wolbachia* substitution-rate estimates vary by only about a factor of three.

For each of our rate estimates, the support range extends from roughly half to roughly double the point estimate. When estimating chronograms, we approximated this uncertainty by multiplying the point estimate by a lognormal random variable, *X*, defined by *X* = Exp[Normal(μ = 0, σ = 0.353654)].

Setting the mean of the underlying normal to 0 produces a median for *X* of 1. The value for σ is chosen so that P[*X* < 0.5] = P[*X* > 2] = 0.025, approximating our empirical support intervals for the rates. The value for σ implies that E[*X*] = 1.06, so that the mean rate is close to the point estimate.

To produce plausible estimates of divergence times and conservative estimates of their biological and statistical uncertainty, we averaged the fastest and slowest of our nematode-based rates, and then constructed a “conservative plausible range” (CPR) for each estimate using the statistical lower bound from the fastest rate and the upper bound from the slowest rate. This CPR more realistically captures the biological and statistical uncertainty of our time estimates than would conventional support intervals based on the statistical uncertainty of a single calibration. In Table 4, we present these new rate estimates as well as older estimates based on the mutation-based *Drosophila melanogaster* estimate from Richardson *et al*. (2012), a *Nasonia* calibration based on plausible codivergence of a specific *Wolbachia* lineage between two *Nasonia* species, and three alternative rates estimated from *Nomada* data of Gerth and Bleidorn (2016). With firm evidence for codivergence of *Wolbachia* and hosts in the two trios of filarial nematodes, and fossil-based estimates of the nematode host divergence times, we believe they are more reliable than previous estimates from *Nasonia* and *Nomada*. Notably, the fastest *Nomada* rate is consistent with the slowest of our nematode-based calibrations, and the *Nasonia* rate is consistent with the fastest of our nematode-based calibrations. However, as shown in Table 4, there are two additional *Nomada* rate estimates, based on the ingroup species, (*N. panzeri*, *N. flava* and *N. leucophtalma*). Those estimates are much slower than any of our nematode estimates. These slower rates seem to be artifacts of horizontal *Wolbachia* acquisition in these three ingroup *Nomada*. The key observations are that (1) the most closely related *Nomada* hosts yield 10-fold slower *Wolbachia* rates, while (2) the estimated mtDNA and nuclear rates are relatively constant. Rather than invoking a recent 10-fold slowdown of *Wolbachia* evolution over the last million years, these data seem more plausibly explained by horizontal *Wolbachia* acquisition within the three-species *Nomada* ingroup.

For our *Wolbachia* chronograms, we focus on age estimates and support intervals derived from the nematodes. The alternative calibrations are considered further in our Discussion and Supplementary Information. Based on our new rate estimates (see Table 3), which were consistent with estimates from *Nasonia* and *Nomada*, the Richardson *et al*. (2012) rate seems too fast by about a factor of seven. Turelli *et al*. (2018) used the Richardson *et al*. (2012) rate. Using our nematode-based calibrations, we now estimate *w*Ri-*w*Suz divergence as 76 KYA (CPR: 14–218 KYA). The implications of this revision are considered in our Discussion.

### Applying nematode-based calibrations to *w*Mel-like *Wolbachia*

#### Estimating *w*Mel-like *Wolbachia* divergence times

Using our new nematode-based calibrations, we estimated the timescale of *w*Mel-like *Wolbachia* host switching. Fig. 4A presents an estimated *Wolbachia* chronogram for the variants presented in Fig. 1A. Using bifurcating genomic regions with little evidence of horizontal allele acquisition, these *Wolbachia* genomes diverged only about 206 KY**–**2.43 MYA (Fig. 4A, Table 5). The *w*Mel-like *Wolbachia* (*w*Dal, (*w*Sbr, *w*Tei)) in the most-diverged hosts (*Diachasma alloeum*, (*S. brevicornis*, *D.teissieri*)) diverged about 650 KYA, with a CPR of 132 KYA–1.7 MYA. *i.e.* over a timescale comparable to a single host speciation event (Coyne and Orr 2004). The estimate for *w*Mel-like divergence is based on the *w*Dal-*w*SYT (*i.e.,* the clade including *w*San, *w*Yak, and *w*Tei) node in Fig. 4A, which is the smallest clade that includes *w*Dal, *w*Sbr, and *w*Tei. Among *w*Mel-like *Wolbachia* observed in 18 drosophilids in our study (Fig. 1), *w*Zts in tropical *Zaprionus tscasi* is the most closely related to *w*Mel in *D. melanogaster*, diverging about 336 KYA (CPR: 67–924 KYA). We are unaware of any other *Wolbachia* more closely related to *w*Mel (Supplemental Information, Fig. S1), although we expect others will be discovered as additional *Wolbachia* are sequenced. *Zaprionus* flies are members of the (paraphyletic) *Drosophila* subgenus that diverged from the *Sophophora* subgenus about 47 MYA (95% CI: 43.9–49.9 MYA), highlighting proliferation of *w*Mel-like *Wolbachia* across species that span the entire paraphyletic genus also named *Drosophila* (Suvorov *et al*. 2022).

**Figure 4.**
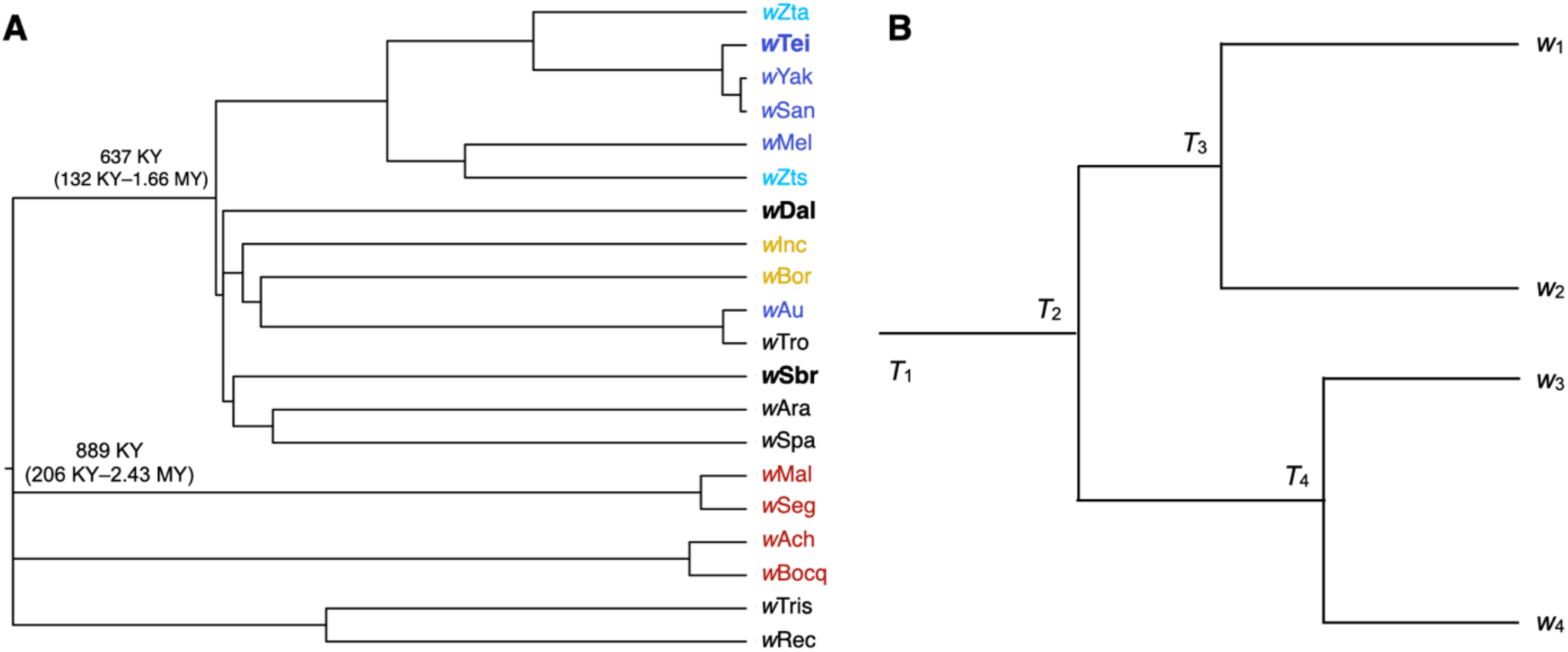
(A) The timescale of *w*Mel-like *Wolbachia* movements and limitations of inferences. An absolute chronogram for the *w*Mel-like *Wolbachia* in our study. Labels for the *Wolbachia* associated with the most distantly related hosts presented in Fig. 1B are in bold and include: *w*Dal (in *D. alloeum*), *w*Sbr (in *S. brevicornis*) and *w*Tei (in *D. teissieri*) diverged 637 KY, with a CPR of 132 KY–1.66 MY. The colored *Wolbachia* labels match the host clades presented in Fig. 1B. The crown age is 889 KY, with CPR of 206 KY–2.43 MY. The several polytomies reflect the rapid periods of diversification noted in Fig. 1A. **(B)** An illustration clarifying the limitations of inferences from a chronogram based on single *Wolbachia* genomes from individual hosts. These limitations do not affect our central inferences concerning the timing of *Wolbachia* movement and molecular evolution.

**Table 5.** *w*Mel-like *Wolbachia* node age point estimates and 95% CIs in years for all nodes presented in Fig. 4A. We present estimates produced using five different calibrations described in the Materials and Methods and Results sections. The final columns provide our plausible point estimates calculated as the average of the point estimates of the fastest (*Dirofilaria*, lognormal) and slowest (*Brugia*, lognormal) nematode-based calibrations. We report the conservative plausible range (CPR), which ranges from the lower-bound of the fastest and the upper-bound of the slowest nematode-based calibrations. As described in the text, the range of our new nematode-based calibrations are consistent with estimates from *Nasonia* and *Nomada*. In contrast, the mutation-based rate estimate of Richardson *et al*. (2012) is significantly faster (Table 4).

|  | *Drosophila*<br>mtDNA |  | *Dirofilaria* (fixed) |  | *Dirofilaria*<br>(lognormal) |  | *Brugia* (fixed) |  | *Brugia* (lognormal) |  | Plausible Time<br>Estimates |  |
| --- | --- | --- | --- | --- | --- | --- | --- | --- | --- | --- | --- | --- |
| Node | Point | 95% CI | Point | 95% CI | Point | 95% CI | Point | 95% CI | Point | 95% CI | Point | CPR |
| Root | 122104 | (98666,<br>152513) | 460910 | (372437,<br>575693) | 466869 | (206051,<br>900601) | 1296533 | (1047660,<br>1619410) | 1311857 | (618461,<br>2433090) | 889363 | (206051,<br>2433090) |
| (*wSan,wYak*) | 877 | (157,<br>2248) | 3311 | (593,<br>8488) | 3330 | (496,<br>10133) | 9314 | (1670,<br>23878) | 9498 | (971,<br>28857) | 6414 | (496,<br>28857) |
| (*wTeis,(wSan,wYak)*) | 3904 | (1760,<br>7043) | 14739 | (6645,<br>26588) | 14777 | (4948,<br>34677) | 41461 | (18692,<br>74791) | 42241 | (12162,<br>93988) | 28509 | (4948,<br>9398) |
| (*wZta,(wTeis,(wSan,wYak)*) | 35452 | (22890,<br>48999) | 133824 | (86406,<br>184959) | 134736 | (46652,<br>266041) | 376446 | (243059,<br>520287) | 376929 | (130538,<br>734069) | 255832.5 | (46652,<br>734069) |
| (*wMel, wZts*) | 46829 | (35337,<br>61459) | 176766 | (133390,<br>231991) | 177819 | (66835,<br>333151) | 497242 | (375224,<br>652588) | 494263 | (195489,<br>924345) | 336041 | (66835,<br>924345) |
| ((*wMel,wZts*),(*wZta,wSYT*)) | 59793 | (48403,<br>73030) | 225702 | (182710,<br>275670) | 227964 | (96325,<br>434420) | 634896 | (513960,<br>775454) | 640685 | (268781,<br>1161800) | 434324.5 | (96325,<br>1161800) |
| (*wTro,wAu*) | 3796 | (1395,<br>7104) | 14332 | (5267,<br>26818) | 14860 | (4106,<br>35845) | 40315 | (14818,<br>75439) | 41231 | (10924,<br>93382) | 28045.5 | (4106,<br>93382) |
| (*wBor,(wTro,wAu)*) | 80825 | (68085,<br>94318) | 305093 | (257003,<br>356026) | 309441 | (134625,<br>595353) | 858222 | (722944,<br>1001500) | 866659 | (370028,<br>1562040) | 588050 | (134625,<br>1562040) |
| (*wInc,(wBor,(wTro,wAu))*) | 83855 | (71365,<br>96353) | 316530 | (269383,<br>363707) | 320611 | (125589,<br>601643) | 890395 | (757771,<br>1023100) | 900733 | (387630,<br>1600610) | 610672 | (125589,<br>1600610) |
| (*wAra,wSpa*) | 78828 | (60822,<br>95936) | 297556 | (229588,<br>362133) | 297397 | (126652,<br>584033) | 837021 | (645826,<br>1018670) | 842347 | (312464,<br>1521920) | 569872 | (126652,<br>1521920) |
| (*wSbr,(wAra,wSpa)*) | 85407 | (72389,<br>94497) | 322389 | (273248,<br>371800) | 324318 | (134375,<br>617210) | 906875 | (768643,<br>1045870) | 919205 | (371034,<br>1620360) | 621761.5 | (134375,<br>1620360) |
| (*wDal,wAra-wSbr,wTro-wInc*) | 87120 | (75334,<br>100665) | 328856 | (284368,<br>379984) | 332375 | (132290,<br>622578) | 925068 | (799922,<br>1068890) | 940738 | (394476,<br>1658090) | 636556.5 | (132290,<br>1658090) |
| *wSYT-wDal* | 88309 | (77169,<br>101950) | 333344 | (291294,<br>384833) | 337106 | (134946,<br>627989) | 937691 | (819406,<br>1082530) | 949467 | (401110,<br>1684930) | 643286.5 | (134946,<br>1684930) |
| (*wMal,wSeg*) | 7496 | (3328,<br>12777) | 28297 | (12564,<br>48230) | 29227 | (7380,<br>62302) | 79600 | (35343,<br>135672) | 81049 | (26366,<br>174569) | 55138 | (7380,<br>174569) |
| (*wBocq,wAch*) | 9404 | (4754,<br>15762) | 35500 | (17948,<br>59497) | 36314 | (11089,<br>76932) | 99861 | (50488,<br>167354) | 100637 | (29261,<br>221975) | 68475.5 | (11089,<br>221975) |
| (wTris,wRec) | 69956 | (44884,<br>113664) | 264067 | (169425,<br>429051) | 272767 | (86711,<br>565736) | 742817 | (476590,<br>1206910) | 764425 | (246849,<br>1566220) | 518596 | (86711,<br>1566220) |

#### Determining *w*Mel-like *Wolbachia* acquisition modes

Many obligate mutualistic endosymbionts like the *Wolbachia* in filarial nematodes (Comandatore *et al*. 2013) and *Buchnera* in aphids (Baumann *et al*. 1995) are acquired cladogenically. In contrast, because the divergence times for our *w*Mel-like *Wolbachia* are so much shorter than for their hosts, all but one of these variants must have been acquired through introgression or non-sexual horizontal transfer (though as explained in the next subsection, we cannot exclude that one host cladogenically acquired its *Wolbachia* from a closely related host we did not sample). Introgression is plausible only between the most closely related drosophilid species in our study (*i.e.,* within the colored triangles in Fig. 1B), some of which may hybridize (Matute and Cooper 2021).

Confirming introgressive acquisition requires demonstrating concordant divergence times for mtDNA and the associated *Wolbachia*, and these must be more recent than host divergence.

Joint analysis of mtDNA and *Wolbachia* sequence divergence in Cooper *et al*. (2019) implied that the three-species *D. yakuba* clade (*D. teissieri*, (*D. yakuba*, *D. santomea*)) first acquired *Wolbachia* by horizontal transfer from an unknown host. The *Wolbachia* were then transferred within the clade through hybridization and introgression. *D. yakuba* hybridizes with its endemic sister species *D. santomea* on the island of São Tomé (Lachaise *et al*. 2000; Comeault *et al*. 2016; Cooper *et al*. 2017), and with *D. teissieri* on the edges of forests on the nearby island of Bioko (Cooper *et al*. 2018). *Wolbachia* and mtDNA chronograms were generally concordant for these three hosts and indicated more recent common ancestry for these maternally inherited cytoplasmic factors than for the bulk of their hosts’ nuclear genomes (Cooper *et al*. 2019).

*w*Zta from *Z. taronus* is the most closely related *Wolbachia* to *w*SYT, diverging about 256 KYA (CPR: 47–734 KYA). *Z. taronus* also occurs on São Tomé, particularly co-occurring with *D. santomea* at high altitudes on Pico de São Tomé; but it diverged from the *D. yakuba* triad about 47 MYA (Fig. 1B), making introgression between *Z. taronus* and the *D. yakuba*-clade species impossible (*cf.* Coyne and Orr 2004). Yet, its *Wolbachia* (*w*Zta) diverged from the *w*Yak-clade *Wolbachia* only about 46–734 KYA (Fig. 4A). These data illustrate horizontal *Wolbachia* transfer between distantly related species with overlapping ranges and habitats. The mechanisms of horizontal transfer are largely unresolved, but plausible alternatives are considered in our Discussion.

To identify possible cases of introgressive *Wolbachia* transfer, we compared estimated divergence times for mtDNA and *Wolbachia*. Table 6 presents the estimated mtDNA substitution rates per third-position site per year for the five *Drosophila* pairs described in our Materials and Methods, comparing rates derived from recently diverged species, which may plausibly hybridize, with the slower rates estimated from more distantly related species unlikely to produce fertile hybrids. To calibrate mtDNA divergence, we averaged the point estimates for mtDNA substitution rates per third-position site per year for *D. melanogaster*-*D. simulans* [1.56×10^-8^ with support interval (1.28–1.93)×10^-8^] and *D. erecta*-*D. yakuba* [1.76×10^-8^ with support interval (1.50–2.10)×10^-8^]. We used the average rate, 1.66×10^-8^, to estimate mtDNA divergence times for relatively recently diverged *Drosophila*. We considered the fastest and slowest mtDNA substitution rates across both credible intervals as a conservative substitution-rate range (1.28×10^-8^, 2.10×10^-8^).

**Table 6.** Rates used to estimate mtDNA divergence times. Five *Drosophila* species with increasing divergence times were used to compute mtDNA 3^rd^-position differences and estimate mtDNA substitution rates. As described in the main text, we consider the average of the rates derived from the two most recently diverged species pairs (*i.e.,* 1.66×10^-8^) as a plausible point estimate for mtDNA substitutions per third-position site per year between pairs of species that may hybridize. We consider the fastest and slowest substitution rates across the credible intervals for these two pairs as a conservative range of substitution rates (1.28×10^-8^, 2.09×10^-8^) for approximating mtDNA divergence times for pairs of hosts with *w*Mel-like *Wolbachia*. MRCA time estimates and confidence intervals for the hosts are taken from Suvorov *et al*. (2022). Point estimates and confidence intervals for mtDNA rates are calculated from the observed mtDNA divergences and MRCA time estimates.

| Species pair | MRCA (Point, MY)<sup>1</sup> | MRCA (95% CI, MY)<sup>1</sup> | mtDNA Length Compared (bp) | mtDNA 3<sup>rd</sup>-position differences (tip-to-tip)<sup>2</sup> | mtDNA substitutions per 3<sup>rd</sup>-position site per year (Point) | mtDNA substitutions per 3<sup>rd</sup>-position site per year (CI) |
| --- | --- | --- | --- | --- | --- | --- |
| *D. melanogaster*-*D. simulans* | 3.62 | (2.92–4.4) | 11,184 | 11.3% | $1.56 \times 10^{-8}$ | ($1.28 \times 10^{-8}$, $1.93 \times 10^{-8}$) |
| *D. erecta*-*D. yakuba* | 5.47 | (4.59–6.38) | 11,172 | 19.2% | $1.76 \times 10^{-8}$ | ($1.50 \times 10^{-8}$, $2.09 \times 10^{-8}$) |
| *D. mojavensis*-*D. virilis* | 24.34 | (21.94–26.88) | 11,164 | 33% | $6.78 \times 10^{-9}$ | ($6.14 \times 10^{-9}$, $7.52 \times 10^{-9}$) |
| *D. pseudoobscura*-*D. suzukii* | 32.78 | (29.77–35.76) | 11,182 | 29.4% | $4.48 \times 10^{-9}$ | ($4.11 \times 10^{-9}$, $4.94 \times 10^{-9}$) |
| *D. grimshawi*-*D. willistoni* | 46.84 | (43.85–49.85) | 11,184 | 32.6% | $3.48 \times 10^{-9}$ | ($3.27 \times 10^{-9}$, $3.72 \times 10^{-9}$) |

<sup>1</sup>Nuclear MRCA based on the fossil-calibrated chronogram of Suvorov *et al.* (2022).
<sup>2</sup>Nucleotide differences were counted and the Jukes-Cantor correction for multiple substitutions was applied.

We tested for introgressive *w*Mel-like *Wolbachia* transfer between four species pairs for which hybridization and introgression may be possible by comparing *Wolbachia* and mtDNA divergence-time estimates with estimates of the hosts’ divergence time. We considered two relatively closely related species pairs in the *D. montium* subgroup, (*D. seguyi*, *D. malagassya*) and (*D. bocqueti*, *D.* sp. aff. *chauvacae*) that diverged approximately 2.3 MYA (95% CI: 1.1–4.1 MYA) and 2.5 MYA (95% CI: 2.0–3.1 MYA), respectively (Fig. 1B). Their mtDNA third-position coding sites differed by 0.53% and 1.15%, respectively. Applying the Jukes-Cantor formula to estimate the numbers of substitutions and our mtDNA calibration, we estimated that the mtDNA of *D. seguyi* and *D. malagassya* diverged approximately 160 KYA (with plausible interval 126–207 KYA), suggesting mtDNA introgression.

Given our CPR of *w*Seg-*w*Mal *Wolbachia* divergence (7–174 KY), we determined that introgressive *Wolbachia* acquisition is possible, but we could not rule out horizontal acquisition after mtDNA introgression. In contrast, the mtDNA of *D. bocqueti* and *D.* sp. aff. *chauvacae* diverged approximately 346 KYA (with plausible interval 274–449 KYA), while their *w*Bocq-*w*Ach *Wolbachia* diverged more recently (CPR: 11–222 KY) (Fig. 4A). Thus, we inferred horizontal *Wolbachia* acquisition.

The more distantly related pairs (*Z. taronus*, *Z. tsacasi*) and (*D. borealis*, *D. incompta*) diverged approximately 14.2 MYA (95% support interval: 12.3–16.6 MYA) and 8.2 MYA (95% support interval: 3.7–14.5 MYA), respectively. For these pairs, mtDNA third-position coding sites differed by 19.3% and 30%, respectively (Suvorov *et al*. 2022), making mtDNA introgression plausible for the *Zaprionus* pair. Applying the mtDNA rate 1.66×10^-8^ after Jukes-Cantor correction, we estimated mtDNA divergence times on the order of 5.8 MYA (with plausible interval 4.6–7.5 MYA) for *Z. taronus* and *Z. tsacasi* and 9 MYA (with plausible interval 7.1–11.7 MYA) for *D. borealis* and *D. incompta*. The latter argued against mtDNA introgression between *D. borealis* and *D. incompta.* In comparison, our plausible divergence interval for their *Wolbachia*, *w*Bor and *w*Inc, was (126 KY–1.6 MY), which strongly supported horizontal *Wolbachia* acquisition. The shorter estimated mtDNA divergence time for the *Zaprionus* pair may have reflected a slowdown in mtDNA divergence associated with saturation (Ho *et al*. 2005), as illustrated in Table 6 for the three more diverged *Drosophila* species pairs. However, irrespective of whether mtDNA introgression occurred in the *Zaprionus* pair, the CPR for *w*Zta-*w*Zts divergence (96 KY–1.2 MY) was much shorter than plausible mtDNA divergence-time estimates, again supporting horizontal *Wolbachia* acquisition.

In summary, we ruled out introgressive *Wolbachia* acquisition for three of the four possible cases we examined. Introgression was consistent with our analyses only for *w*Seg and *w*Mal, but horizontal acquisition could not be confidently rejected.

### Identifying limitations of inferences from *Wolbachia* phylogenies and chronograms

Inferences from our chronograms of *w*Mel-like variants are limited because we considered only one *Wolbachia* sequence from each host. Specifically, we cannot determine which *Wolbachia* infection was acquired first, whether two sister tips on our *Wolbachia* chronograms correspond to one of those hosts acquiring its *Wolbachia* (by either introgression or non-sexual horizontal transmission) from the host of the sister *Wolbachia* variant – or whether one or both *Wolbachia* were recently acquired from an unsampled intermediate host. Nevertheless, our central inferences concerning the timing of *Wolbachia* movement and molecular evolution should be robust. Turelli *et al*. (2018) previously proposed that *w*Ri in *D. simulans* was acquired horizontally from *D. ananassae*, because multiple *Wolbachia* sequences were available from each species, and the *w*Ri sequences formed a clade nested within a paraphyletic cluster containing the *Wolbachia* from *D. ananassae*. As noted by Richardson *et al*. (2012), divergence times for extant *Wolbachia* variants in a host species may be much shorter than the duration of its current *Wolbachia* variant, because selective sweeps and genetic drift will eliminate sequence variation that might provide information about the duration of associations.

Fig. 4B illustrates the inferences about *Wolbachia* divergence and acquisition possible from individual *Wolbachia* sequences from various hosts. We considered four hypothetical host species with horizontally acquired *Wolbachia*. Host 1 acquired its *Wolbachia*, denoted *w*_1_, at time T_1_, and we denote by T_i_(j » i) the time at which host i acquired its *Wolbachia*, denoted *w*_i_, from host j. Note that T_1_ may be much longer than T_2_, T_3_ and T_4_, corresponding possibly to cladogenic inheritance of *w*_1_. Our chronogram of wMel-like variants says nothing about the length of this root branch. If we postulate T_1_ > T_2_(1 » 2) > T_3_(1 » 3) > T_4_(2 » 4), the resulting chronogram is Fig. 4B. The chronogram provides no indication that variant *w*_1_ was acquired first (or how it was acquired) or that *w*_2_ was acquired from host 1. Thus, we can infer only that all but one of the variants shown in Fig. 4A were acquired in the last 206 KY–2.43 MY and that a transfer between a dipteran and a hymenopteran host occurred within the last 132 KY– 1.66 MY, possibly mediated by intermediate hosts. Moreover, the chronogram Fig. 4A is consistent with many of these hosts, for which we do not have multiple *Wolbachia* sequences, acquiring their current *Wolbachia* very recently, followed by spatial spread in the new host, as observed for several *Wolbachia*=-host associations (Turelli *et al*. 2018).

### Estimating *Wovirus* and *cif* turnover in *w*Mel-like and *w*Ri-like *Wolbachia*

We next applied our nematode-based calibration to comparisons of closely related *w*Mel-like and *w*Ri-like *Wolbachia* to understand the timescale of *Wovirus* and *cif* turnover occurring outside the bifurcating genomic regions. While *Wolbachia* genome quality varied among our samples, the timescale estimates below are established by focal comparisons involving fully circularized genomes and based on divergence times of the bifurcating genomic regions.

### Estimating *Wovirus* turnover

We used serine recombinase (sr) alleles to type *Woviruses* in *w*Mel-like and *w*Ri-like genomes (see our Materials and Methods), focusing on alleles (sr1-sr3) of three *Wovirus* Types that are known to contain *cifs* (sr1WO-sr3WO) (Bordenstein and Bordenstein 2022).

Table S9 summarizes our results. While we did not observe sr1WO in any *w*Mel-like *Wolbachia*, identical sr1 alleles were observed in all *w*Ri-like *Wolbachia*, except for *w*Aur and *w*Tri. Based on the CPR of *w*Ri-like variant divergence, this implies loss of sr1WO by *w*Aur and *w*Tri in the last 14–218 KYA. We observed closely related sr2WO in all *w*Ri-like *Wolbachia*, with each sr2 allele differing from the copy in *w*Ri at ≤ 2 sites. Among *w*Mel-like *Wolbachia*, an sr2WO was observed only in *w*SYTZ (*w*SYT plus the *w*Zta outgroup) and in *w*Tris genomes (Fig. S2A). The absence of this sr2WO in the *w*Mel and *w*Zts genomes implies gain or loss by *w*SYTZ in the last 96 KYA–1.16 MYA. The *w*SYT and *w*Mel genomes are fully circularized. The absence of this sr2WO in the *w*Rec genome, implies gain or loss since its divergence from *w*Tris 87 KYA–1.57 MYA. sr3WO *Wovirus* was more common, occurring in all *w*Ri-like and *w*Mel-like genomes, and often in multiple copies (Fig. S2B, Table S9). Analysis of intralocus recombination using GARD identified two breakpoints in sr3 that we visually confirmed, resulting in three partitions. Nodes within partitions were often unresolved and relationships between partitions often varied (Fig. S3; Supplemental Information). Hence we refrained from explicitly inferring the timescale of sr3WO gains and losses because this inference would rely on the sr3 phylogram.

Ignoring sr3WO, our conservative comparisons indicate that *Woviruses* based on sr typing can be gained and lost on the order of about 10^4^**–**10^6^ years.

### Estimating *cif* turnover

CI is common (Fig. 5A) and *cif* complements are diverse (Fig. 5B) among *w*Mel-like and *w*Ri-like *Wolbachia*, with 14/21 characterized strains in our study causing CI (Fig. 5A; Table S8; Supplemental Information) (Martinez *et al*. 2021; Turelli *et al*. 2018). This includes two *w*Mel-like *Wolbachia* (*w*Seg in *D. seguyi* and *w*Bocq in *D. bocqueti*), which we confirmed cause relatively intense CI (Supplemental Information). *w*Mel-like and *w*Ri-like *Wolbachia* carry between one and three *cif* operons that span four *cif* Types (Fig. 5B; Table S9; Supplemental Information). Applying our nematode-based calibrations to comparisons of *cifs* observed in closely related *Wolbachia*, we confirmed that rapid *cif* turnover contributes to this diversity. Among *w*SYT variants, two *cif*_[T4]_ operons are observed in *w*San and *w*Yak, while *w*Tei contains only one (Baião *et al*. 2021). These two copies are identical, implying plausible duplication of a *cif*_[T4]_ operon after *w*SYT diverged, and prior to *w*San-*w*Yak divergence 10^3^–10^5^ years ago. *w*Mel and *w*Zts genomes did not contain *cif*_[T4]_ operons, indicating acquisition of *cif*_[T4]_ operons by *w*SYTZ occurred 96 KYA–1.16 MYA. (With the exception of *w*Zta, all genomes in this comparison are fully circularized.) In the ((*w*Ara, *w*Spa), *w*Sbr) clade (CPR: 134 KYA–1.62 MYA), we observed turnover of multiple *cif* Types: *w*Ara contained two *cif*_[T1]_ operons, its sister *w*Spa contained one *cif*_[T1]_ operon and one *cif*_[T2]_ operon, and outgroup *w*Sbr contained only *cif*_[T2]_ and *cif*_[T5]_ operons. Similarly, in the (*w*Inc, (*w*Bor, (*w*Au, *w*Tro))) clade (CPR: 126 KYA–1.6 MYA), *w*Inc and *w*Bor each contained a *cif*_[T1]_ operon, but *w*Bor contained a *cif*_[T5]_ operon, and *w*Au and *w*Tro did not contain any *cif* operons. Finally, all *w*Ri-like variants contained a single *cif*_[T2]_ operon, with the exception of *w*Aur and *w*Tri, that instead had a *cif*_[T5]_ operon absent from all other *w*Ri-like genomes, including the fully circularized genomes of *w*Ri and *w*Ana. This implies turnover of both *cif* Types since the *w*Ri-clade diverged 14–218 KYA (Table 7).

**Figure 5.**
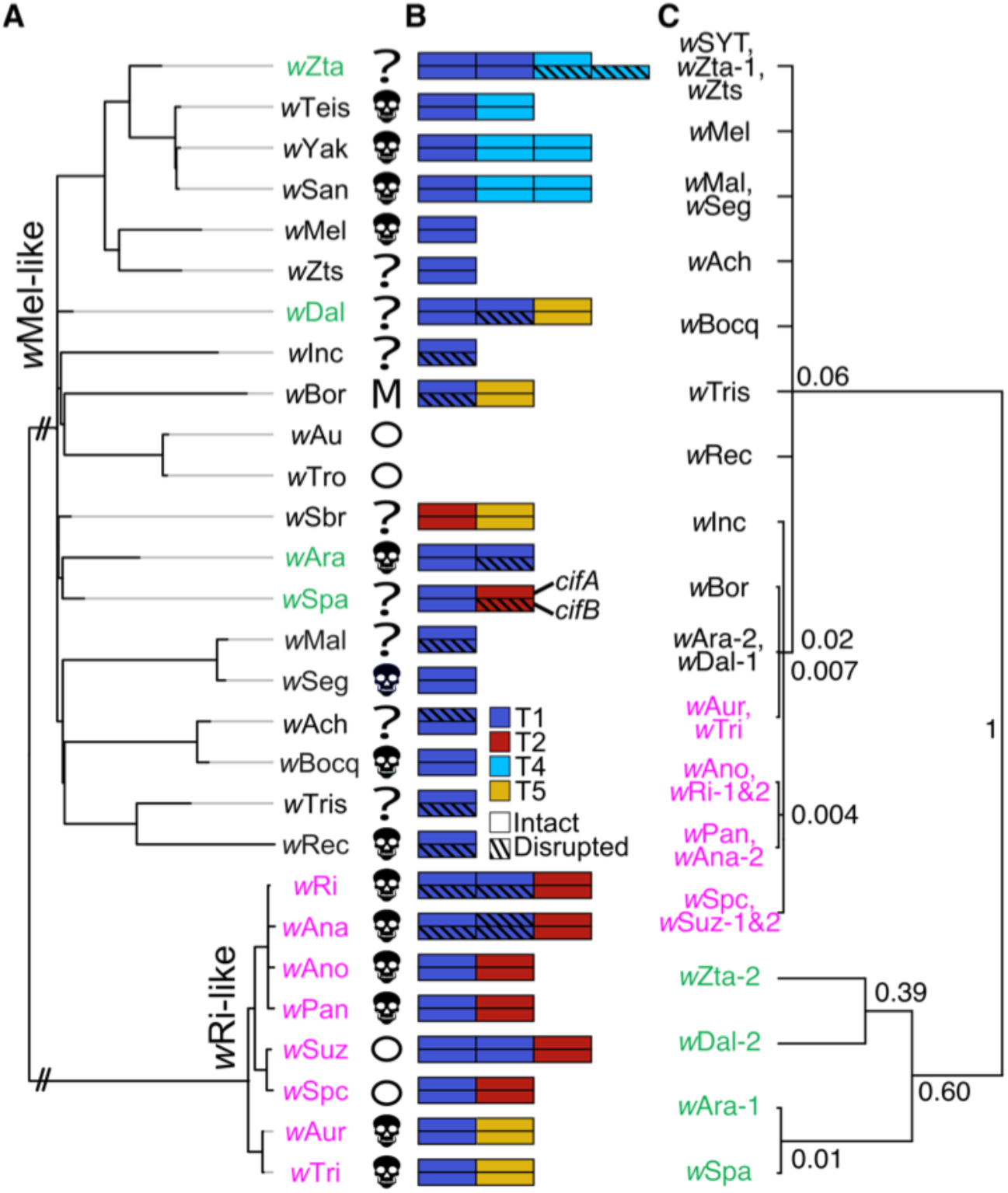
Diverse *cif* operons rapidly turning over among *w*Mel-and *w*Ri-like genomes. **(A)** A phylogram of *w*Mel-like (black and green) and *w*Ri-like (pink) *Wolbachia*, including variants that cause CI 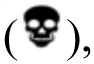 do not cause CI (circles), or where CI status is unknown CI (?). *w*Bor does not cause CI, but it does kill males (M) (Table S8). Our new calibrations indicated that the *w*Ri-like clade diversified over about 14–218 KY, the *w*Mel-like clade diversified over about 206 K–2.4 MY, and the MRCA for the bifurcating genomic regions of *w*Mel and *w*Ri is about 1.4–22 MYA. Branches leading to these clades are shortened for presentation (//), and light gray branch extensions are used to improve visualization. **(B)** Across *w*Mel-like and *w*Ri-like *Wolbachia*, four of the five characterized *cif* operon Types (T1–T5) are observed (Table S9). *cifA* (top) and *cifB* (bottom) schematics are presented with multiple copies adjacent to one another and ordered by their sequence integrity. The inset legend denotes *cif* Type by color, with integrity classified as intact (fully shaded) or disrupted (hashed) based on whether mutations split the gene into multiple ORFs or truncate the protein, causing loss of a functional domain. **(C)** A relative chronogram for *cifA*_[T1]_ copies with node labels indicating relative ages, scaled to 1 for the root node.

**Table 7.** Estimates of root ages of four sets of *Wolbachia*: *w*Mel-like (*N* = 20), *w*Ri-like (*N* = 8), both clades combined (*N* = 28) (which span the seven supergroup A *Wolbachia* described in our Materials and Methods and in our Results [(((*w*Mel, *w*Au), (*w*Bic, *w*Bar)), ((*w*Ri, *w*Tri), *w*Ha))]), and A and B supergroup members from Fig. 1 in Meany *et al*. (2019) (*N* = 15 variants). Plausible point estimates are calculated as the average of the point estimates of the fastest (*Dirofilaria*, lognormal) and slowest (*Brugia*, lognormal) nematode-based calibrations. We also report the CPR, which ranges from the lower-bound of the *Dirofilaria* range to the upper-bound of the *Brugia* range.

| Plausible Time Estimate |  |  |
| --- | --- | --- |
| Comparison: | Point | CPR |
| wMel-like root | 889,363 | (206,051; 2,433,090) |
| wRi-like root | 76,421 | (14,051; 218,089) |
| wMel-like and wRi-like root | 7,545,595 | (1,441,730; 22,004,200) |
| A-B supergroup root | 151,570,339 | (34,100,400; 414,809,000) |

Because *cif*_[T1]_ operons are the most common *cif* Types in our dataset (Table S9), we focused on homologs of *cifA*_[T1]_ to further illustrate rapid *cif* turnover. We present a relative chronogram for *cifA*_[T1]_ copies (Fig. 5C). We observed two distantly related clades of *cifA*_[T1]_ alleles (Fig. 5C). The first included 26 alleles observed across 16 *w*Mel-like *Wolbachia* and 8 *w*Ri-like *Wolbachia. w*Ri and *w*Suz each carry two closely related *cifA*_[T1]_ alleles (Table S10). In both *w*Ri and *w*Suz, these alleles occurred 8,019 bp from an sr, implying ancestral duplication of the phage is plausible (Supplemental Information). *cifA*_[T1]_ alleles observed in *w*Mel-like *w*Dal, *w*Ara, *w*Bor and *w*Inc were most closely related to *cifA*_[T1]_ alleles observed in *w*Ri-like *Wolbachia*. They shared particularly high identity with *cifA*_[T1]_ alleles observed in *w*Ri-like *w*Aur and *w*Tri (99.8–99.2% aa identity). The second distantly related *cifA*_[T1]_ clade included additional *cifA*_[T1]_ copies observed in *w*Mel-like *w*Zta, *w*Dal, and *w*Ara genomes that were more closely related to each other than they are to the second *cifA*_[T1]_ copy in their respective *Wolbachia* genomes. The relationships of copies within this second clade, which also contained a single *cifA*_[T1]_ allele observed in *w*Spa, differed across two *cifA*_[T1]_ partitions based on GARD recombination calls (Fig. S4; Supplemental Information). Finally, we observed that many *cifA*_[T1]_ copies of high sequence similarity spanned relatively deep phylogenetic divergences among plausibly associated sr3 alleles (Fig. S5), a pattern indicative of *cif* turnover among *Woviruses* (LePage *et al*. 2017; Supplemental Information).

In summary, we conclude that *cif* gains and losses occurred in *w*Mel-like and *w*Ri-like genomes on the order of about 10^4^–10^6^ years, excluding cases that likely involved duplications, which occurred even more rapidly. In the Discussion, we consider plausible mechanisms that may have contributed to these patterns.

Identical sequences are collapsed into a single tip, and nodes with posterior probability < 0.95 are collapsed into polytomies. The majority of *cifA*_[T1]_ operons were closely related, with four divergent copies colored in green.

Colors in C correspond to those for hosts in panel A. When multiple copies of the same Type were present in a genome and complete genomes were available, the dash and number following the variant name indicates the order in which copies occurred in their respective genomes. In cases where complete genomes are not available, copy numbers are assigned alphabetically based on NCBI accession names for contigs. All *cifA*_[T1]_ copies are represented, with the exception of *w*Ana-1 that could not be confidently aligned. In the Supplemental Information we present phylograms for *cifA*_[T1]_ partitions based on GARD calls (Fig. S4), which differed in the specific placement of the four outgroup copies.

### Estimating selection on *cifs*

CI phenotypes and the *cifs* that underlie them are common among *w*Mel-like, *w*Ri-like and other *Wolbachia* (Fig. 5A; Tables S8 & 9) (Shropshire *et al*. 2020a; Turelli *et al*. 2022). Putative loss-of-function mutations are more commonly observed and occur faster in *cifB* than in the *cifA* rescue locus (Martinez *et al*. 2021). These patterns also emerged from our comparisons of Cif protein similarity and identity (Table S10). For example, CifA_[T1_ proteins were more similar than CifB_[T1]_ proteins from the same pairs (Figs. 5A & S6; Supplemental Information). To further understand these observations, we tested for selection during the divergence of putatively intact *cifA*_[T1]_ and *cifB*_[T1]_ alleles with no observed disruptions. Our analyses focus on Type 1 loci because they are common among our *w*Mel-like and *w*Ri-like variants (Fig. 5B). For *cifB*_[T1]_, we also assessed selection on domains involved in CI and other functions, relative to its non-domain regions (Beckmann *et al*. 2017; LePage *et al*. 2017; Chen *et al*. 2019; Beckmann *et al*. 2021; Kaur *et al*. 2022; Kaur *et al*. 2024; Deehan *et al*. 2021). These include two PD-(D/E)XK nuclease domains (Nuc1 and Nuc2) and a single Deubiquitinase domain (Dub). We analyzed the estimated normalized ratio of non-synonymous to synonymous substitutions (*ω*), comparing *cifA*_[T1]_ and *cifB*_[T1]_, as well as *cifB*_[T1]_ domains to non-domain regions.

Across all pairwise contrasts, *ω* for *cifA*_[T1]_ (range: 0.26–0.38) was consistently and significantly lower than *ω* for *cifB*_[T1]_ (range: 0.34–0.46) (Fig. 6B; HL median difference = −0.06; Wilcoxon *V* = 0, *P* = 0.02; Table S11), with Bayesian bootstrap posteriors indicating with high probability (*P* = 1.0) that *cifA*_[T1]_ *ω* was lower, *i.e., P*(Δ < 0) where Δ = *ω_cifA_* – *ω_cifB_*. Sliding window analyses assessed in 3D space agreed, with pooled medians for *ω* across all windows lower for CifA_[T1]_ (0.29) than CifB_[T1]_ (0.37) (Δ*_p_* =−0.23; BCa 95% CI: −0.38, −0.08; *P* < 0.0002; Supplemental Information). Estimates of *ω* imply selection acts to preserve intact copies of *cifA*_[T1]_ and *cifB*_[T1]_, with lower *ω* for *cifA*_[T1]_ implying this locus involved in CI rescue is under greater constraint.

**Figure 6.**
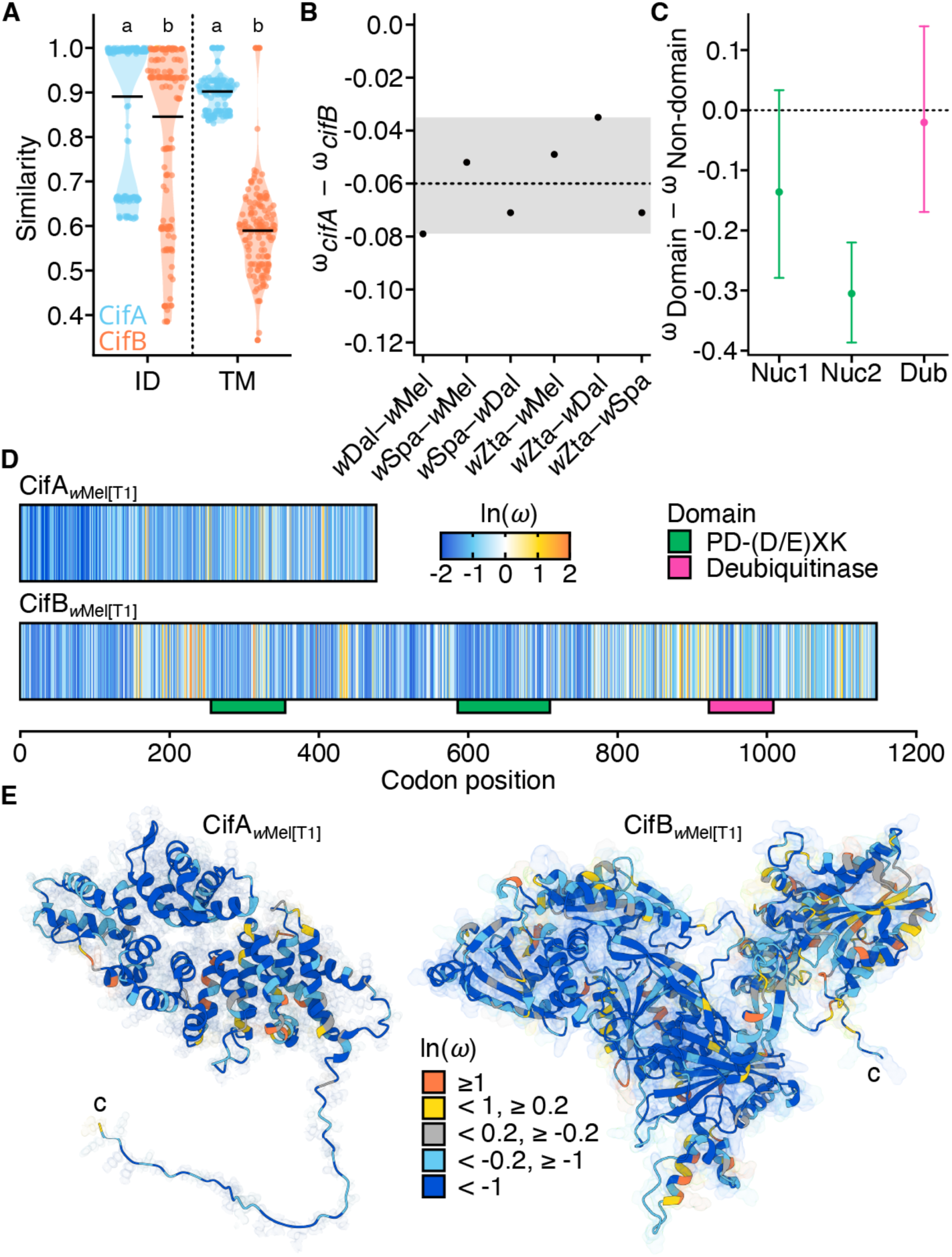
**Selection acts on intact *cifA*_[T1]_ and *cifB*_[T1]_**. **(A)** Putatively intact CifA_[T1]_ proteins are more similar than intact CifB_[T1]_ from the same pairs (*N* = 18) in both sequence identity (ID) and structural similarity (TM). **(B)** Pairwise median differences between *cifA*_[T1]_ and *cifB*_[T1]_ *ω* are consistently negative, with statistically significant *ω_cifA_* < *ω_cifB_* across all contrasts (*P* = 0.02). The gray bar denotes the 95% CI for the Hodges-Lehmann estimator of the median of paired differences. **(C)** Differences in mean per-site posterior *ω* between the three *cifB*_[T1]_ domains and pooled non-domain regions. The difference between one PD-(D/E)XK nuclease domain (Nuc2) and pooled *cifB*_[T1]_ non-domain regions was statistically significant (*P* < 0.0001) **(D)** Linear schematic of the per-site posterior *ω* estimates across CifA_[T1]_ and CifB_[T1]_. For CifB_[T1],_ colored boxes below the schematic denote HHPred-annotated functional domains as they occur along the codon alignment. **(E)** 3D schematic of the per-site posterior *ω* estimates mapped onto Cif*_w_*_Mel[T1]_ AlphaFold2 structures. “C” denotes the C-terminus of each protein. Per-site posterior *ω* estimates were obtained from the best-supported model for each gene and partition based on likelihood-ratio tests (Tables S12 & S13). As noted in our text, all sites identified by positive-selection models with BEB posterior *P* ≥ 0.95 show evidence of MNMs. Colors indicate ln(*ω*) values such that neutrality (*ω* = 1) maps to 0: dark blue (< –1), light blue (–1 ≤ x < –0.2), gray (–0.2 ≤ x ≤ 0.2), yellow (0.2 < x ≤ 1), and orange (> 1).

We evaluated six models in codeml that differ in their treatment of variation in *ω* across lineages and across sites (Fig. 6C). Prior to implementing these tree-based analyses, we tested for intralocus recombination using GARD, which found well-supported evidence for recombination in *cifA*_[T1]_ but not *cifB*_[T1]_ (Fig. S4; Supplemental Information). We found no evidence for selection pressures differing across lineages for either gene or *cifA*_[T1]_ partitions (Tables S12 & S13). The one-ratio model that assumes a single *ω* across all sites and lineages estimated an average *ω* = 0.39 for *cifB*_[T1]_, while estimates for upstream and downstream *cifA*_[T1]_ partitions (P_1_ and P_2_, respectively) differed from one another (P_1_ average *ω* = 0.22; P_2_ average *ω* = 0.38). This indicates that the constraint on *cifA*_[T1]_ is driven by relatively strong purifying selection on the N-terminus partition (sites 1–146). The N-terminus regions contribute to both CI and rescue (Shropshire *et al*. 2020b), while the C-terminus is important only for inducing CI (Shropshire *et al*. 2022).

We observed heterogeneity in *ω* across *cifB*_[T1]_ and *cifA*_[T1]_ partitions (Fig. 6 D & E; Tables S12 & S13; Supplemental Information), but models allowing a positively selected site class were supported only for *cifB*_[T1]_ and P_2_ of *cifA*_[T1]_. Among them, only *cifB*_[T1]_ sites 256, 258, and 341 were identified by both positive-selection models with Bayes Empirical Bayes (BEB) posterior *P* ≥ 0.95 (Table S14). Further analysis indicated that the site-specific *cifB*_[T1]_ results may be artifacts. Site 341 is located within the Nuc1 domain of *cifB*_[T1]_, while the other two sites are located in a 5’ non-domain region. We evaluated alignments of all three sites for evidence of multi-nucleotide mutations (MNMs) that violate assumptions of the codon substitution model (Venkat *et al*. 2018). We observed four unique codons at position 341: tgg (Trp) in *w*Mel, ttt (Phe), in *w*Dal, agg (Arg) in *w*Spa, and gga (Gly) in *w*Zta. All pairwise contrasts involve at least two nucleotide changes, with the exception of tggΗagg. All other BEB-supported sites have at least 2 contrasts that appear MNM-like.

Given that all analyses supported preservation of intact *cifB*_[T1]_, we summarized codon *ω* estimates by functional domain and compared them to the pooled non-domain background using label-permutation tests. Using the site-level posterior estimates for *ω* from the beta plus positive-selection model, we observed mean posterior *ω* < 1 for Nuc1 (0.55, 95% bootstrap interval: 0.42–0.71), Nuc2 (0.38, 95% bootstrap CI: 0.33–0.45), Dub (0.67, 95% bootstrap interval: 0.53–0.81), and for non-domain regions (0.69, 95% bootstrap interval: 0.63–0.75). Only the mean posterior difference involving Nuc2 was statistically significant (Δ = –0.31; 95% bootstrap interval: –0.39, –0.22; *P* = 0.0001; Fig. 6C). The observed difference for Nuc1 was also negative (Δ = –0.14), but with a relatively wide bootstrap interval that included 0 (95% bootstrap interval: –0.28, 0.03;, *P* = 0.06). In contrast, the mean posterior difference between the Dub and non-domain regions was small and statistically nonsignificant (Δ = –0.02; 95% bootstrap interval: –0.17, 0.14; *P* = 0.43), reflecting their similar mean posterior estimates. Siding-window analyses in 3D space broadly agreed, with the mean of the per-pair median differences on the log scale negative for all comparisons, but only statistically significant for Nuc2 (Δ*_p_* = −0.38; *P* = 0.003, Supplemental Information).

In summary, all analyses supported *cifA*_[T1]_ being more constrained than *cifB*_[T1]_, with constraint on *cifA*_[T1]_ driven by relatively strong purifying selection on the N-terminus regions involved in CI rescue (Shropshire *et al*. 2020b). However, our analyses indicated that all regions of intact *cifB*_[T1]_ are also constrained, with particularly strong purifying selection on the Nuc2 domain. We discuss these results in light of several theoretical predictions and the timescale of *cifB* degradation (Beckmann *et al*. 2021) and the loss of CI (Meany *et al*. 2019; Martinez *et al*. 2021).

## DISCUSSION

Decades of progress have increased our understanding of the timescale and processes of *Wolbachia* acquisition and the nature of *Wolbachia* genomic evolution. However, time estimates have remained uncertain because there have been few well-established examples of codiverging *Wolbachia* and hosts, combined with fossil-based estimates of host divergence times. Here we used recently available chronograms for filarial nematodes and complete sequences of their codiverging, obligately symbiotic *Wolbachia* to provide new calibrations and conservative estimates of their uncertainty. We also provide detailed analyses of *Wolbachia* genomic evolution, focusing separately on *cifs* that cause and rescue CI, the *Wovirus* prophages that contain *cifs*, and single-copy, non-phage *Wolbachia* genes. Our results demonstrate that these are temporally nested sets, with *Wolbachia* moving rapidly among hosts and *Wovirus* and *cifs* moving even more rapidly among *Wolbachia* genomes. Along their journey, efficient selection maintains *cifA*, while weaker stabilizing selection maintains intact *cifB.* Although the clade boundaries for the young *w*Mel-like and *w*Ri-like *Wolbachia* are arbitrary, and the range of plausible calibrations is wide, our central conclusions are robust to uncertainty in timing and incomplete sampling of these clades. We discuss our major findings.

### Timing the Mosaic and Bifurcating Evolution of Some *Wolbachia* Genomes

Ever since the discovery of intragenic recombination involving genes from distantly related *Wolbachia* (Werren and Bartos 2001; Jiggins *et al*. 2001; Baldo *et al*. 2006; Ellegaard *et al*. 2013; Wang *et al*. 2020) and the availability of complete *Wolbachia* genomes (including *w*Mel, Wu *et al*. 2004; *w*Pip, Klasson *et al*. 2008; and *w*Ri, Klasson *et al*. 2009), *Wolbachia* have been understood as “mosaics” because different portions of their genomes (and different portions of individual genes) follow discordant phylogenies. We here estimate the timing and genomic extent of that mosaicism, applying new divergence-time estimates from two distinct calibrations to four sets of *Wolbachia*: some closely related to *w*Ri (with conservative plausible range [CPR] for their divergence times: 14–218 KYA), some closely related to *w*Mel (CPR: 0.2–2.4 MYA), seven supergroup A *Wolbachia* spanning the divergence of *w*Mel and *w*Ri (CPR: 1.4–22 MYA), and the highly diverged *Wolbachia* from filarial nematodes that provide our calibrations (Table 7).

Within each of these clades, including the youngest, the *w*Ri-like variants, we document extensive movement of phages and *cifs* that occur within those phages. Hence, a significant fraction of these *Wolbachia* genomes are clearly “mosaic”. However, when we consider only single-copy, non-phage loci, we find no evidence of horizontal acquisition of distantly related alleles within the *w*Ri-like variants.

Increasing divergence time by an order of magnitude to our *w*Mel-like clade, we also observe no plausible evidence for horizontal allele acquisition. Zooming out another order of magnitude to seven supergroup A *Wolbachia* that span the divergence between *w*Mel and *w*Ri, we find a single example of horizontal acquisition involving an allele homologous to RefSeq_WP_010962975.1 in *w*Ha. While the *w*Ha allele differs substantially from the homologous alleles in the other six supergroup A *Wolbachia* (18.5–19.9%), it is identical to copies in *Wolbachia* from several diverse host species. Given that the divergence time between the *w*Mel-like and *w*Ri-like clades is more than twice as old as the crown age of either clade, our two-fold outlier criterion would have identified exchanges between them.

We do find some evidence of phylogenetic discordance from individual single-copy genes among these seven genomes spanning the divergence of *w*Mel and *w*Ri, but there is still a strong signal for a single topology. The sources of discordance could be ILS-like effects associated with short branches deep in the tree or model mis-specification leading to spurious inferences. Our data suggest that horizontal acquisition of distantly related alleles involves very few single-copy loci over 1–2 MYA. We conclude that *Wolbachia* genomes contain bifurcating genomic regions composed of single-copy loci, from which we can robustly estimate divergence times. Moreover, we find no evidence of recombination or horizontal acquisition of distantly related alleles for the supergroup C and D *Wolbachia* from filarial nematodes on which we base our calibrations.

Our results do not contradict previous evidence of extensive recombination involving relatively distantly related *Wolbachia* – because of the very different timescales involved. However, they do contradict the claim of Ellegaard *et al*. (2013) that extensive recombination among genomes within supergroups A and B produces highly discordant phylogenies from single-copy loci. Using only three variants within each supergroup and low stringency for node resolution (> 75% bootstrap support) in their single-gene phylogenetic analyses, Ellegaard *et al*. (2013) found nearly equal support for all three alternative rootings within each supergroup (see their Figure 3). Under purely bifurcating evolution, three taxa possess only one unrooted topology. Their rooting for the three variants within each supergroup is based on comparisons to distantly related outgroups from the other supergroup. Given the deep A–B split (Table 7), this rooting is unreliable because of saturation artifacts related to long branch attraction (see, for instance, Brinkmann *et al*. 2005). Empirical and simulation studies show that these “distant outgroup” artifacts can be resolved by analyzing more ingroup taxa and using more closely related outgroups (*cf*.

Rannala *et al*. 1998; Heath *et al*. 2008, p. 245). Our analyses do both. They indicate that the Ellegaard *et al*. (2013) results may reflect their inability to resolve the root position for each supergroup, rather than pervasive phylogenetic discordance involving single-copy genes within each supergroup.

Overall, we find little evidence of “mosaicism,” or phylogenic discordance, among the single-copy genes we examine within and between our relatively recently diverged *w*Mel-like and *w*Ri-like clusters. This contrast, between extensive evidence for recombination within single-copy loci between distantly related *Wolbachia* [*e.g.,* the Klasson *et al*. (2009) comparison of *w*Ri to distantly related supergroup A and B *Wolbachia*], and little evidence for such recombination between closely related variants, parallels the finding that although horizontal *Wolbachia* acquisition is clearly pervasive among distantly related hosts, it seems rare among individuals within specific host species (*e.g*., Richardson *et al*. 2012; Cooper *et al*. 2019), with some possible exceptions (*e.g*., Scholz *et al*. 2020). Rare horizontal transmission of *Wolbachia* within host species does not affect our divergence-time estimates of *Wolbachia* variants found in distinct host species – nor do rare examples of horizontal acquisition of single-copy *Wolbachia* loci.

Scholz *et al*. (2020) noted extreme variation in relative rates of mtDNA versus *Wolbachia* divergence, but much greater consistency between nuclear and *Wolbachia* divergence. Our nematode calibrations, which show only three-fold rate variation among distantly related *Wolbachia*, are consistent with this observation.

### Estimating *Wolbachia* divergence times

Our nematode-based *Wolbachia* rate calibrations, combined with new and existing genomic data, enabled us to revisit hypotheses concerning *Wolbachia* acquisition by *Nasonia* wasps and *Nomada* bees and to quantify the timescale of *Wolbachia* movements among distantly related hosts. Our analysis of the relatively young *w*Mel-like and *w*Ri-like *Wolbachia* clades, focused on the first *Wolbachia* identified in *Drosophila* species (Hoffmann *et al*. 1986; Hoffmann 1988), suggest that many young *Wolbachia* clades (*i.e*., ones that diverged on the order of 1 MYA or less) spread rapidly among anciently diverged hosts (that diverged on the order of 50–350 MYA). The host range and rapid movement of *w*Mel-like variants parallel those of *w*Ri-like variants (Turelli *et al*. 2018), suggesting that this scenario –– in which closely related *Wolbachia* invade anciently diverged hosts –– may characterize many common *Wolbachia*. Both CI phenotypes and *cif* complements are diverse among relatively recently diverged *Wolbachia*, with *Woviruses* and their associated *cifs* gained and lost on the order of 10^4^–10^6^ years.

An obvious question is whether our calibrations based on obligate C and D supergroup *Wolbachia* in nematodes are applicable to the facultative A and B supergroup *Wolbachia* in insects. The strongest support for this extrapolation comes from the fact that the range of point estimates for third-site substitution rates from nematodes, 6.47×10^-10^– 1.82×10^-9^, closely match the most plausible point estimates based on *Nomada* and *Nasonia*, namely 6.46×10^-10^ and 2.2×10^-9^, respectively. The *Nasonia* estimate rests on separate molecular clock estimates from insects and bacteria rather than fossil calibration. Obtaining more reliable point estimates and levels of uncertainty for *Wolbachia* substitution rates requires additional examples of *Wolbachia*-host codivergence, *i.e*., instances in which specific *Wolbachia* infections persist through host speciation. Such examples will come from increasingly reliable chronograms for insects, based on dated, amber-preserved fossils, and genome-based estimates of host and *Wolbachia* phylogenies and levels of sequence divergence, including mtDNA data. Our new filarial-nematode-based calibrations contribute to this goal. The rates estimated are significantly slower than the mtDNA-mutation-based estimate proposed by Richardson *et al*. (2012). Our chronograms are based on these new calibrations, but Table 5 compares our new time estimates to the shorter ones based on Richardson *et al*. (2012).

Three points are worth noting about these alternative calibrations. First, for eukaryotic nuclear genomes, some data indicate that substitution rates per year for synonymous substitutions and introns can closely match yearly mutation rate estimates. See, for instance, Table 1 of Begun *et al*. (2007) for sequence divergence along the terminal branches leading to *D. melanogaster* and *D. simulans*. The rates per year per site are quite comparable to the *D. melanogaster* mutation rate estimates of Keightley *et al*. (2009), assuming roughly ten generations per year and that *D. melanogaster* and *D. simulans* diverged about 4 MYA.) Second, as noted by Ho *et al*. (2005), mtDNA substitution rates slow dramatically over a timescale of tens of thousands of years. Finally, even if the faster Richardson *et al*. (2012) calibration is more accurate for the *w*Mel-like and *w*Ri-like variants we analyzed than our new calibrations, our key conclusions remain valid: namely, many closely related *Wolbachia* move rapidly between distantly related hosts; and over that rapid between-host-movement timescale, *Woviruses* and *cifs* within those *Wolbachia* turn over even more rapidly.

It is worth revisiting the Meany *et al*. (2019) estimate of supergroup A and B divergence with our new calibration. Using the Richardson *et al*. (2012) rate and both relaxed-clock [branch-rate priors Γ (2,2) and Γ (7,7)] and strict-clock analyses, Meany *et al*. (2019) estimated a plausible divergence interval for supergroups A and B of 6–64 MY (Meany *et al*. 2019). This range overlaps that of Werren *et al*. (1995), based on applying a “universal” bacterial synonymous substitution rate (Ochman and Wilson 1987) to the *ftsZ* locus (59–67 MYA). Both the Meany *et al*. (2019) and Werren *et al*. (1995) A-B divergence-time estimates were more recent than the Gerth and Bleidorn (2016) estimate of 76–460 MYA (point estimate: 217 MYA). However, both the Meany *et al*. (2019) and Werren *et al*. (1995) estimates now seem too recent, because the Richardson *et al*. (2012) calibration seems too fast and the *ftsZ* locus evolves slowly relative to most bacterial genes (Supplemental Information). Meany *et al*. (2019) questioned the Gerth and Bleidorn (2016) estimate for two reasons. First, their chronogram placed *w*No (from *D. simulans*) sister to all other supergroup B *Wolbachia*, which disagreed with their own phylogram and others (Lindsey *et al*. 2018). Second, their estimate of *w*Ri-*w*Suz divergence of 0.9 MY (with support interval 0.2 MY–2.2 MY) was nearly two orders of magnitude higher than the 11 KY estimate of Turelli *et al*. (2018). However, Turelli *et al*. (2018) used the Richardson *et al*. (2012) rate that we think underestimated *w*Ri-like *Wolbachia* divergence by about a factor of seven. Using our new calibrations, we now estimate *w*Ri-*w*Suz divergence as 76 KYA (CPR: 14–218 KYA) and supergroup A and B divergences as 152 MYA (CPR: 34–415 MYA) (Table 7). Our wide plausible ranges overlap those of Gerth and Bleidorn (2016); but as described below, we believe their point estimates are too large because their calibrations conflate cladogenic and horizontal *Wolbachia* acquisition.

Our analyses focus on the timescale over which clades of closely related *Wolbachia* have been acquired by distantly related hosts. In contrast, Bailly-Bechet *et al*. (2017) used a sample of about 1000 arthropod species (specifically OTUs characterized by mtDNA *CO1* sequences) from Pacific islands, to estimate how long, on average, specific *Wolbachia* infections persist in host lineages and how long uninfected host lineages remain uninfected. Bailly-Bechet *et al*. (2017) inferred that *Wolbachia* tend to persist in lineages for approximately seven million years. However, as Cooper *et al*. (2019) noted, this estimate is inconsistent with the fact that few sister species share codiverging *Wolbachia*. Because speciation times for *Drosophila*, and presumably many other insect clades, are on the order of one million years or less (Coyne and Orr 1997; Turelli *et al*. 2014), the infection-duration estimated by Bailly-Bechet *et al*. (2017) would lead to codivergence of *Wolbachia* and hosts for many sister pairs.

The estimated *Wolbachia* substitution rates, with conservative plausible ranges, from *Nomada* (Table 4) suggest a history of both cladogenic and horizontal *Wolbachia* acquisition in this four species clade. As shown in Table 4, the *Wolbachia* substitution rate based on comparisons of *N. ferruginata* and the three ingroup species agrees with that obtained from the *Brugia* nematode pair. In contrast, assuming cladogenic inheritance of *Wolbachia* throughout the four-species *Nomada* clade implies a greater than ten-fold slowdown of *Wolbachia* divergence between *N. flava* and *N. leucophtalma* (which diverged about 0.8 MYA, see Table 4) versus the outgroup *N. ferruginata* (from which they diverged about 2.1 MYA), while their nuclear and mtDNA continued to diverge at a rate typical for the four-species clade as a whole (see Table 3). Such a dramatic recent slowdown of *Wolbachia* divergence seems implausible given the relative constancy over a much longer timescale in the nematodes. We suggest that horizontal transmission of *Wolbachia* across the three-species ingroup is more plausible.

As noted in our discussion of Fig. 4B, individual genomes do not allow us to determine the direction or temporal order of *Wolbachia* acquisitions among hosts. Resolution is possible when multiple *Wolbachia* genomes are available from each host. If the *Wolbachia* haplotypes in species A are nested within the phylogeny of *Wolbachia* in species B, we can reasonably infer that Species A acquired *Wolbachia* from Species B. Conversely, if the closely related *Wolbachia* and mtDNA in host species A and B are reciprocally monophyletic sister clades with comparable divergence times, we can be reasonably confident of cladogenic *Wolbachia* acquisition. However, given the long time required to achieve reciprocal monophyletic gene trees – at least under neutrality (Hudson and Coyne 2002) – cladogenic *Wolbachia* transmission may often produce tangled trees in which at least one of the host species carries *Wolbachia* (and mtDNA) haplotypes that are paraphyletic with respect to those in a closely related host. The relative frequencies of these scenarios can be determined only with additional genomic analyses of the *Wolbachia*, mtDNA and nuclear genomes of closely related hosts.

Turelli *et al*. (2018) analyzed multiple *Wolbachia* sequences from several species, documenting rapid horizontal transfer of *w*Ri-like *Wolbachia* between divergent *Drosophila*. Specifically, the *w*Ri sequences from *D. simulans* formed a clade nested within a paraphyletic cluster of *w*Ana sequences from *D. ananassae*. Given that *D. ananassae* and *D. simulans* span the *D. melanogaster* species group, which diverged on the order of 47 MYA (see Fig. 1B), introgression is impossible. Hence we conjectured that horizontal non-sexual transmission of *Wolbachia* occurred between *D. ananassae* and *D. simulans*, with *D. ananassae* as a plausible source. Turelli *et al*. (2018, Fig. 1) also analyzed eight *Wolbachia* sequences from *D. suzukii* (*w*Suz) and one *D. subpulchrella* (*w*Spc). Two copies of *w*Suz from Asia (the other *w*Suz samples came from North and South America) formed a clade with the single *w*Spc, sister to the other six *w*Suz sequences. We used these data to conjecture that *D. suzukii* was a source of the *Wolbachia* in *D. subpulchrella*. However, the two putative Asian *w*Suz sequences, obtained from GenBank, were in fact derived from *D. subpulchrella* specimens (China-AA27 and Korea-AA7). This misidentification was first kindly pointed out to us by Mathieu Gaultier, then confirmed by Joanna Chiu. With this correction, the data in Fig. 1 of Turelli *et al*. (2018) produce sister clades of six *w*Suz and three *w*Spc, providing no information on the direction of transfer (Supplemental Information).

### Estimating the timescale of *Wovirus* and *cif* movements

The rapidly host-switching *Wolbachia* in our study differ widely in their CI phenotypes, from variants that do not cause CI to others that cause relatively weak or intense CI (Fig. 5A; Table S8). This includes *w*Mel-like *w*Seg and *w*Bocq that we confirmed cause relatively intense CI (Supplemental Information). CI is critical for *Wolbachia* spread within host species (Hoffmann *et al*. 1990; Hoffmann and Cooper 2024), as observed for *w*Ri that rapidly spread through global *D. simulans* populations to relatively high and stable frequencies (Hoffmann *et al*. 1986; Hoffmann *et al*. 1990; Turelli and Hoffmann 1991; Turelli and Hoffmann 1995; Kriesner *et al*. 2013). CI may also increase the likelihood of interspecific *Wolbachia* transfers, like those we have observed, through a process of clade selection (Hurst and McVean 1996; Turelli *et al*. 2022). Unsurprisingly, the effectors of CI have been of intense interest since the discovery that *Wolbachia* cause CI in *C. pipiens* (Yen and Barr 1971; Yen and Barr 1973). The relatively recent discovery of *cifs* (LePage *et al*. 2017; Beckmann *et al*. 2017; Shropshire *et al*. 2018) facilitated genomic analyses that have recently characterized *cif* diversity (*e.g.,* Lindsey *et al*. 2018; Turelli *et al*. 2018; Martinez *et al*. 2021; Vancaester and Blaxter 2023; Amoros *et al*. 2025a). It is well established that *Wovirus* and *cif* diversity reflect extensive horizontal transfers within and among *Wolbachia* lineages (*e.g.,* Masui *et al*. 2000; Bordenstein and Wernegreen 2004; Kent *et al*. 2011; Ellegaard *et al*. 2013; LePage *et al*. 2017; Cooper *et al*. 2019; Madhav *et al*. 2020; Martinez *et al*. 2021; Bordenstein and Bordenstein 2022; Vancaester and Blaxter 2023; Amoros *et al*. 2025a). What has remained unresolved is the timescale over which these exchanges occur. We estimate that *Wovirus* and *cif* exchanges involving *w*Mel-like and *w*Ri-like *Wolbachia* occur on the order of 10^4^–10^6^ years, excluding more rapid changes plausibly involving duplications. These bounds are conservative and derive from our nematode-based divergence estimates of bifurcating genomic regions and comparisons of *cifs* observed in closely related *Wolbachia*. High quality *Wovirus* assemblies from additional *Wolbachia* genomes are required to further resolve the timing of these gains and losses.

The most plausible mechanisms of rapid *cif* exchanges include natural transformation and movements involving *Wovirus* (Bordenstein and Wernegreen 2004; Gao *et al*. 2022; Vancaester and Blaxter 2023), plasmids (Martinez *et al*. 2022), and transposons (Gillespie *et al*. 2018; Cooper *et al*. 2019). While characterization of *Wolbachia* plasmids remains limited (*e.g.,* Reveillaud *et al*. 2019), IS elements are abundant and transcriptionally active in *Wolbachia* genomes (Duron *et al*. 2005; Iturbe-Ormaetxe *et al*. 2005; Riegler *et al*. 2005; Cordaux 2008; Klasson *et al*. 2009; Ellegaard *et al*. 2013; Cerveau *et al*. 2015), and frequently occur near *cifs* (Table S15). IS-mediated transfer was first identified as a plausible mechanism involved in the acquisition of *cif*_[T4]_ loci by *Wovirus* observed in *w*Yak and closely related genomes (Cooper *et al*. 2019) (Fig. 5A; Supplemental Information). Evidence for IS involvement in *cif* transfer has subsequently been documented in other *Wolbachia* (Madhav *et al*. 2020) and in diverse *Orientia tsutsugamushi* (Oswalt *et al*. 2025). These exchanges resemble cases where IS element-based composite and other compound transposons (Berg and Howe 1989) are responsible for moving antibiotic resistance genes among bacterial species (Razavi *et al*. 2020; Siguier *et al*. Chandler 2014). *cif* exchanges mediated by IS elements most likely occur when *Wolbachia* co-occur and actively replicate in the germline, enabling IS elements to mobilize *cif* modules and generate short extrachromosomal DNA fragments, which can subsequently integrate via recombination or other end-joining pathways in the recipient. Other transposons have been implicated in horizontal *cif* transfers, including PDDEXK2-family transposases that occur near *cif* loci (Table S15) in predicted transposon-like modules (Tan *et al*. 2024; Amoros *et al*. 2025a). These transposases cluster phylogenetically with *cifs* that span eight bacterial genera and show a history of horizontal transfer (Amoros *et al*. 2025b). Hence, transposons along with well-documented contributions of *Wovirus* transfers and recombination, and potentially other mechanisms, contribute to the rapid *cif* turnover we document. Determining the timescale of *cif* exchanges involving other symbionts (*e.g.,* Gillespie *et al*. 2018; Amoros *et al*. 2025b) will require estimating their divergence times using reliable calibrations.

### Estimating natural selection on *cifs*, and assessing the maintenance of CI

Our analyses of selection on *cif* loci is based on comparing synonymous versus non-synonymous substitutions for intact *cif*_[T1]_ pairs across *Wolbachia* found in different hosts. We find that intact *cifA*_[T1]_ is more constrained than intact *cifB*_[T1]_ (Fig. 6; Supplemental Information). Our analyses support relatively stronger purifying selection on the 5’ region of *cifA*_[T1]_ (Fig. 6D & E). Mutagenesis of CifA proteins previously found that N-terminus regions contribute to both CI induction and rescue (Shropshire *et al*. 2020b), while the C-terminus is important only for inducing CI. This suggests that selection to maintain functional *cifA*_[T1_ is related to CI rescue. We also find that weaker stabilizing selection maintains *cifB*_[T1]_ involved in producing CI (Fig. 6D & E). Constraint on intact *cifB*_[T1]_ is driven in part by relatively strong preservation of its Nuc2 domain relative to pooled non-domain regions (Fig. 6C). In contrast, we estimate similar *ω* for Nuc1, Dub, and pooled non-domain regions that are each also subject to weaker stabilizing selection. Overall, we estimate stronger selection to maintain CI rescue than to maintain CI induction.

How do these empirical patterns of selection compare to theoretical predictions? It is critical to recognize that different evolutionary predictions concerning CI induction and CI rescue emerge from the analyses of selection among *Wolbachia* variants within a specific host species versus “clade selection” analyses that consider the differential proliferation of *Wolbachia* variants across host species. Theory predicts that for *Wolbachia* lineages within an individual host species, natural selection acts to maintain resistance to CI, but it does not act to maintain the ability to induce CI against uninfected hosts (Prout 1994; Turelli 1994). In contrast, there is direct selection for *Wolbachia* to enhance host fitness (Prout 1994; Turelli 1994). These theoretical predictions are robust to population subdivision within a host species (Haygood and Turelli 2009). Even with extreme population subdivision, *Wolbachia* variants that greatly increase the intensity of CI will be selected against if they induce even very small costs on host fitness. In contrast, when considering which *Wolbachia* variants are most likely to spread among host species, an inherently slower “clade selection” process (Lewontin 1970), those that induce CI are favored. For *Wolbachia* that initially decrease host fitness, this is explained in terms relative chances of establishment in new hosts (Hurst and McVean 1996). For *Wolbachia* that enter new hosts as mutualists, CI is favored because it produces higher infection frequencies and persistence times within host species (Turelli *et al*. 2022). Selection within host species always acts to maintain CI rescue; whereas selection to induce CI is driven primarily by weaker “clade selection.” These predictions are broadly consistent with evidence for stronger selective constraints on *cifA*_[T1]_ than intact *cifB*_[T1]_.

The contributions of specific *cifB*_[T1]_ domains to CI induction are unresolved. The CifB Dub includes the only known catalytic Cif motif, and mutating its conserved cysteine active site ablates CI (Beckmann *et al*. 2017), although it does not consistently ablate yeast toxicity (Terretaz *et al*. 2023). Naturally occurring single-nucleotide variants in the Dub can also alter CI intensity, as observed for a valine-to-leucine variant in *w*Yak *cifB*_[T1]_. This single carbon change reduces in vitro deubiquitylating efficiency and also the intensity of CI when expressed transgenically in *D. melanogaster* (Beckmann *et al*. 2021), potentially underlying observed differences in *w*Yak and *w*Mel CI intensities (Hoffmann 1988; Cooper *et al*. 2017; Beckmann *et al*. 2021). While the CifB_[T1]_ Nuc1 and Nuc2 PDDEXK homologs lack canonical PD-(D/E)XK catalytic residues (Beckmann *et al*. 2017; Oladipupo and Hochstrasser 2025), mutating conserved sites in the *w*Mel copy ablated CI (Shropshire *et al*. 2020; Kaur *et al*. 2024), and this superfamily exhibits substantial functional diversity, with some members retaining nuclease activity despite non-canonical motifs (Knizewski et al. 2007). Our results support that these *cifB*_[T1]_ domains (and non-domain regions) are preserved in intact *cifB*_[T1]_ copies, with particularly strong constraint on Nuc2.

Despite this preservation of intact *cifB*_[T1]_ copies, putative loss-of-function mutations are known to be more common and to occur faster in *cifB* relative to *cifA* (Martinez *et al*. 2021). This includes observations of the loss of CI by *w*Mau in *D. mauritiana* via *cifB* degradation and putative preservation of *cifA* (Meany *et al*. 2019).

Clade selection, as discussed above, can explain our observation of selective constraint on *cifB*_[T1]_. Our analysis is restricted to putatively intact *cifB*_[T1]_ copies, and most loss-of-function mutations in *cifB* have occurred relatively recently, tending to occur on terminal gene-tree branches (Martinez *et al*. 2021). It is also clear that CifB contributes to other functions besides CI induction, and such pleiotropy could contribute to the selective constraint we observe (Turelli 1994). For example, Cifs are known to interact with host chromatin (Terretaz *et al*. 2023), and CifB seems to modulate *w*Mel interactions with host autophagy to maintain *w*Mel density in certain tissues (Deehan *et al*. 2021; Strunov *et al*. 2022). Given that *Wolbachia* densities in host tissues influence diverse host phenotypes and *Wolbachia* transmission rates (Bordenstein *et al*. 2006; Martinez *et al*. 2014; Chrostek and Teixeira 2015; Hague *et al*. 2022; Porter and Sullivan 2023; Radousky *et al*. 2023; Hague *et al*. 2024), modulation of host autophagy could significantly affect host and *Wolbachia* fitness. Regardless of whether the CifB Nuc2 domain functions enzymatically or through chromatin binding, the selection we observe is consistent with significant functional constraint. Those functions may involve CI, pleiotropic effects (*e.g.,* involving host autophagy), or both. Future work focused on pleiotropic effects of CifB and its domains will contribute to our understanding of the preservation of intact *cifB*_[T1]_, which nevertheless seems destined to degrade through evolutionary time faster than *cifA*_[T1]_. Similarly, phylogenetic analyses of the persistence of the CI phenotype within *Wolbachia* lineages, such as our *w*Ri-like and *w*Mel-like clades, can quantify the relative persistence times of *Wolbachia* lineages that do or do not cause CI.

### Conclusions and Implications for *Wolbachia* biocontrol

While the specific ecological mechanisms underlying rapid *Wolbachia* host switching in these closely related clades are unresolved, species interactions – and especially those involving parasitoid wasps (Vavre *et al*. 1999; Huigens *et al*. 2004; Gehrer and Vorburger 2012; Ahmed *et al*. 2015) and plant hosts (Caspi-Fluger *et al*. 2012; Chrostek *et al*. 2017; Li *et al*. 2017; Shi *et al*. 2024) – are plausible mechanisms underlying horizontal endosymbiont transfer (Cordaux *et al*. 2001; Kittayapong *et al*. 2003; Sintupachee *et al*. 2006; Jaenike *et al*. 2007; Oliver *et al*. 2010; Hurst *et al*. 2012; Hoffmann and Cooper 2024). Regardless of the mechanisms, pervasive and rapid *Wolbachia* host switching helps explain why *Wolbachia* are the most common invertebrate endosymbionts known and provides the physical opportunity for genomic exchanges between *Wolbachia*. These genomic exchanges also occur rapidly, with unique *cifs* sometimes observed in *Wolbachia* that diverged as recently as 10^3^–10^4^ years, with evidence mounting for transposon involvement in these rapid transfers. These movements produce unique *cif* complements among closely related *w*Mel-like and *w*Ri-like variants and other *Wolbachia* that contribute to observed variation in CI intensity. Because CI directly contributes to *Wolbachia* spread within host populations (Hoffmann *et al*. 1990; Turelli and Hoffmann 1991), and may contribute to the likelihood of *Wolbachia* movements between host species (Hurst and McVean 1996; Turelli *et al*. 2022), understanding the phylogenetic distributions of CI and the loci that underlie it remains critical.

CI enables *Wolbachia*-based suppression of vector and pest populations and the alteration of natural vector populations with pathogen-blocking *Wolbachia* transinfections. When *w*Mel *Wolbachia* are moved from *D. melanogaster* to *Aedes aegypti* – species that diverged about 250 MYA – the transinfection induces strong CI that can drive the endosymbiont to high frequencies (Walker *et al*. 2011; Hoffmann *et al*. 2011). *w*Mel introductions into *Ae. aegypti* have been very effective in locations where they have spread to high, stable frequencies (Hoffmann *et al*. 2011; Utarini *et al*. 2021; Lenharo 2023; Velez *et al*. 2023), with one study reporting an 86% reduction in dengue in an Indonesian city (Utarini *et al*. 2021) and a reduction of greater than 90% in two Colombian cites (Velez *et al*. 2023; Lenharo 2023). However, *w*Mel has been lost in some release locations (Hien *et al*. 2021; Moledo Gesto *et al*. 2021), particularly under very hot conditions. Temperature can affect CI intensity (Reynolds and Hoffmann 2002; Ross *et al*. 2019; Shropshire *et al*. 2021b) and the fidelity of maternal *Wolbachia* transmission (Ross *et al*. 2017; Hague *et al*. 2022; Hague *et al*. 2024). Identifying strong-CI-causing and virus-blocking *Wolbachia* from the tropics may facilitate *Wolbachia* biocontrol, and *w*Mel-like variants that naturally associate with tropical host species are obvious candidates for new applications (Gu *et al*. 2022; Gong *et al*. 2023; Thia *et al*. 2025). Notably, *w*Mel-like *Wolbachia* with diverse *cif* profiles have naturally colonized ecologically and geographically diverse hosts that are approximately 100 MY more diverged than are donor *D. melanogaster* and recipient *Ae. aegypti* dipterans (MRCA about 250 MYA, Wiegmann *et al*. 2011). We report a tropical *Wolbachia* variant (*w*Zts) that to our knowledge is the closest relative to *w*Mel in *D. melanogaster* (Supplemental Information). While *w*Zts CI has not yet been tested, all six *Z. tscasi* sampled in nature carry *w*Zts (producing 0.61 as the 95% lower bound for *w*Zts frequency), consistent with strong CI. Identification of *w*Zts and other variants closely related to pathogen-blocking *w*Mel has the potential to add versatility and contributes towards the development of more customized and environment-specific *Wolbachia* applications (Hoffmann and Cooper 2025).

## Materials and Methods

### Host and *Wolbachia* genomes

The sources for the genomes and their hosts are given in Table S1. For the *Wolbachia* that overlap with those in the study of Vancaester and Blaxter (2023), we used the same genome (*N* = 22), an alternative complete genome (*N* = 2), or a higher quality genome (*N* = 3) (Table S1). The latter were the circularized *w*Tei and *w*Yak genomes produced by Baião *et al*. (2021), and a *w*Dal genome from Scholz *et al*. (2020) that provided a higher quality assembly. We also include new genomes (*N* = 10) (Table S3). Table S1 also presents the sources for the draft nuclear, mitochondrial and *Wolbachia* genomes for *Nomada* bees, (*N. ferruginata*, (*N. panzeri*, (*N. flava*, *N. leucophthalma*))), and two trios of filarial nematodes – ((*O. volvulus, O. ochengi)*, *D. immitis*) and ((*B. malayi, B. pahangi*), *W. bancrofti*) – used for calibration. Data from *Nasonia* wasps, *N. longicornis* and *N. giraulti* include nine single-copy nuclear regions (Supporting Table 2 in Raychoudhury *et al*. 2009); and two mitochondrial genomic fragments: a 1048bp fragment of *CO1* and a ∼1200 bp fragment that includes complete *ATP6* and *ATP8* gene sequences (presented in Oliveira *et al*. 2008). For the *Nasonia* host nuclear genomes, we estimated divergence using loci specified below; for their mitochondria, we used data from all 13 protein coding loci.

### Estimation of phylogenies and chronograms

Our estimates of all phylogenies and chronograms were obtained using RevBayes 1.1.1 (Höhna *et al*. 2016), and used only protein-coding loci. We describe our general methods here, whereas we report the data used for specific analyses in the following sections. For host nuclear genes, we partitioned by locus and codon position. For *Wolbachia* and host mtDNA, we partitioned only by codon position, with the exception of *Wolbachia* single-gene trees, which were unpartitioned. Phylograms were estimated using the substitution model GTR + I + Γ(7,7) and chronograms were estimated using GTR + Γ(7,7). We did not assess the relative adequacy of simpler substitution models, because the results of Abadi *et al*. (2019) suggest that results from GTR + Γ are likely to be robust. (Analyses with a strict clock produced smaller divergence-time estimates, and more relaxed clocks, *e.g*., Γ(2,2), lose node resolution.)

We estimated relaxed-clock relative chronograms with the root age fixed to 1 and used the same birth-death prior as Turelli *et al*. (2018). Each partition had an independent rate multiplier with prior Γ(1,1), as well as stationary frequencies and exchangeability rates drawn from flat, symmetrical Dirichlet distributions. The branch-rate prior was Γ(7,7), normalized to a mean of 1 across all branches.

We ran phylogram analyses for 100,000 RevBayes generations and chronograms for 200,000 generations. Unlike MrBayes (Ronquist *et al*. 2012), in which “one generation” corresponds to a single proposed move in parameter space, one generation in RevBayes involves many proposals to accommodate model complexity (Höhna *et al*. 2017). For each analysis, we completed four independent RevBayes runs. Convergence for each run was assessed using Tracer (Rambaut *et al*. 2018). In all our analyses, the four runs converged to the same result. Nodes with posterior probability less than 0.95 were collapsed into polytomies.

### Phylogenies and chronograms for hosts of *w*Mel-like *Wolbachia*

Our estimates concerning host divergence rely on recent calibrations from the literature, as discussed in our Results. Our host species (presented in Tables 1 & S1) are extraordinarily diverse (see Results).

For the drosophilids in Table 1, node ages and approximate support intervals presented in Fig. 1B and Table 2 were estimated from the fossil-calibrated chronogram of Suvorov *et al*. (2022), using supplementary information as needed. For drosophilid species in Table S1 that are not explicitly provided in Fig. 1 of Suvorov *et al*. (2022), we used the NCBI Taxonomy Browser to determine the closest relative(s) included in Suvorov *et al*. (2022) (Table S1). The node ages and approximate support intervals not explicitly shown in their Fig. 1 were kindly provided by A. Suvorov (pers. comm.).

We used the Suvorov *et al*. (2022) calibrations for host divergence when possible. For species not included in Suvorov *et al*. (2022), we estimated divergence with a relative chronogram including the species pair in question and a node present in Suvorov *et al*. (2022). Specifically, to estimate the *D. borealis*-*D. incompta* divergence time, we estimated a relative chronogram for {*D. hydei*, *D. virilis*, *D. borealis*, *D. incompta*}, for which *D. hydei* is the outgroup and an age estimate for the root is available from Suvorov *et al*. (2022). Similarly, to estimate the *D. malagassya-D. seguyi* and *D. sp. aff. chauvacae-D. bocqueti* divergences, we estimated a relative chronogram for (*D. jambulina*, ((*D. bocqueti*, *D. aff chauvacae*), (*D. malagassya*, *D. seguyi*))) with an age estimate for the root from Suvorov *et al*. (2022). For these chronograms, we estimated relative divergence using relaxed-clock relative chronograms under the GTR + Γ(7,7) model of molecular evolution, following the methods in Conner *et al*. (2021), except that we partitioned by both gene and codon position. For these trees, we used the coding sequences for the 20 nuclear genes used in the analyses of Turelli *et al*. (2018): *aconitase, aldolase, bicoid, ebony, Enolase, esc, g6pdh, GlyP, GlyS, ninaE, pepck, Pgi, Pgm1, pic, ptc, Tpi, Transaldolase, white, wingless,* and *yellow*. Coding sequences for these genes in *D. melanogaster* were obtained from FlyBase. We used tBLASTn to identify homologs in the other genome assemblies. The sequences were aligned with MAFFT v. 7 (Katoh and Standley 2013) and trimmed of introns using the *D. melanogaster* sequences as a guide. The resulting sequences were visually inspected for gaps, which only occurred in multiples of three and did not create frameshifts.

Estimation of mtDNA divergence times requires a plausible range of mtDNA substitution rates.

Because these rates may decline with species divergence (Ho *et al*. 2005), we computed mtDNA third-position substitution rates for five *Drosophila* species pairs that differ in their estimated divergence times (*Drosophila* 12 Genomes Consortium; Suvorov *et al*. 2022): *D. melanogaster*-*D. simulans* (3.62 MY, 95% CI: 2.92–4.4 MY), *D. erecta*-*D. yakuba* (5.47 MY, 95% CI: 4.59–6.38 MY), *D. mojavensis*-*D*.

*virilis* (24.34 MY, 95% CI: 21.94–26.88 MY), *D. pseudoobscura*-*D. suzukii* (32.78 MY, 95% CI: 29.77–35.76 MY), and *D. grimshawi*-*D. willistoni* (46.84 MYA, 95% CI: 43.85–49.85 MY). For each of these 10 species, coding sequences for the 13 protein-coding mitochondrial genes in the inbred reference strains were obtained from FlyBase. They were aligned with MAFFT v.7 and concatenated. We counted nucleotide differences and applied the Jukes-Cantor correction for multiple substitutions. mtDNA third-position substitution rates were computed using the Jukes-Cantor estimates for the number of substitutions and fossil-calibrated divergence times estimates of Suvorov *et al*. (2022). Our possible examples of introgressive *Wolbachia* transfer involve relatively recently diverged pairs (see below). Thus, we averaged the point estimates for third-position mtDNA substitution rates for the two most recently diverged *Drosophila* pairs, *D. melanogaster*-*D. simulans* and *D. erecta*-*D. yakuba*. We used the fastest and slowest substitution-rate estimates across both sets of credible intervals to produce a conservative plausible range of mtDNA third-position substitution rates. We used these rate estimates and the observed mtDNA third-position differences between species pairs to estimate their mtDNA divergence times and plausible ranges.

### Chronograms for hosts used to calibrate *Wolbachia* divergence rates

***Nasonia*.** Raychoudhury *et al*. (2009) used a *Drosophila*-based calibration (Tamura *et al*. 2004) for synonymous divergence of host nuclear genes to estimate that *N. giraulti* and *N. longicornis* diverged about 0.51 MYA.

**Filarial nematodes.** We consider two trios of filarial nematodes, each with a pair of congeners and an outgroup containing a *Wolbachia* variant belonging to the same supergroup as those in the congeners: ((*O. volvulus, O. ochengi)*, *D. immitis*) and ((*B. malayi, B. pahangi*), *W. bancrofti*). For both trios, Qing *et al*. (2025) estimated the time to the MRCA of the three species. For each trio, we used their crown-age estimates and then interpolated internal node ages from multilocus-based relative chronograms. For each of these six nematode species, we obtained 33,099 bp of nuclear data (Table S1). Our host relative chronograms were estimated using data from the 20 single-copy nuclear loci: *aldolase, ddb, fdh, g6pdh, galactokinase, glyp, glys, idh, ldh, lias, mes, pgi, pgm, pisy, ptc, rpi, sqv, tag, transaldolase,* and *vem*.

These metabolic housekeeping genes exist as single-copy homologs in each of the six species. Homologs were aligned with MAFFT v. 7 (Katoh and Standley 2013). We visually inspected the alignments for gaps, which occurred only in multiples of three and produced no frameshifts in the alignments. In assessing our *Wolbachia* calibrations, we also analyzed host mtDNA to cross-calibrate divergence with the nuclear and *Wolbachia* loci.

### Nomada

We reanalyzed Gerth and Bleidorn’s (2016) proposed example of codivergence of *Wolbachia* within the four-species clade of *Nomada* bees, (*N. ferruginata*, (*N. panzeri*, (*N. flava*, *N. leucophthalma*))). Our analyses depend on comparing rates of divergence for the *Nomada* nuclear genomes, *Nomada* mtDNA, and their associated *Wolbachia* (Table S1). For the crown age of the four-species clade, we used the estimate and 95% credible interval, 2.42 (1.16–3.37 MY), obtained from Fig. 3 and p. 6 of Gerth and Bleidorn (2016). Internal node ages were estimated from relative chronograms derived from 36,402 bp of nuclear protein-coding sequence from 20 loci from each of the four species: *aconitase, arl2, beta-hexosaminidase, citrate synthase, cytidine deaminase, esc, g6pdh, galactokinase, glys, houki, idh, las, pic, pis, protein kinase JNK, rpi, sgl, shmt, transaldolase,* and *UBFD1*. The point estimates for the ages of the internal nodes were based on the crown age estimate of 2.42 MY. Support intervals for internal node ages are based on the 95% credible intervals estimated from the relative chronogram.

### *Wolbachia* assembly and phylogeny estimation

For the 10 genomes novel to this study, we trimmed the reads using Sickle v. 1.3 (Joshi and Fass 2011) and assembled them with ABySS v. 2.2.3 (Jackman *et al*. 2017) with Kmer values of 51, 61,…, 91. From these host assemblies, scaffolds with best nucleotide BLAST matches to known *Wolbachia* sequences, with E-values less than 10^-10^, were extracted as the *Wolbachia* assembly. For each host, the best *Wolbachia* assembly (fewest scaffolds and highest N50) was retained. To assess the quality of these draft assemblies, we used BUSCO v. 3.0.0 (Simão *et al*. 2015) to search for homologs of the near-universal, single-copy genes in the BUSCO proteobacteria database (https://busco.ezlab.org/). As controls, we performed the same search using the reference genomes for *w*Ri, *w*Au, *w*Mel, *w*Ha and *w*No.

To identify genes for the phylogenetic analyses, all our *w*Mel-like and *w*Ri-like *Wolbachia* genomes (Table S1) were annotated with Prokka 1.14.5 (Seemann 2014). In cases where genes in different genomes match the same reference gene according to Prokka, we considered them homologous. To avoid pseudogenes, paralogs and frameshifts, we used only genes present in a single copy in each genome and required that at most one draft genome had gaps in the alignment; the deletions all were in frame, involving sets of three base pairs. Genes were identified as single-copy if they uniquely matched a bacterial reference gene identified by Prokka. They were then aligned with MAFFT v. 7 (Katoh and Standley 2013). We estimated three separate phylogenies. For the set of the 20 *w*Mel-like genomes, 438 genes, a total of 409,848 bp, met these criteria. For the 20 *w*Mel-like genomes plus the 8 *w*Ri-likes, 346 genes, a total of 310,977 bp, met these criteria. For the 8 *w*Ri-like genomes above and here, we used the same data set used in Turelli *et al*. (2018), which includes 525 genes, a total of 506,307 bp. These “core” genes within the *w*Mel-like and *w*Ri-like clades show little evidence for intragenic recombination or horizontal acquisition (see below).

### Assessing recombination and the mosaic structure of closely related *Wolbachia* genomes

In the following subsections, we describe single-locus tests for recombination that search for discordance between phylogenies associated with different segments of individual genes. In addition to looking for recombination within coding loci, we consider a range of other tests to assess the accuracy of our phylograms and chronograms. We determine whether individual *Wolbachia* loci support phylogenies different from those estimated from concatenating data across loci and assess possible causes of discordance. We also determine the loci that show the highest levels of allelic variation across various sets of reference strains. These most-variable loci are plausible candidates for horizontal acquisition of alleles from outside the reference set. We examined patterns of pairwise differences among alleles, looking for outliers suggesting horizontal acquisition.

### Detecting recombination within genes among genomes

To test for intralocus recombination across *Wolbachia* genomes, we used a genetic algorithm for recombination detection (GARD) (Kosakovsky Pond *et al*. 2006), plus all three statistical methods implemented in PhiPack: the Pairwise Homoplasy Index (PHI), Maximum Chi Square (χ2), and Neighbor Similarity Score (NSS) tests (Bruen *et al*. 2006). The PHI test detects recombination by evaluating phylogenetic incompatibility between sites, χ2 divides the alignment into windows and assesses differences in the distribution of substitutions to identify potential breakpoints, and NSS measures similarity along the sequence to identify reductions indicative of a breakpoint. GARD is widely implemented, including in analyses of intralocus *Wolbachia* recombination (Wang *et al*. 2020). We focused our analyses on single-copy genes at least 300 bp in length, with minimum recombination segments of 100 bp. The 300 bp criterion excludes a small number of the single-copy genes within each set described above. We set GARD to detect a maximum of two potential breakpoints, and used the GTR + Γ model with four rate categories. For our 20 *w*Mel-like variants, 411 genes (403,206 bp) met our length criteria.

We used these same analyses to test for recombination in three additional sets of *Wolbachia* with different levels of sequence divergence. We analyzed the 8 *w*Ri-like *Wolbachia* presented in Fig. 1 in Turelli *et al*. (2018). 517 genes (506,307 bp) met our criteria. We also examined the seven more diverged supergroup A *Wolbachia* genomes introduced above: *w*Mel, *w*Au, *w*Ri, *w*Tri, *w*Ha, *w*Bic and *w*Bar. For this set, 401 genes (398,602 bp) met our criteria. Finally, we considered more anciently diverged *Wolbachia* in filarial nematodes relevant to our calibrations. We analyzed four supergroup D *Wolbachia* with the following phylogeny: (*w*Wb in *W. bancrofti*, (*w*Bm in *B. malayi, w*Bp in *B. pahangi*)) and the outgroup *Wolbachia* (*w*Ls) associated with *Litomosoides sigmodontis*. For these four, 356 genes (349,602 bp) met our criteria. For our three supergroup C *Wolbachia* trio (*w*Ov in *O. volvulus, w*Oo in *O. ochengi*, *w*Di in *D. immitis*), we could not find additional supergroup C *Wolbachia* outgroup. This precludes analysis of within-gene discordance for supergroup C because there is only one unrooted phylogeny for three taxa.

We assessed the reliability of the GARD recombination results for the seven supergroup A *Wolbachia* by visually inspecting conflicting trees inferred by GARD and the data underlying them. As discussed in our Results, GARD makes artifactual recombination calls when there is insufficient sequence variation to produce reliable phylograms (Supplemental Information).

We next describe several independent analyses that provide strong support for bifurcating evolution of major portions of the *Wolbachia* genome, including for the seven supergroup A *Wolbachia*.

### Discordance of gene trees

As an alternative assay for mosaic genome structure, we looked for variation among the unrooted topologies inferred from 442 single-copy protein-coding loci (total 413,115 bp) from the focal seven supergroup A *Wolbachia*. For each gene, we attempted to infer an unrooted tree for all seven *Wolbachia*. For illustration, we highlighted the subset of 52 loci that produced fully resolved trees, with posterior probabilities > 0.95 for each node, and compared them to the consensus topology estimated from the concatenated sequence data (52 genes; 72,027 bp). To assess discordance, we estimated an unrooted species tree from all 442 gene trees using weighted ASTRAL v1.23 (Zhang and Mirarab 2022), and used IQ-TREE v. 3.0.1 to calculate gene concordance factors (gCF) on the reference topology (Wong *et al*. 2025). Using 50%, 60%, 80%, and 95% bootstrap thresholds, we collapsed nodes with low support in the gene trees and compared results across thresholds. Default settings were used for ASTRAL and IQ-TREE analyses.

We applied the same discordance analysis to seven nematode-derived *Wolbachia* genomes, the four supergroup D genomes used for our intralocus recombination assay above and three supergroup C genomes used for our calibration. This tests for horizontal transmission of fragments across these genomes.

### Polymorphism and pairwise allele-difference assays for horizontal transmission

A limitation of gene-tree discordance analyses is that they require sufficient differences among the alleles to produce a fully resolved phylogeny. If only one genome horizontally acquired a distantly related allele (or allele fragment) at a specific locus, the remaining alleles may not be sufficiently different to provide full phylogenetic resolution. As an alternative assay for horizontal transmission, we ranked the single-copy loci by their levels of polymorphism, measured as the fraction of variable nucleotide sites, across our *w*Mel-like and *w*Ri-like *Wolbachia* and across our seven more diverse supergroup A *Wolbachia*. The most polymorphic loci are candidates for having horizontally acquired alleles. To account for variation in rates of molecular evolution across loci, we directly examined pairwise differences between all alleles, looking for outliers as candidates for horizontal acquisition.

Based on a preliminary examination of pairwise differences between alleles at single-copy loci, we applied a heuristic criterion for identifying plausible examples of horizontally acquired alleles. For individual loci within a specific set of *Wolbachia*, such as our *w*Mel-like and *w*Ri-like variants, we categorized an allele as a “two-fold outlier” if it differs from all other alleles in the set at twice as many nucleotides as any of the remaining alleles differ from one another. This heuristic criterion is obviously most easily satisfied for closely related *Wolbachia*. For loci with two-fold outliers, we examined the flanking loci and mapped the nucleotide differences within the locus to assess the size of the fragment that was plausibly horizontally acquired.

### Calibrating rates of *Wolbachia* divergence

We produce new calibrations based on codivergence of *Wolbachia* and filarial nematode hosts. Raychoudhury *et al*. (2009) pioneered codivergence calibrations. Before describing our new filarial nematode analyses and our reanalysis of data from *Nomada* bees, we describe our reanalysis of the *Nasonia* calibration from Raychoudhury *et al*. (2009).

### *Nasonia* calibration

To determine whether any of the 11 *Wolbachia* variants found in 4 *Nasonia* species were plausible candidates for cladogenic acquisition, Raychoudhury *et al*. (2009) compared molecular-clock-based estimates for host divergence times (described above) with molecular-clock-based divergence-time estimates for particular pairs of their *Wolbachia*. To produce a plausible divergence time for the *Wolbachia w*NgirB and *w*NlonB1, they used a synonymous bacterial divergence rate (double the substitution rate) of 9×10^-9^. This produces an age estimate of 0.41 MYA for the MRCA of *w*NgirB and *w*NlonB1, comparable to that for their hosts. This rough agreement based on two plausible molecular clocks is the key evidence supporting *Wolbachia*-host codivergence in these *Nasonia*. To understand the implications of this hypothesized *Nasonia-Wolbachia* codivergence for *Wolbachia* substitution rates, we averaged the eukaryotic-based and prokaryotic-based divergence-time estimates to approximate the age of the MRCA of both pairs (*N. giraulti*, *N. longicornis*) and (*w*NgirB, *w*NlonB1) as about 0.46 MYA. We reanalyzed the raw data available from Raychoudhury *et al*. (2009) for these host nuclear genomes and these two *Wolbachia* variants to produce the divergence-time estimates. We also compared the hosts’ mtDNA to compare with other examples of putative cladogenic *Wolbachia* acquisition.

### Filarial nematode calibrations

For each species of the two host trios – ((*O. volvulus, O. ochengi)*, *D. immitis*) and ((*B. malayi, B. pahangi*), *W. bancrofti*) – we extracted 598,257 bp from single-copy *Wolbachia* loci. To look at relative rates of *Wolbachia* and host mtDNA divergences, we also extracted the 13 protein-coding mtDNA genes, with a total of 10,361 bp. To estimate pairwise divergences, we aligned the genes, counted nucleotide differences and applied the Jukes-Cantor correction for multiple substitutions. When computing rates associated with each outgroup, we averaged the two sequence divergence estimates from each sister to the outgroup. We estimated rates and their support intervals using our point estimates and support intervals for host divergence times.

### *Nomada* calibrations

The new nematode data allow us to critically examine the putative codivergence of *Wolbachia* with hosts in the four-species clade of *Nomada* bees (*N. ferruginata,* (*N. panzeri,* (*N. flava, N. leucophtalma*))) (Gerth and Bleidorn 2016). To assess the implications and plausibility of *Wolbachia*-host codivergence throughout this clade, we compared rates of *Wolbachia* sequence divergence to the divergence of host mtDNA (using data kindly provided by M. Gerth and C. Bleidorn) and nuclear loci. Our analyses used the 13 protein-coding gene sequences, a total of 9249 bp. We analyzed 613,605 bp of *Wolbachia* protein-coding sequence from single-copy loci. *Wolbachia* substitution rates were estimated as for the nematodes and *Nasonia*. We estimated separate *Wolbachia* rates from each node of the *Nomada* tree.

### *Wolbachia* chronograms

We estimated chronograms for three datasets—*w*Mel-like clade, *w*Ri-like clade (Turelli *et al*. 2018), and both clades combined. We first estimated a relaxed-clock relative chronogram with the root age fixed to 1. There were not enough substitutions to partition by gene as well as codon position. In Turelli *et al*. (2018), we converted the relative chronogram into an absolute chronogram using the rate prior Γ(7,7)×6.87×10^-9^ substitutions per third-position site per year, derived from the mutation-based posterior distribution estimated by Richardson *et al*. (2012). The mean assumes 10 generations per year. Absolute branch lengths (times) were calculated as the relative branch length times the third position rate multiplier estimated from the substitutions per third-position site per year.

Here we consider alternative calibrations for the third-position rate, based on putative examples of cladogenic *Wolbachia* transmission in *Nasonia* and *Nomada*. Turelli *et al*. (2018) used the Γ distribution to approximate the variation in the mutation-rate estimate obtained by Richardson *et al*. (2012). Although Γ(7,7) approximated the per-generation variance estimate, Turelli *et al*. (2018) used it to approximate the per-year variance. Because Turelli *et al*. (2018) also assumed 10 generations per year, the variance per year was underestimated in this analysis. Here we use a different approach to approximating the uncertainty in substitutions per third-position site per year and explore the consequences of incorporating that uncertainty in our divergence-time estimates. We estimate plausible ranges of substitution rates for different subsets of *Nomada* host-*Wolbachia* comparisons based on estimated ranges for host divergence times (see Tables 2 and 3 below). The alternative calibrations of *Wolbachia* divergence rates produce a wide range of plausible divergence times, as illustrated below. The variation among estimates based on alternative calibrations probably better represents the uncertainty of divergence-time estimates than do the probabilistic support intervals produced associated by individual calibrations.

For their *Nasonia* analyses, Raychoudhury *et al*. (2009) used widely applied eukaryotic and bacterial molecular clock calibrations to separately estimate the divergence times for hosts *N. longicornis* and *N. giraulti* and the *Wolbachia* inferred to be cladogenically acquired, *w*NlonB1 and *w*NgirB. We averaged those estimated divergence times for hosts and their *Wolbachia* to produce a divergence-time estimate of 0.46 MY, which produces an average third-site *Wolbachia* substitution rate of 2.2×10^-9^ substitutions per third site per year over the 4,486 bp of their *Wolbachia* DNA sequence data (see Table 4 below).

For their *Nomada* pairs with plausible cladogenic *Wolbachia* transmission, Gerth and Bleidorn (2016) used fossil-based estimates of the host divergence times. Specifically, they estimated the crown age of the clade (*N. ferruginata,* (*N. panzeri,* (*N. flava, N. leucophtalma*))) as 2.42 MY, with 95% credible interval (1.16, 3.37) MY. As described above, we used additional single-copy nuclear data from these four species to more accurately estimate the relative ages of the internal nodes of this clade. We then converted the relative chronogram to an absolute chronogram by assuming that the crown age is 2.42 MY. We used the *Wolbachia* divergence rates between the hosts to estimate mean substitution rates.

Support intervals for the rates were approximated based on the support intervals for the host divergence times. The data-motivated models used are described in our Results. To illustrate the consequences of approximating uncertainty in the individual *Nomada* rate estimates, we compare divergence-time estimates and their support intervals produced by analyses assuming constant versus stochastically distributed divergence-rate priors. That uncertainty, associated with individual calibrations, is contrasted with the larger effects associated with alternative rate calibrations.

### CI phenotyping

While estimates of CI intensity exist for many of the *Wolbachia*-host pairs in our study, we tested for CI in two previously uncharacterized *D. montium* subgroup host species that carry *w*Mel-like variants: *w*Seg in *D. seguyi* and *w*Bocq in *D. bocqueti.* We secured *D. seguyi* and *D. bocqueti* stocks carrying *w*Mel-like *Wolbachia* from the National *Drosophila* Species Stock Center. To generate lines of both species without *Wolbachia*, we exposed *Wolbachia*-carrying lines to tetracycline-supplemented (0.03%) cornmeal medium (see Cooper and Shropshire 2024a for details) for three generations. We initially confirmed the absence of *Wolbachia* in the treated flies using PCR within two generations of tetracycline treatment using primers for the *Wolbachia*-specific *wsp* gene (Braig *et al*. 1998; Baldo *et al*. 2005) and a second reaction for the arthropod-specific 28S rDNA as a host control (Nice *et al*. 2009; Cooper and Shropshire 2024b). Our PCR thermal profile began with 3 min at 94C, followed by 34 rounds of 30 sec at 94C, 30 sec at 55C, and 1 min and 15 sec at 72C. The profile finished with one round of 8 min at 72C. We visualized PCR products using 1% agarose gels that included a molecular weight ladder. The stocks were maintained and experiments were conducted in an incubator at 25C. Tetracycline-treated stocks were given at least four generations to recover prior to our CI experiments, and subsequent PCR-based tests confirmed that these stocks remained *Wolbachia*-free. All stocks were maintained under standard conditions using established protocols (Wheeler *et al*. 2024; Hartman *et al*. 2024).

We reciprocally crossed *Wolbachia*-carrying *D. seguyi* and *D. bocqueti* lines to their tetracycline-treated conspecifics and estimated the egg hatch frequencies from potentially incompatible crosses between females without *Wolbachia* and males with *Wolbachia* (denoted IC) and from the reciprocal compatible cross (CC) between females with and males without *Wolbachia*. With cytoplasmic incompatibility, we expect lower egg hatch from IC crosses than from CC crosses. Virgins were collected from each line and placed into holding vials for 48 hr. We set up each IC and CC cross with one female and one male in a vial containing a small spoon with cornmeal medium and yeast paste for 24 hr (Shropshire 2025). Males and females were two days old at the beginning of these experiments. Each pair was transferred to a fresh vial every 24 hr for 5 days. We counted the number of eggs laid at the time the adults were transferred to new vials and the number of eggs that hatched were scored after an additional 24 hrs. The data analyzed were hatch proportions for crosses across the 5-day period. To control for cases where females may not have been inseminated, we excluded crosses that produced fewer than 10 eggs.

We used one-sided Wilcoxon tests to determine whether IC crosses produce lower egg hatch proportions than do CC crosses.

### Extracting sr alleles to type *Wovirus*

We used sr alleles to type *Woviruses* in *w*Mel-like and *w*Ri-like genomes, focusing on *Woviruses* (*i.e.,* sr1WO-sr3WO) known to contain *cifs* (Bordenstein and Bordenstein 2022). sr genes assist with phage genome integration into and excision from the bacterial chromosome. Their phylogeny corresponds to module synteny and genome integration, making sr alleles a useful *Wovirus* typing tool (*sensu* Bordenstein and Bordenstein 2022). Across all *w*Mel-like and *w*Ri-like *Wolbachia*, we used WOCauB3, WOVitA1, WOMelB, and WOFol2 sr alleles as queries in BLAST searches to characterize sr1WO, sr2WO, sr3WO, and sr4WO *Wovirus*, respectively. If the *Wolbachia* assembly clearly assigned an sr sequence to a phage, we typed the phage (sr1WO, sr2WO, sr3WO, or sr4WO). For all of these cases, sr sequences in the *Wolbachia* assemblies mapped to only one of the four query sequences.

### Extracting *cif* sequences

For our set of *w*Mel-like and *w*Ri-like *Wolbachia*, we used BLAST to identify contigs with *cif* sequences with E-values less than 10^-10^. Our search used the following reference genes to query and categorize *cif* sequences: Type 1 from *w*Mel (*cif_w_*_Mel[T1]_), Type 2 from *w*Ri *D. simulans* (*cif_w_*_Ri[T2]_), Type 3 from *w*No in *D. simulans* (*cif_w_*_No[T3]_), Type 4 from *w*Pip *C. pipiens* (*cif_w_*_Pip[T4]_), Type 5 from *w*Stri in *Laodelphax striatellus* (*cif_w_*_Stri[T5]_), Type 5 from *w*Tri *D. triauraria* (*cif_w_*_Tri[T5]_), and Type 5 from *w*Bor *D. borealis* (*cif_w_*_Bor[T5]_). These *cifs* are also classified based on predicted enzymatic domains of the *cifB* locus involved in CI induction (Oladipupo and Hochstrasser 2025). Type 1 *cifs* are referred to as *cid* loci based on a C-terminal deubiquitylase domain, Types 2-4 are referred to as *cin* loci based on predicted nuclease domains, and Type 5 *cifs* are referred to as *cnd* loci based on predicted nuclease and deubiquitylase domains (Beckmann *et al*. 2017; Chen *et al*. 2020). We use “Type” classification here, while acknowledging alternative proposals for naming of these loci (Shropshire *et al*. 2019; Beckmann *et al*. 2019).

We assigned the *cif* sequences to Types based on similarity to the reference genes noted above. We describe specific *cifs* using up to three pieces of information in a subscript behind the gene name: strain, Type designation, and copy number. For example, *w*Ri encodes two T1 *cifA* variants that we denote *cifA_w_*_Ri[T1-1]_ and *cifA_w_*_Ri[T1-2]_. We used Genious Prime to extract ORFs with blast homology to *cif* sequences for downstream analyses (Kearse *et al*. 2012). Among all *cif* sequences, only *cifB_w_*_Mal[T1]_ did not have a clear ORF; however, we did observe sequence homology in the region. To establish sequence integrity, we classified *cif* genes as intact or disrupted based on whether mutations either split the gene into multiple ORFs or truncate the protein, causing loss of a functional domain.

Because recombination can bias phylogenetic inference (Bonneau *et al*. 2018; Cooper *et al*. 2019; Martinez *et al*. 2021), we tested for intragenic recombination using GARD (Kosakovsky *et al*. 2006) and all three statistical methods implemented in PhiPack (Bruen *et al*. 2006). We tested for recombination segments with a minimum length of 100 bp, allowing no more than two breakpoints. GARD calls were visually assessed for reliability. When recombination was detected, we estimated maximum likelihood phylograms for each gene region using RAxML-NG v1.2.2 under the GTR+Γ model with default settings, and performed 1,000 bootstrap replicates to assess node support.

### Approximating *cif-Wovirus* associations

To approximate *cif* associations with *Woviruses*, we searched for contigs containing one or more sr and *cifA* alleles. We then quantified the distance between each *cifA/B* allele and the nearest sr. We evaluated the orientation of sr alleles relative to putatively associated *cifs* and characterized the number and type of genes between them based on NCBI annotation. In the Supplemental Information, we report several lines of evidence that support the plausible association of several *cifA*_[T1]_ alleles and sr alleles of sr3WO (sr3). Because they are the most common *cifs* and srs in our data, we estimated *cifA*_[T1]_ and sr3 phylograms for copies that are plausibly linked. The sequences were aligned with MAFFT v. 7 (Katoh and Standley 2013). We again used GARD (Kosakovsky *et al*. 2006) and all three statistical methods implemented in PhiPack (Bruen *et al*. 2006) to test for recombination in this set of *cifA*_[T1]_ alleles and in the set of associated sr3 alleles. Phylograms for sr3, and separately for GARD-identified partitions, were estimated with RevBayes v. 1.1.1. We used a GTR+Γ+I model with four rate categories and no partitioning. We used a rate multiplier with prior Γ(1,1) [*i.e.*, Exp(1)], as well as stationary frequencies and exchangeability rates drawn from flat, symmetrical Dirichlet distributions [*i.e.*, Dirichlet(1,1,1.…)]. The model used a uniform prior over all possible topologies. Branch lengths were drawn from a flat, symmetrical Dirichlet distribution and thus summed to 1. Four independent runs were performed, and all converged to the same topology. For additional details on the priors and their justifications, consult Turelli *et al*. (2018).

### Characterizing Cif protein structures

We ran HHPred on a Linux kernel to identify putative functional domains using the Pfam-A_v35 and SCOPe70_2.07 databases (Zimmermann *et al*. 2018). We considered only annotations with > 80% probability. If alternative annotations produced probability > 0.8, we selected the annotation with the highest probability. We used AlphaFold2 (Jumper *et al*. 2021) to predict Cif structures. Entries in the “reduced database” provided with AlphaFold prior to 5/10/22 were used to generate multiple sequence alignments (MSA) within AlphaFold. We generated five structures for each protein and performed amber relaxation to prevent unrealistic folding patterns. We sorted the five relaxed models by mean pLDDT and used the top result in other analyses. We visualized protein structures using PyMol 2.5.2 (Schrödinger, LLC 2015) and aligned Cif_[T1]_ proteins relative to Cif*_w_*_Mel[T1]_ for imaging using Cealign.

To investigate Cif protein sequence and structural similarity, we calculated pairwise amino-acid identity ID and template-modeling scores (TM) (Zhang and Skolnick 2004). We used Muscle 5.0 to generate an ensemble of 100 MSAs from CifA and CifB protein sequences. From the ensemble, we extracted the alignment with the highest confidence score for each column, and calculated pairwise amino-acid identity from this alignment. TM-scores indicate the degree of similarity between two superimposed structures, with TM = 0 indicating no similarity, TM = 1 corresponding to complete congruence, and TM > 0.5 indicating a homologous structure (Xu and Zhang 2010). We generated TM-scores for Cif proteins pairwise in PyMol using the psico module (Holder 2023). We performed each analysis twice, switching the reference and target proteins. The directionality of the structure impacts the TM-score because the score is normalized to the length of the target protein. We used a Mann-Whitney U test in R to compare the TM-scores from CifA and CifB, excluding those that were either truncated or positioned at the edge of a contig. We assessed the relationship between TM-score and percent identity using a Spearman correlation.

### Characterizing selective pressures on *cifs*

Our tests for selection on *cifs* rely on the estimated normalized ratio of non-synonymous (*d_N_*) to synonymous (*d_S_*) substitutions. We focus our analyses on *cifA/B*_[T1]_ since Type 1 copies are particularly common among *w*Mel-like and *w*Ri-like *Wolbachia.* The ratio *ω* = *d_N_*/*d_S_* can be unstable on very short timescales, as the estimates fluctuate due to stochastic sampling effects and *ω* approaches its long-term expectation only asymptotically (Yang and Nielsen 2000; Kryazhimskiy and Plotkin 2008). On very long timescales, high *d_S_* leads to saturation and reduces power (Anisimova *et al*. 2001). For our analyses, we selected putatively intact *cifA/B*_[T1]_ and disrupted *cifA*_[T1]_ copies that met 0.1 ≤ *d_S_* ≤ 2.0, which is established as a conservative and suitable range for analyses of *ω* (Yang and Bielawski 2000; Yang 2006).

The majority of putatively intact *w*Mel-like and *w*Ri-like *cifA/B*_[T1]_ copies are closely related to one another and distantly related to a clade of four copies found in *w*Dal, *w*Zta, *w*Spa, and *w*Ara. We excluded *w*Ara from our analyses, because *cifA/B*_[T1]_ in *w*Spa and *w*Ara are nearly identical, with only one difference in *cifB*_[T1]_ and four in *cifA*_[T1]_. We used the *cifA/B*_[T1]_ copies from *w*Dal, *w*Zta, *w*Spa, and *w*Mel for all analyses of intact *cif*_[T1]_ described below (pairwise *d_S_* range: *cifA*_[T1]_ = 0.23–0.35, *cifB*_[T1]_ = 0.38–0.48). Relative to the *w*Mel copies, all *cif*_[T1]_ copies observed in *w*Ri-like genomes and the other *w*Mel-like genomes were too closely related for inclusion (see Fig. 5C). Our analyses focus on the three putatively functional *cifB*_[T1]_ domains, Nuc1, Nuc2 and Dub, involved in CI (Beckmann *et al*. 2017; LePage *et al*. 2017; Chen *et al*. 2019; Beckmann *et al*. 2021; Kaur *et al*. 2022), which we annotated on the codon alignment. We also annotated *cifB*_[T1]_ non-domain regions. Given an insufficient number of disrupted *cifA*_[T1]_ copies in our *w*Mel-like and *w*Ri-like genomes, we searched for additional copies in the data of Martinez *et al*. (2021). Our analyses included *w*Bic from *D. bicornuta*, *w*Stv from *D. sturtevanti*, and *w*Nfla from *N. flava,* along with the disrupted *w*Mel-like *w*Ach copy (pairwise *d_S_* range: 0.34 – 0.86). Results related to disrupted *cif* _[T1]_ copies are presented in the Supplemental Information. We used these data and multiple methods to compare selection on *cifA* and *cifB*, and separately, to compare selection on the three *cifB*_[T1]_ protein domains (Nuc1, Nuc2 and Dub) and associated non-domain *cifB*_[T1]_ regions.

### codeml analyses

We applied several likelihood-based models implemented in PAML using codeml (Yang 1997, 2007). We aligned our sets of sequences using TranslatorX’s PRANK mode (Abascal *et al*. 2010). We also used a direct PRANK alignment with flags-F to skip insertions and-codon that uses an empirical codon model (Löytynoja 2014). The latter was used only to assess the sensitivity of a subset of results to alignment, and such cases are noted explicitly as they appear. In all cases, we visually inspected the alignments for frameshifts, finding none. For disrupted *cifA*_[T1]_, we first aligned with MAFFT v7 using default parameters, and then manually deleted any frameshift-causing insertions, and manually filled any frameshift-causing deletions with “N”. We then realigned sequences using the TranslatorX PRANK mode unless noted otherwise. All codon-based analyses used the Goldman-Yang codon model (GY94) (Goldman and Yang 1994) with F3×4 codon frequencies (CodonFreq = 2) (Nielsen and Yang 1998; Yang *et al*. 2000). For each analysis, we initiated model fits from multiple starting values for the transition/transversion rate parameter (*κ*) (0.5, 1.0, 2.0) and *ω* (0.25, 0.5, 0.75, 1.0) to account for potential sensitivity of likelihood maximization to initial values. For each sequence pair, we used the estimates of (*κ*, *ω*) that produced the highest log-likelihood across the 12 combinations of initial values. We first assessed *ω* using runmode = –2 for pairwise comparisons. We focus on the median *ω,* rather than the mean, because of our small sample size (*N* = 6).

Using paired contrasts, we tested whether *ω_cifA_* < *ω_cifB_* using a one-sided Wilcoxon signed-rank test. We report the Hodges–Lehmann (HL) estimator (Hodges and Lehmann 1963) of the median difference with an exact 95% confidence interval obtained by inverting the signed-rank test (Hollander *et al*. 2015).

Confidence intervals are obtained by identifying the set of parameter values that would not be rejected by the Wilcoxon signed-rank test, *i.e.,* by inverting the test (Lehmann 1975). To test whether *ω_cifA_*_-intact_ < *ω_cifA_*_-disrupted_ using unpaired *cifA*_[T1]_ copies, we applied the Mann-Whitney test with a one-sided alternative and the corresponding two-sample HL estimator that provides an exact 95% CI via rank-sum inversion. We also estimated nonparametric Bayesian effect sizes using the Bayesian bootstrap. The Bayesian bootstrap is analogous to the classical bootstrap, but instead of resampling observations with replacement, it assigns random weights to each observation drawn from a Dirichlet(1,…,1) distribution. This can be implemented by normalizing independent Exp(1) samples, which yields the same distribution of weights. For each contrast, we applied these random weights to compute the linearly interpolated weighted median of the pairwise differences (the 0.5 quantile of the weighted empirical CDF with linear interpolation). We repeated this procedure *B* = 20,000 times to obtain a posterior sample of median effects. From this posterior, we report the posterior mean of the median effect, the 95% equal-tailed credible interval, and the posterior probability of the hypothesized directional effect, *P*(Δ < 0). These Bayesian summaries are reported alongside the Wilcoxon and Mann–Whitney results.

We also analyzed codon models on phylogenetic alignments to estimate lineage-specific and site-specific variation in *ω*. Because recombination can bias phylogenetic inference and has been observed for *cifs* (Bonneau *et al*. 2018; Cooper *et al*. 2019; Martinez *et al*. 2021), we first tested for intragenic recombination using GARD (Kosakovsky *et al*. 2006) and all three statistical methods implemented in PhiPack (Bruen *et al*. 2006), as described in the *cif* extraction section above. When recombination was detected, we estimated maximum likelihood phylograms for each gene region using RAxML-NG v1.2.2 under the GTR+Γ model with default settings, and performed 1,000 bootstrap replicates to assess node support. We partitioned our data according to their respective unrooted topologies in all subsequent selection analyses. This produced highly supported two-part partitions for intact *cifA*_[T1]_ and disrupted *cifA*_[T1]_.

We evaluated six models in codeml that differ in their treatment of variation in *ω* across lineages and across sites. The one-ratio model (M0) assumes a single *ω* across all sites and lineages, and the branch model allows *ω* to vary among lineages. We assigned each ingroup branch to its own category, while constraining the two branches associated with the outgroup to share a single *ω*. This treats the tree as effectively unrooted under the branch model. To test for heterogeneity among sites, we fit models M1a, M2a, M7, and M8. The “nearly neutral” model (M1a) assigns sites to classes with *ω* < 1 or *ω* = 1, while the positive-selection model (M2a) adds a class with *ω* > 1. Model M7 assumes a beta distribution of *ω* across sites, restricted to values between 0 and 1, while the beta plus positive-selection model (M8) adds an extra site class in which *ω* estimates exceed 1. Each recombination partition was analyzed under its GARD and RAxML-supported topology, and likelihood-ratio tests (LRTs) were used for partition-specific model comparisons.

For models that include a positive-selection class (M2a, M8), we obtained BEB posterior probabilities and reported sites with posterior *P* ≥ 0.95, prioritizing sites concordant across M2a and M8 (Yang *et al*. 2005). We visually evaluated alignments for these sites, looking for evidence of multi-nucleotide mutations (MNMs) that may produce false signatures of positive selection. MNMs that occur simultaneously – *e.g.,* due to error-prone DNA polymerases – violate codon substitution models that assume mutations occur independently (Venkat *et al*. 2018). For downstream site-specific analyses, we masked sites supported by BEB with evidence of MNMs. For comparisons of domains to non-domain regions within *cifB*_[T1]_ to test whether *ω*_Domain_ < *ω*_Non-domain_, we summarized means per-site posterior *ω* by region from model M8 or M7 and their 95% bootstrap confidence intervals, resampling sites within each region 10,000 times. We assessed significance with a site-level label permutation test, treating sites as exchangeable. We computed the observed Δ (*ω*_Domain_ – *ω*_Non-domain_) and generated an empirical null by permuting region labels 10,000 times. *P* values were obtained by comparing the observed statistic to the permutation-based null distribution. We computed 95% confidence intervals by bootstrapping the difference of means directly – resampling sites with replacement within each region 10,000 times – and taking percentile bounds. We report effect sizes, permutation *P* values, and 95% confidence intervals from codon-level bootstrap resampling within partitions.

### SWAKK Analyses

We also used the Sliding Window Analysis of Ka and Ks (SWAKK) webserver to evaluate constraint on putatively intact CifA_[T1]_ and on the three CifB_[T1]_ domains (Liang *et al*. 2006). SWAKK generates an alignment of two nucleotide sequences and maps it onto a provided tertiary structure. The average amino acid width spans ∼3.8-4.3 Å (Jumper *et al*. 2021), and SWAKK calculates *ω* (*i.e.,* Ka/Ks) using a 10 Å spherical sliding window across the reference structure. This enables tests for selection on regions within three-dimensional (3D) structures that may be functionally important. We used AlphaFold structures for tertiary mapping with CifA/B*_w_*_Mel[T1]_ as our reference, calculating *ω* for pairwise comparisons between the *w*Mel proteins and each of the three other putatively intact CifA/B_[T1]_ proteins described above. We filtered SWAKK output to include only sites where 0.1 ≤ Ks ≤ 2.0 and pLDDT ≥ 70, which is considered a confident prediction for a given amino acid residue in an AlphaFold protein structure model (Jumper *et al*. 2021; Guo *et al*. 2022). Undefined entries and sentinel rows with Ks = 99 were excluded.

For each pair, we computed the difference of the median log(*ω* + 10^-6^) between CifA_[T1]_ and CifB_[T1]_ (Δ), where Δ = median{log(*ω* + 10^-6^)}_CifA_ − median{log(*ω* + 10^-6^)}_CifB_. To summarize the effect size, we averaged the three Δ values to obtain the mean of the per-pair median differences on the log scale (Δ*_p_*).

We also computed the geometric-mean ratio [exp(Δ*_p_*)], where exp(Δ*_p_*) < 1 indicates lower CifA_[T1]_ *ω.* To obtain 95% BCa (bias-corrected and accelerated) intervals for Δ*_p_*, we used a non-circular, non-overlapping moving-block bootstrap (Künsch 1989; Liu and Singh 1992) with B = 5,000 resamples. This approach preserves the short-range autocorrelation introduced by the 10 Å window. Within each pair and protein, we estimated the integrated autocorrelation time (IAT) of the ordered log(*ω* + 10^-6^) series (Sokal 1997). Provisional block lengths were set to 2×IAT for each series, rounded to an integer, and truncated to the range 6–30 residues. The block length established for each pair (*L*) was the maximum of the CifA_[T1]_ and CifB_[T1]_ provisional lengths. For each bootstrap replicate, we resampled contiguous blocks with replacement until their combined length equaled or exceeded the original series length, truncating any excess to match the original size. The three Δ values were then recomputed and averaged to obtain a bootstrap Δ*_p_*. BCa intervals used jackknife acceleration computed by leave-one-pair-out Δ*_p_*. To obtain a one-sided *P* value for (Δ*_p_* < 0), we used a block label permutation within each pair that preserves local dependence (Good 2005; Winkler *et al*. 2014, Winkler *et al*. 2015). We partitioned the ordered log(*ω* + 10^-6^) series for CifA_[T1]_ and CifB_[T1]_ within each comparison into non-overlapping blocks of length *L* (same as in the bootstrap), pooled all blocks, and randomly reassigned blocks back to CifA_[T1]_ or CifB_[T1]_ independently for each label without replacement until the original series lengths were matched. For each permuted dataset, we recomputed Δ for each comparison and then averaged across comparisons to obtain Δ*_p_*. The one-sided *P*-value for (Δ*_p_* < 0) is (r+1)/(B+1), where r is the number of permuted Δ*_p_* values ≤ the observed Δ*_p_*, and B = 10,000 permutations.

To evaluate constraint on CifB_[T1]_ domains, we compared Nuc1, Nuc2, and Dub CifB_[T1]_ domains individually to pooled non-domain regions. For each domain, we computed Δ = median{log(*ω* + 10^-6^)}_domain_ − median{log(*ω* + 10^-6^)}_non-domain_. We averaged the three Δ values to compute Δ*_p_*, with exp(Δ_p_) interpreted as the domain/non-domain *ω* ratio. All domain analyses used the same moving-block bootstrap/BCa framework and the same permutation logic as above, except *L* was chosen separately for domain and non-domain regions within each pair. Permutations reassembled both series independently from the pooled blocks, ensuring that a block can appear in both labels within a permutation. We excluded the *w*Mel-*w*Dal pair from the Dub analysis due to a paucity of domain windows.

### Figure Generation

We produced and/or edited figures in R, Figtree, Inkscape 1.1, Adobe Illustrator, and Keynote.

