## Supplementary Material (File S1) for "Calibrating and documenting host-switching and evolution of incompatibility loci for two closely related *Wolbachia* clades"

Our supplementary materials are ordered to correspond to the sections of the main text, with matching section and subsection headings when possible. For completeness, we reiterate some of the most relevant results from the main text to make the summaries below relatively self-contained. However, we also present new results that we believe are less significant than the key findings presented in the main text. An example is our discussion of *w*Zts as the closest known variant to *w*Mel.

### Supplementary Introduction

#### Progress based on increasingly comprehensive molecular data and new calibrations

In the first analysis of *Wolbachia* divergence times, Werren *et al.* (1995) used a "universal molecular clock" for bacteria (Ochman and Wilson 1987) applied to *ftsZ* gene sequences of facultative *Wolbachia* extracted from flies and wasps, host species that diverged approximately 350 million years ago (MYA; Wang *et al.* 2016). Based on no differences over 265 synonymous substitution sites within a 937 bp region of *ftsZ* between *Wolbachia* in the parasitic hymenopteran *Asobara tabida* and its dipteran host *Drosophila simulans* (from Riverside California; Hoffmann *et al.* 1986), Werren *et al*. (1995) estimated a 95% (99%) confidence interval for *Wolbachia* divergence of only 0–1.6 (0–2.5) million years (MY). Raychoudhury *et al.* (2009) cross-calibrated facultative *Wolbachia* divergence with host nuclear and mtDNA divergence and developed a refined chronology of *Wolbachia* movements, exploiting a plausible example of cladogenic acquisition (*i.e.*, codivergence) of facultative *Wolbachia* in the wasps *Nasonia longicornis* and *N. giraulti*. The key evidence supporting codivergence was concordant divergence-time estimates for the hosts and some of their *Wolbachia*, based on independently derived molecular clocks for synonymous-site divergence of eukaryotic nuclear genes and an updated rate estimate for coding-region divergence across bacteria (Ochman *et al.* 1999). For the host-*Wolbachia* pairs showing plausible cladogenic *Wolbachia* transmission, *Wolbachia* divergence was estimated at about one-third the rate of the host nuclear genomes for synonymous sites. In contrast, mtDNA diverged at synonymous sites about $1.2\times10^{2}$ times as fast as co-inherited *Wolbachia.* The Raychoudhury *et al*. (2009) data imply an average *Wolbachia* yearly substitution rate at third-position sites of approximately $2.2\times10^{-9}$ (see our Materials and Methods).

Richardson *et al*. (2012) produced an alternative approach to calibrating rapid *Wolbachia* divergence, comparing full genomes of facultative *w*Mel *Wolbachia* (Hoffmann 1988) and mitochondria among *Drosophila melanogaster* lineages. As expected under joint maternal inheritance, *D. melanogaster* isofemale lines produced generally concordant mtDNA and *w*Mel *Wolbachia* phylogenies (but see Scholz *et al*. 2020). Having estimated *relative* sequence divergence for *Wolbachia* and mtDNA among isofemale lines, Richardson *et al*. (2012) estimated an *absolute* rate for *Wolbachia* evolution, based on observed mtDNA mutations. Using the per-generation mtDNA mutation rate as a prior in a Bayesian analysis of isofemale-line divergence, they estimated a "short-term" *Wolbachia* third-site substitution rate of $6.87\times10^{-9}$ per site per year (see our Materials and Methods). This enabled Turelli *et al*. (2018) to estimate divergence times for facultative *Wolbachia* closely related to *w*Ri in *D. simulans* (Hoffmann *et al*. 1986).

They estimated that "*w*Ri-like" strains which diverged less than 30 thousand years ago (KYA) occupy *Drosophila* hosts that diverged 10–50 MYA. Ahmed *et al.* (2016) used a similar calibration to estimate horizontal transmission times for *Wolbachia* among Lepidoptera species. As reported in the main text, our new (long-term) nematode calibration rates are significantly slower than the mutation-based calibration proposed by Richardson *et al.* (2012). The new calibrations suggest that Turelli *et al.* (2018) may have underestimated the divergence of closely related facultative *w*Ri-like variants associated with divergent *Drosophila* hosts by about a factor of seven. Our alternative calibrations do not alter the qualitative conclusion of Turelli *et al.* (2018) that several orders of magnitude separate the divergence times of many *Drosophila* hosts from their current *w*Ri-like *Wolbachia*.

### Supplementary Results

#### Host chronograms and *Wolbachia* phylogenies

**Hosts for *w*Mel-like *Wolbachia*.** We describe the insects in our study that carry *w*Mel-like *Wolbachia* (Fig. 1; Tables 1 & S1). These hosts are diverse taxonomically and in their life histories, ranging from cosmopolitan human-associated fly species (*D. melanogaster*, *D. simulans* and *S. pallida*) to endemics restricted to small oceanic islands (*D. santomea* and *D. arawakana*). The drosophilid dipterans include a species that breeds and feeds on flowers (*D. incompta*), a mushroom specialist (*D. recens*), and generalists (*e.g., D. melanogaster* and *D. simulans*). As expected, hosts with closely related *Wolbachia* co-occur (or did in the recent past). For instance, *w*Au was previously observed in *D. simulans* in Florida and Ecuador (Turelli and Hoffmann 1995) before *w*Au was displaced by *w*Ri, and *w*Au-carrying *D. simulans* probably co-occurred with *D. tropicalis* restricted to Central and South America and the Caribbean islands, where *D. tropicalis* harbors *w*Tro, sister to *w*Au. However, close associations of hosts need not lead to *Wolbachia* horizontal transfer. For instance, although the hymenopteran wasp *Diachasma alloeum* parasitizes the tephritid *Rhagoletis pomonella*, none of *R. pomonella*'s several *Wolbachia* seem to be *w*Mel-like (Schuler *et al.* 2011).

According to the fossil-calibrated chronograms in Wang *et al*. (2016, their Fig. 3), Diptera and Hymenoptera diverged ~350 MYA, with 95% highest posterior density credibility interval (HPD CI) 378–329 MYA (Devonian–Carboniferous). This is consistent with the point estimates produced by Misof *et al.* (2014) and Johnson *et al.* (2018, Fig. 1) and comparable to the crown age of all extant tetrapods, ~373 MY (Simões and Pierce 2021). The placement of the family Diopsidea, the stalk-eyed fly clade that includes *Sphyracephala rbevicornis*, within superfamily Diopsoidea in the paraphyletic acalyptrate group of Schizophora, remains uncertain (Bayless *et al.* 2021). Hence, the maximum divergence time between *S. brevicornis* and any drosophilid is the crown age of the Schizophora, which includes both the Drosophilidae and Diopsoidea. The minimum divergence time between the Drosophilidae and Diopsoidea is the crown age of the Drosophilidae (which certainly excludes the Diopsoidea). Wiegmann *et al.* (2011, Fig. 3) estimate the crown age of the Schizophora at ~70 MYA. Suvorov *et al.* (2022, Fig. 1) estimate the crown age of the Drosophilidae at ~47 MYA (with 95% HPD CI of 43.9–49.9 MYA). Our approximate point estimate in Fig. 1B for the divergence of Drosophilidae and Diopsoidea, 59 MY, is the midpoint of these upper- and lower-bound point estimates. We number the nodes in Fig. 1B based on their divergence and present the node ages and approximate confidence intervals in Table 2.

**Assessing support for bifurcating evolution of major portions of *Wolbachia* genomes across two orders of magnitude of genomic divergence**

**Analyzing intragenic recombination.** We first assessed evidence for intralocus recombination using a genetic algorithm for recombination detection (GARD) (Kosakovsky Pond *et al.* 2006), plus all three statistical methods implemented in PhiPack (Bruen *et al.* 2006). These analyses focus on single-copy genes that are least 300 bp in length, with minimum recombination segments of 100 bp. The 300 bp criterion excludes a small number of the single-copy genes within each set described above. For the set of *w*Ri-like *Wolbachia*, we found no evidence for intragenic recombination within the 517 genes that met our criteria, and we found almost no evidence of intragenic recombination within any of the 411 genes that met our criteria for our set of *w*Mel-like *Wolbachia*: GARD: 0/411 genes; PHI test: 0/411, Max $\chi 2$ test: 1/411, Neighbor Similarity Score: 0/411, FDR-adjusted $P < 0.01$). It is notable that *wsp* does not meet our alignment criteria for either of these recombination analyses (cf. Baldo *et al*. 2005); whereas five of the other six MLST loci, *coxA, ftbA, ftsZ, gatB* and *hcpA* meet the criteria for our *w*Ri-like analysis; and of those, all but *coxA* appear in our *w*Mel-like analysis. In summary, these comparisons of young *Wolbachia* clades indicated little to no intragenic recombination within our gene sets. It is worth noting that given the extreme similarity of the *w*Ri-like genomes, recombination between variants within this clade would be essentially impossible to detect, and our ability to detect recombination increases with the divergence of our sets of variants.

We next tested for intragenic recombination in four samples of supergroup D *Wolbachia* with significantly higher levels of sequence divergence: (*w*Wb in *W. bancrofti*, (*w*Bm in *B. malayi, w*Bp in *B. pahangi*)) and outgroup *w*Ls from *L. sigmodontis*. Over 356 genes (349,602 bp), *w*Wb differs from *w*Bm by 2.23% and from *w*Ls by 9.21%; whereas over 442 genes (413,115 bp), *w*Mel differs from *w*Tri by 1.48% and *w*Ha differs from *w*Bic by 1.67%. Again, we found almost no evidence of recombination within any of the 356 genes that met our criteria for this comparison: GARD: 0/356 genes; PHI test: 0/356, Max $\chi 2$ test: 1/356, Neighbor Similarity Score: 0/356, FDR-adjusted $P < 0.01$). In summary, these results indicated little to no intragenic recombination for these relatively anciently diverged obligate *Wolbachia*.

To supplement our tests of the younger *w*Mel-like and *w*Ri-like clades and the older supergroup C and D clades presented in the main text, we tested for intragenic recombination among a larger set of supergroup A *Wolbachia* that spans the *w*Mel-like and *w*Ri-like clades. The seven *Drosophila*-derived variants have consensus phylogeny: (((*w*Mel, *w*Au), (*w*Bic, *w*Bar)), ((*w*Ri, *w*Tri), *w*Ha)) (Fig. 3). In contrast to our analyses of the *w*Mel-like and *w*Ri-like *Wolbachia* clades, we found putative evidence of recombination within many of the 401 genes that met our criteria: GARD: 187/401 genes; PHI test: 40/401, Max $\chi 2$ test: 110/401, Neighbor Similarity Score: 63/401, FDR-adjusted $P < 0.01$. However, inspection of the trees underlying the GARD results suggested that many of these recombination calls may be artifacts. GARD attempts to estimate separate maximum likelihood trees for putatively recombined regions, but does so without bootstrap replicates to assess support for the estimated topologies. When sufficient variation exists to estimate a tree, GARD always uses a fully resolved tree, regardless of its statistical support.

We illustrate how artifactual recombination calls emerge using the gene with the second highest number of variable sites across these seven strains. As explained in the text, highly variable loci are strong *a priori* candidates for horizontal acquisition of sequence fragments. The locus is homologous to RefSeq_WP_010962405.1 and annotated as a peptide deformylase (540 bp). In principle, GARD identifies loci that have undergone recombination by finding alternative topologies for different gene segments. However, without assessing the levels of support for those topologies, the algorithm seems highly error prone when closely related sequences are considered. For this gene, GARD identified a 163 bp recombined region based on a fully resolved tree for this region, even though five of the seven samples are identical in this region and thus can only be resolved randomly. Using a random topology for the five identical samples, GARD identified this region as recombinant. Based on this example and several others, it seems that GARD produces false positives for recombination when there is enough variation to estimate a topology but not enough variation to produce strong support. We believe the ad hoc alternative procedures we described in the text more accurately identify histories of recombination in such cases. These alternative procedures found strong support for bifurcating evolution over major portions of these seven supergroup A *Wolbachia* genomes.

**Analyzing polymorphism and allelic differences at individual loci.** As with our discussion of the *w*Ri-like variants in the main text where there were no two-fold outlier loci, it is useful to have some benchmarks for allelic differences among our *w*Mel-like variants. If we compare the two variants, *w*Rec and *w*Zts, that show the largest sequence divergence over the 438 single-copy loci in our analyses, only 12 loci show 6 or more nucleotide differences and 325 loci show at most 2 differences. Hence, horizontal acquisition of distantly related alleles at single-copy loci must be rare over the timescale of *w*Mel-like divergence.

To identify outlier alleles in the *w*Mel-like gene set, we ranked loci by the fraction of variable nucleotides, under the assumption that the most variable loci are the most promising candidates for horizontal transmission. Among the 20 *w*Mel-like loci with the largest fraction of variable nucleotides, only 4 showed a two-fold outlier allele (Table S6), and only the second most-variable gene initially appeared as a plausible candidate for horizontal acquisition of distantly related alleles (Table S6). This gene is homologous to RefSeq_WP_010962738.1, peptidase M48, with 43 variable sites across the 1,282 bp (3.35%). The *w*Bor allele differs from the other 19 alleles at 34–36 sites (2.7–2.8%), while the other alleles differ from one another at fewer than 5 sites (≤0.39%). BLAST found that the *w*Bor allele shares

99.9% similarity with an allele in *w*Inn from *D. innubila*. In the main text we present evidence that our data are consistent with IS activity within the genome of the *w*Inn-*w*Bor MRCA generating the 55 bp fragment that distinguishes them from the 19 other alleles in our analysis. We extracted 365 single-copy genes (334,889 bp) from the 20 *w*Mel-like, *w*Inn, and *w*Ri and *w*Tri as outgroups. We estimated a phylogram (see Materials and Methods). The phylogram shows that *w*Inn is yet another *w*Mel-like *Wolbachia*, most closely related to *w*Bor (Fig. S8).

The other three most variable genes also showed no evidence for horizontal acquisition of distantly related alleles. The locus with the largest fraction of variable sites is homologous to NCBI reference RefSeq_WP_006279576.1, NADH-quinone oxidoreductase subunit A, at which 12 of 354 nucleotide sites (3.39%) differ among the 20 alleles. The allele from *w*Inc differed from the other 19 alleles at 8–10 sites (2.3–2.8% different), whereas the other 19 alleles differed from one another at 3 or fewer sites (≤ 0.8% different). When compared to the most distant allele in *w*Seg, 8 of 10 variable sites occurred within 12 positions of a deletion at 346–348. BLAST of this deletion and the remaining downstream 3’ *w*Inc sequence revealed no hits. The fourth most variable gene is homologous to RefSeq_WP_010963163.1, a ribonuclease, with 21 of 816 sites variable (2.57%). The allele in *w*Ara differed from the 19 others at 12–16 sites (1.5–2.0%), while the other 19 alleles differed from one another at 6 or fewer sites (maximum 0.73%). When compared to the most distant allele in *w*Rec, 13 of 16 variable sites occurred within 9 bp of either side of a deletion at positions 736–804. BLAST revealed that the first 727 bp 5’ of this deletion share 97% similarity to several alleles in other *w*Mel-like *Wolbachia*. There are only 12 bp 3’ of the deletion in the *w*Ara allele, with the deletion plausibly affecting the alignment. Finally, the fifth most variable gene is homologous to RefSeq_WP_010962626.1, a transcriptional regulator, with 19 of 742 sites variable (2.56%). The allele in *w*Inc differed from the 19 others at 13–15 sites (1.8–2.0%), while the other 19 alleles differed from one another at 4 or fewer sites (maximum 0.54%). When compared to the most distant allele in *w*Seg, 12 of the 15 variable sites occur within 19 bp of a deletion at positions 701–706. The *w*Inc allele also BLASTS uniquely to itself.

In summary, we observed no evidence for horizontal acquisition of distantly related alleles at single-copy loci meeting our criteria in the *w*Mel-like gene set. In the main text we illustrate evidence for only a single horizontal transfer from a distantly related *Wolbachia* in the seven supergroup A set involving the most variable locus meeting our two-fold outlier criteria: the gene homologous to RefSeq_WP_010962975.1, a membrane protein. The *w*Ha allele showed 92–94 bp differences from the homologous alleles in the other 6 *Wolbachia* (18.5–19.9%), whereas the other 6 alleles had a maximum of 3 pairwise differences (≤ 0.6%). BLAST revealed that the *w*Ha copy is identical to copies in *Wolbachia* from divergent host species, including sawflies, hoverflies, moths, carpet beetles, and Tachinidae flies (*e.g.,* Accessions: OZ034713.1, OX366332.1, AP028948.1, OX366359.1, OZ034764.1). Hence the majority of single-copy loci in our analyses did not acquire exogenous alleles over the timescales of differentiation for our *w*Ri-like and *w*Mel-like variants.

**Analyzing the clustering of loci supporting non-consensus phylogenies.** Using our seven reference supergroup A *Wolbachia*, we attempted to assess the size and scope of horizontally acquired genomic fragments by examining the locations of: 1) loci that support "non-consensus" topologies and 2) loci that show allele differences suggestive of horizontal acquisition. Because of rearrangements among the seven reference *Wolbachia* genomes, we describe the genomic locations of loci that seem candidates for horizontal allele acquisition using the location notation corresponding to four alternative reference genomes: *w*Mel, *w*Au, *w*Ri and *w*Ha, with consensus phylogeny ((*w*Mel, *w*Au), (*w*Ri, *w*Ha)) (Fig. 3).

Over the seven supergroup A *Wolbachia* genomes, we compare the levels of divergence for two classes of phylogenetically informative loci: those that support the consensus topology, versus those that support alternative topologies. We used two measures of divergence associated with these loci: the fraction of polymorphic sites and the estimated number of substitutions per site. The latter was estimated from the total length of phylograms produced with GTR+I+Γ substitution model with no partitioning among codon positions. The levels of differentiation for the two classes of loci were compared with a *t*-test.

Using this approach, we examined the spatial arrangement of the 38 loci that supported non-consensus topologies (see main text). If large fragments had been acquired horizontally, we would expect spatial clustering of the loci supporting specific alternative topologies. Table S4 provides no evidence of such clustering. Across all 38 loci, there was only one example of two "adjacent" phylogenetically informative loci that supported the same alternative topology, namely those homologous to RefSeq_WP_010963073.1, phosphoglycerate kinase**,** and RefSeq_WP_007548470.1, coproporphyrinogen III oxidase, separated by about 50K bp. Both loci supported topology A7, and they were the only loci that supported A7. They did not show either high levels of allelic differentiation or large total branch lengths that would be expected if they were horizontally acquired. As discussed in the main text, the conflicting phylogenetic signals may have arisen from horizontal transmission of multiple *Wolbachia* genomes between host species. An alternative explanation involves ambiguous phylogenetic resolution of relatively short branches.

#### ***Wolbachia*** **substitution-rate calibrations**

We summarize the following previously applied *Wolbachia* calibrations that we compared to our new filarial nematode-based calibrations in the main text (Tables 3 & 4).

**Applying "universal" rates of bacterial evolution to** ***Wolbachia*****.** Ochman and Wilson (1987) estimated a "universal" bacterial synonymous-site substitution rate of 7–8×$10^{-9}$ substitutions per synonymous site per year, using differences observed over 21 kb of coding sequences between *E. coli* and *Salmonella typhimurium*. Applying this calibration to partial *Wolbachia ftsZ* coding sequences from the *Wolbachia* in *D. simulans* and its parasitic wasp *Asobara tabida,* Werren *et al*. (1995) inferred that these *Wolbachia* were separated by only 0–2.5 MY, making horizontal acquisition seem plausible. However, substitution rates differ by several orders of magnitude among loci and among bacterial lineages (*e.g.*,

Duchene *et al.* 2016; Gibson and Eyre-Walker 2019), and *ftsZ* sequences seem to diverge particularly slowly. If instead, we look at 560 bp of *wsp* coding sequence, *w*Ri differed from each of the three *Wolbachia* variants found in *Asobara tabida* by at least as much as *w*Ri differed from *w*Mel (Table S7). Based on our new genomic calibrations provided below, the MRCA of *w*Ri and *w*Mel occurred at least 1.4 MYA. This is comparable to the divergence times of *D. simulans* from *D. sechellia* and *D. mauritiana* (Suvorov *et al*. 2022), precluding recent horizontal acquisition of *Wolbachia* between *D. simulans* and its parasitoid A. *tabida*. As illustrated by Conner *et al*. (2017), reliable evaluation of modes and timing of *Wolbachia* acquisition require genome-scale data and calibrations.

***Wolbachia* calibration from *Drosophila*.** Richardson *et al.* (2012) combined the relative rates of divergence of *Wolbachia* and mtDNA genomes among *D. melanogaster* lines with an estimate of the mtDNA mutation rate per generation to infer a "short term" rate of evolution of third-position sites in *Wolbachia* coding regions of $6.87\times10^{-9}$ per third-position site. Based on our new rate estimates from filarial nematodes (see Table 3), which were consistent with estimates from *Nasonia* and *Nomada*, the Richardson *et al.* (2012) rate may be too fast by about a factor of seven.

***Wolbachia* calibration from *Nasonia* wasps.** While rough agreement of two plausible molecular clocks provided evidence for *Wolbachia*-*Nasonia* codivergence, Raychoudhury *et al.* (2009) note that their codivergence hypothesis was only tentatively supported because substitution rates vary across taxa. Their *Wolbachia* divergence-time estimate of 0.41 MYA for the MRCA of *w*NgirB and *w*NlonB1uses an *E. coli* synonymous substitution rate of $4.5\times10^{-9}$, from Table 1 of Ochman *et al*. (1999). An alternative and higher *Buchnera* synonymous substitution rate of $8.2\times10^{-9}$ also appears in Table 1 of Ochman *et al*. (1999) and is comparable to the mutation-based estimate of Richardson *et al*. (2012). Ochman *et al.* (1999) postulated that because of smaller effective population sizes, higher substitution rates might be expected for endosymbionts, like *Buchnera* and *Wolbachia*, versus free-living bacteria, like *E. coli*. In contrast to the single *E. coli*-based divergence-time estimate used by Raychoudhury *et al.* (2009), the *Buchnera* calibration produces 0.23 MYA for the MRCA of these *Wolbachia*. This alternative suggests horizontal versus cladogenic *Wolbachia* acquisition for one of the *Nasonia* hosts. However, our new nematode calibrations are consistent with the Raychoudhury *et al.* (2009) codivergence hypothesis.

We averaged the eukaryotic-based and prokaryotic-based divergence-time estimates to approximate age of the MRCA of both *N. giraulti* and *N. longicornis* and *w*NgirB and *w*NlonB1 as about 0.46 MYA. We reanalyzed the raw data available from Raychoudhury *et al.* (2009) for these host nuclear genomes, their mtDNA, and these *Wolbachia* variants to produce the divergence estimates in the first line of Table 3. Doubling the age estimate of the MRCA produced the *Wolbachia* substitution rate estimates given in Table 4. Our synonymous rate differs slightly from that reported in Table 3 of Raychoudhury *et al.* (2009) because of our different divergence-time approximation. Importantly, despite the uncertainties, the *Nasonia*-based rate estimates in our Table 4 are consistent with estimates derived from filarial nematodes for which much stronger evidence supports *Wolbachia*-host codivergence. Empirical variation in rates estimated from different *Wolbachia* lineages provides a biological basis for plausibly estimating the uncertainty of substitution rates, beyond the statistical uncertainty associated with individual calibrations.

***Wolbachia* calibration from *Nomada* bees.** The Gerth and Bleidorn (2016) analyses of nuclear and *Wolbachia* genomes across the bee clade (*N. ferruginata*, (*N. panzeri*, (*N. flava*, *N. leucophthalma*))) provided additional putative examples of *Wolbachia*-host codivergence. Based on the concordant phylogenies of the hosts and their *Wolbachia*, Gerth and Bleidorn (2016) conjectured codivergence across the three host speciation events. However, very different *Wolbachia* rate estimates follow from different subsets of their data. The lowest estimate based on the sister species (*N. flava*, *N. leucophthalma*) that diverged about 1 MYA, is only $5.71 \times 10^{-11}$ substitutions per third site per year. Both this point estimate and the inferred temporal variation in rates seem incompatible with existing data on rates of molecular evolution. Evolutionary rates estimated over shorter time intervals tend to exceed estimates from larger intervals. This was documented for mtDNA in vertebrates by Ho *et al*. (2005), who noted a marked increase in substitution rates estimated over time intervals less than 2 MY. In light of this "recent acceleration" pattern, the extreme apparent slowdown, discussed below, of *Wolbachia* molecular evolution relative to host nuclear and mtDNA over less than two MY suggested by the *Nomada* data in Table 3 is striking.

The first seven rows of Table 3 summarize data describing the divergence of *Wolbachia*, nuclear loci and mtDNA loci across proposed examples in which *Wolbachia* codiverged with their *Nasonia* and nematode hosts. Particularly notable was the relative consistency of the ratios of *Wolbachia* versus nuclear sequence divergence. The next six rows show the comparable data for the *Nomada* species studied by Gerth and Bleidorn (2016). When the outgroup species, *N. ferruginata*, and its *Wolbachia* are compared pairwise to the nuclear genomes and *Wolbachia* of the three ingroup species, (*N. panzeri*, *N. flava* and *N. leucophtalma*), the ratios of *Wolbachia* to nuclear divergence are lower, but within a factor of three of the ratios estimated from *Nasonia* and *Brugia*. In contrast, the pairwise divergences estimated within the ingroup (*N. panzeri*, *N. flava* and *N. leucophtalma*) suggest a relative *Wolbachia* to nuclear divergence rate that is about five to ten times slower than the comparisons of *N. ferruginata* to the three ingroup species. Most dramatically, the comparison of the sister species *N. flava* and *N. leucophtalma* suggested that the relative rate of *Wolbachia* to nuclear divergence between this pair is more than ten times slower than estimated by comparing either sister to *N. ferruginata*. Unlike the *Wolbachia*-nuclear comparisons, the estimated ratios of host mtDNA to nuclear divergence were slightly higher for the sisters (*N. flava*, *N. leucophtalma*) than when either is compared to *N. ferruginata*. Hence, cladogenic inheritance of *Wolbachia* throughout this four-species clade would imply a greater than ten-fold slowdown of *Wolbachia* divergence between *N. flava* and *N. leucophtalma* versus the outgroup *N. ferruginata*, while their mtDNA continued to diverge at a rate typical for the four-species clade as a whole. Such a dramatic recent slowdown of *Wolbachia* divergence seems implausible given the relative constancy over a much longer timescale in the nematodes.

The estimated *Wolbachia* substitution rates based on the *N. ferruginata* comparisons agrees with that obtained from the *Brugia* pair. Assuming codivergence across the four-species clade requires a dramatic slowdown of *Wolbachia* evolution relative to the steady rates of change for mtDNA and nuclear loci. Given that only evidence for codivergence is concordant host and *Wolbachia* phylogenies, we suggest that horizontal transmission of *Wolbachia* across the three-species ingroup is more likely.

**Applying nematode-based calibrations to *w*Mel-like *Wolbachia***

***w*Zts appears to be the mostly closely related *Wolbachia* to *w*Mel in *D. melanogaster.*** Our analysis of the 20 *w*Mel-like *Wolbachia* identified *w*Zts in *Z. tsacasi* as the most closely related variant to *w*Mel in *D. melanogaster* (Fig. S1). Using our nematode-based calibrations, we estimated these *Wolbachia* diverged about 67–924 KYA (Table 5). Following our analyses in the main text, we were made aware of a *Wolbachia* variant identified by Schloz *et al.* (2020) in the robber fly *Holocephala fusca* that was 99.52% identical to *w*Mel across 316 core genes (their Fig. 2A). To determine if this variant is more closely related to *w*Mel than *w*Zts, we obtained Illumina libraries for the *D. melanogaster* lines DGRP256, EZ25, RG34, UG7, DGRP822, CO10N, ED10N, EZ2, DGRP335 and DGRP338 from Richardson *et al.* (2012). These genotypes span the *D. melanogaster* mtDNA haplotypes I–IV and VI. (No genotypes with haplotype V carry *w*Mel.) Libraries for these lines were aligned to the *D. melanogaster* and *w*Mel references with minimap2 (Li 2018), and consensus sequences were produced for each *w*Mel variant using samtools. We used the pipeline and criteria described in the main text to extract genes of single-copy and equal length from the *w*Mel variants, *w*Zts, the *Wolbachia* of *H. fusca*, *w*Yak, and *w*Inc. We estimated a phylogram using 370 genes (355,836 total bp) and the substitution model GTR + I + $\Gamma$ [7,7], as described in the main text.

We found that *w*Zts is most closely related to a clade that includes all *w*Mel variants from the different *D. melanogaster* genomes described above (Fig. S1). Across our 370 genes, *w*Zts-*w*Mel divergence (0.08%) is half that observed between *w*Mel and *Wolbachia* in *H. fusca* (0.16%). In contrast to the results of Schloz *et al.* (2020) based on 316 genes, we found that the *Wolbachia* from *H. fusca* is even more distantly related to *w*Mel than is *w*Yak. The topology of our phylogram is fully consistent with the results of our larger analysis of *w*Mel-like *Wolbachia* presented in Fig. 1A. While we expect additional insect sampling and sequencing will reveal additional *w*Mel-like variants, *w*Zts appears to be the closest known relative to *w*Mel in *D. melanogaster*.

**Cytoplasmic incompatibility is commonly produced by *w*Mel-like and *w*Ri-like *Wolbachia***

We focused on the phylogenetic distribution of CI-inducing *Wolbachia* in our study because CI is the most common *Wolbachia* effect on host reproduction (Shropshire *et al.* 2020; Turelli *et al.* 2022), and because *cif* genes that cause it have been identified (LePage *et al.* 2017; Beckmann *et al.* 2017; Shropshire *et al.* 2018) (Fig. 5A). Of the 20 *w*Mel-like and 8 *w*Ri-like *Wolbachia* in our study, all but 11 *w*Mel-like strains have been tested for CI. Host stocks were available for us to assess CI for only 2 of these 11 *w*Mel-like strains (*w*Seg in *D. seguyi* and *w*Bocq in *D. bocqueti*). Putatively incompatible conspecific crosses between females without *Wolbachia* and males with *Wolbachia* (IC) produced lower egg hatch than did conspecific compatible crosses (CC) for both *w*Seg in *D. seguyi* (IC egg hatch = 0.34 ± 0.21 SD, $N = 14$; CC egg hatch = 0.89 ± 0.11 SD, $N = 18$; $P < 0.001$) and *w*Bocq in *D. bocqueti* (IC egg hatch = 0.18 ± 0.24 SD, $N = 13$; CC egg hatch = 0.60 ± 0.30 SD, $N = 17$; $P = 0.002$). This confirms relatively intense CI for two new *w*Mel-like *Wolbachia*. In total, 8 *w*Mel-like and 6 *w*Ri-like *Wolbachia* in our study cause CI, 2 *Wolbachia* from each clade do not cause detectable CI, 9 *w*Mel-like *Wolbachia* remain uncharacterized, and 1 (*w*Bor) causes male killing (Fig. 5A; see Table S6 for references).

#### Analyzing Cif protein diversity

The Cifs in *w*Mel-like and *w*Ri-like genomes vary in protein sequence, length, and domain composition. While putatively intact *w*Mel-like CifA proteins are relatively similar in size, ranging from 344 (CifA$_{w\mathrm{Zta[T4]}}$) to 493 (CifA$_{w\mathrm{Dal[T1\text{-}2]}}$) amino acids (aa), CifB proteins range from 546 (CifB$_{w\mathrm{San[T4]}}$) to 4433 aa (CifB$_{w\mathrm{Sbr[T5]}}$) (Fig. S7A). CifB$_{\mathrm{[T5]}}$ copies are especially large, as exemplified by the shortest CifB$_{\mathrm{[T5]}}$ copy (CifB$_{w\mathrm{Tri[T5]}}$ = 2833 aa) being 1304 aa longer than the shortest CifB copy across CifB Types in our study (CifB$_{w\mathrm{Zta[T1\text{-}2]}}$ = 1529 aa) (Fig. S7B). We also observe size variation within Cif Types, as exemplified by CifB$_{w\mathrm{Zta[T1\text{-}2]}}$ (*i.e.,* the second CifB$_{\mathrm{[T1]}}$ copy found in the *w*Zta genome). CifB$_{w\mathrm{Zta[T1\text{-}2]}}$ is 357 amino acids longer than any other CifB$_{\mathrm{[T1]}}$ copy and shares at most 51% sequence identity across the alignable region (Fig. S7B; Table S10). As observed by others (Lindsey *et al.* 2018; Martinez *et al.* 2021), CifA homologs lacked confident structural-homology-based domain annotations ($P > 0.8$, see Söding *et al.* 2005 for a description of HHpred homology detection and structure prediction) (Fig. S7A). In contrast, all CifB homologs encoded a PD-(D/E)XK nuclease (Kaur *et al.* 2024), including CifB$_{w\mathrm{Zta[T1\text{-}2]}}$ described above (Fig S7B). CifB$_{\mathrm{[T1]}}$ encodes a Ulp1 deubiquitilase, and CifB$_{\mathrm{[T5]}}$ uniquely encodes OTUB2 deubiquitilase, R12E2.13-like mammalian-wide interspersed repeat, BurrH DNA-binding, Latrotoxin_C toxin, and tetratricopeptide repeat domains (Fig. S7B). The larger size of CifB relative to CifA – and specifically the size of CifB$_{\mathrm{[T5]}}$ – is largely due to the presence of these domains.

Across intact copies of the three Cif Types observed in *w*Mel-like and *w*Ri-like genomes (Cif$_{\mathrm{[T1,T2\ and\ ,T4]}}$), CifA proteins are significantly more similar than CifB from the same pairs ($N = 28$), in both sequence (95% BCa confidence intervals: $\mathrm{ID_{CifA}} = 0.78 – 0.86$, $\mathrm{ID_{CifB}} = 0.69–0.79$; $P = 0.0002$) and AlphaFold structure ($\mathrm{TM_{CifA}} = 0.89 – 0.92$, $\mathrm{TM_{CifB}} = 0.56–0.62$, $P < 10^{-10}$) (Fig. S6A). Fig. S6B and C present similar results for the 12 *w*Mel-like and 6 *w*Ri-like Cif$_{\mathrm{[T1]}}$ proteins, respectively, that comprise the data presented in Fig. 6A.

#### Estimating *Wovirus* and *cif* turnover in *w*Mel-like and *w*Ri-like *Wolbachia*

***cif* operons are common among *w*Mel-like and *w*Ri-like *Wolbachia*.** *Wolbachia* genomes often contain multiple *cif* operons from different described clades (*i.e.,* Types) (LePage *et al.* 2017; Bonneau *et* *al.* 2018; Lindsey *et al.* 2018; Martinez *et al.* 2021; Turelli *et al.* 2018). Excluding *w*Mel-like *w*Au and *w*Tro that do not cause CI (Turelli and Hoffmann 1995; Hoffmann *et al.* 1996; Martinez *et al.* 2015), the *Wolbachia* genomes in our analyses contained between one and three *cif* operons that spanned four *cif* Types (Fig. 5B; Table S9). All genomes except *w*Sbr contained a *cif*$_{[T1]}$ operon, eight contained a *cif*$_{[T2]}$ operon, four contained a *cif*$_{[T4]}$ and five contained *cif*$_{[T5]}$ operons (Fig. 5B). While *cif*$_{[T2]}$ and *cif*$_{[T5]}$ operons are observed in both *w*Mel-like and *w*Ri-like genomes, *cif*$_{[T4]}$ operons are observed only in the *w*Mel-like *w*SYTZ clade. The closely related *w*Ri-like variants *w*Suz and *w*Spc do not cause CI (Hamm *et al.* 2014; Cattel *et al.* 2016), but each carry a *cif*$_{[T1]}$ operon and at least one *cif*$_{[T2]}$ operon (Turelli *et al.* 2018). Finally, the male-killing *w*Bor variant has both *cif*$_{[T1]}$ and *cif*$_{[T5]}$ operons (Sheeley and McAllister 2009). In the main text, we demonstrate that rapid *cif* turnover underlies the diversity of *cif* complements observed in these *Wolbachia* genomes.

**Nearly identical *cifs* span relatively deep sr3 phylogenetic divergences.** Given patterns of rapid *Wovirus* and *cif* turnover presented in the main text, we attempted to associate *cifs* with *Wovirus* typing alleles and compare their respective phylograms. To approximate *Wovirus-cif* associations, we first identified the serine recombinase (sr) typing allele (see main text; Bordenstein and Bordenstein 2022) nearest to a *cif* present on the same contig. We then quantified the distance between them. Table S16 presents these results. We observed 33 *cif* alleles that are closest to an sr3WO (sr3) allele, including 22 *cif*$_{[T1]}$ alleles. 21 of these *cifA*$_{[T1]}$ alleles are separated by 14K bp or fewer from their nearest sr3 (range: 7,477–13,770 bp), while the next closest is over 200 kbp away. We focus our analyses on these 21 cases from 12 *w*Mel-like and 8 *w*Ri-like *Wolbachia*, including *w*Ri that has two *cifA*$_{[T1]}$ alleles associated with distinct sr3 alleles. In each case the sr3 allele is physically located on the 3' end of the associated *cif*$_{[T1]}$ operon, and in closer physical proximity to *cifB*$_{[T1]}$. Only 1,900 bp separate *cifB*$_{[T1]}$ and sr3 alleles in *w*Mel, *w*Inc, and in all 9 *w*Ri-like examples, with a single gene (an ankyrin domain protein) between them. In the *w*Mel-like *w*Ach and *w*Bocq genomes, 3,814 bp separate *cifB*$_{[T1]}$ and sr3 alleles, with two genes (annotated as hypothetical proteins) between them. The sr3 and *cifB*$_{[T1]}$ alleles in the *w*Dal genome are separated by the largest physical distance (10,194 bp), with eight genes between them. Table S17 presents these results.

We next tested for intralocus recombination using GARD and the three Phipack tests. GARD identified two sr3 breakpoints that were supported by the three PhiPack tests ($P < 0.01$), which we verified by visually inspecting the alignment. This corresponded to three partitions (P) (P1: 1–471, P2: 472–732, and P3: 733–1503) (Fig. S3). For *cifA*$_{[T1]}$, GARD also identified *cifA*$_{[T1]}$ recombination in analysis of the dataset presented in Figure 5C. For these analyses, issues with GARD recombination calls did not emerge, whereby a lack of variation may lead GARD to resolve three or more groups randomly. Across partitions, we observed sufficient pairwise differences (minimum pairwise difference: P1 = 49, P2 = 29, and P3 = 83). Fig. S4 illustrates that differences in *cifA*$_{[T1]}$ partition phylograms are related to the placement of four outgroup *cifA*$_{[T1]}$ alleles. Among these four alleles, only two (*w*Zta and *w*Ara) *cifA*$_{[T1]}$ alleles are plausibly associated with sr3 based on our criteria.

Several patterns are worth noting from our analyses. The *w*Zta and *w*Ara *cifA*$_{[T1]}$ alleles are distantly related to a clade that includes the 19 other very closely related *cifA*$_{[T1]}$ alleles (Fig. S5). Among these 19, 9 *w*Ri-like *cifA*$_{[T1]}$ alleles differ from each other by at most 2 bp, the *w*Mel-like *w*Ach allele differs from *w*Ri-like alleles by at most 22 bp, and the remaining 8 *w*Mel-like alleles differ from the *w*Ri-likes by at most 5 bp. The 9 *w*Ri-like *cifA*$_{[T1]}$ alleles are plausibly associated with 9 identical sr3 alleles. Based on our nematode calibrations, *w*Ri-like *Wolbachia* core genomes diverged about 14–218 KYA. It is plausible that this sr3 and associated *cifA*$_{[T1]}$ were present in the *w*Ri-like MRCA. In contrast, the 19 very closely related *cifA*$_{[T1]}$ alleles span relatively deep phylogenetic divergences among the sr3 alleles with which they are plausibly associated (Fig. S5). Some *w*Mel-like sr3 alleles are more closely related to the 9 *w*Ri-like sr3 alleles than they are to other *w*Mel-like sr3 alleles; however, the specific alleles and their placements vary across sr3 partitions (Fig. S3). For partition 1 of sr3, the *w*Mel and *w*Zts alleles were identical to all *w*Ri-like alleles, while the *w*Inc allele differed at only 1 site. In partition 2, these 3 alleles and the *w*Bocq and *w*Ach alleles form a clade that is sister to a clade of all *w*Ri-like alleles, with 7 other *w*Mel-like alleles forming a more distantly related clade. Finally, in partition 3, the *w*Mel, *w*Inc, and *w*Zts alleles and 5 other *w*Mel-like alleles are more distantly related to the clade of *w*Ri-like alleles than are 4 other *w*Mel-like alleles. This supports a history of sr3 recombination between *w*Mel-like and *w*Ri-like copies.

Given the similarity of *cifA*$_{[T1]}$ alleles and the history of recombination of sr3, tests for topological concordance were not possible; however, many very closely related and nearly identical *cifA*$_{[T1]}$ alleles spanning relatively deep sr3 divergences supports that mechanisms other than *Wovirus* transfers plausibly contribute to the patterns we observed. These include *Wovirus* recombination (*e.g.,* Bordenstein and Wernegreen 2004), and transfers involving plasmids (*e.g.,* Martinez *et al.* 2022) or transposons (*e.g.,* Gillespie *et al.* 2018; Cooper *et al.* 2019; Amoros *et al.* 2025a). For example, *cif* transfers involving IS elements are supported in multiple *Wolbachia* systems (Cooper *et al.* 2019; Madhav *et al.* 2020), in addition to IS5-mediated *cif* transfer observed in *Orientia tsutsugamushi* (Oswalt *et al.* 2025).

***cif* transfers involving IS elements.** IS-mediated transfer was first identified as a plausible mechanism involved in the acquisition of *cif*$_{[T4]}$ loci by *Wovirus* observed in *w*Yak and closely related *w*Mel-like genomes (Cooper *et al.* 2019). A single sr3WO *Wovirus* in *w*SYT contains *cif*$_{[T1]}$ (single copy) and *cif*$_{[T4]}$ operons (one copy in *w*Tei and two in *w*SY, Baião *et al.* 2021), whereas the sr3WO in closely related *w*Mel (and *w*Zts) in *D. melanogaster* does not contain a *cif*$_{[T4]}$ operon (Fig. 5B; Cooper *et al.* 2019). An intact 5′ ISWpi1 flanks the *cif*$_{[4]}$ loci in *w*SYT (Cooper *et al.* 2019; Baião *et al.* 2021). Comparison of *cif*$_{[T4]}$ operons in divergent *w*Yak and in supergroup B *w*Pip (*w*Yak-*w*Pip MRCA: CPR: 22–359 MYA) genomes found less than 3% divergence between these operons. In contrast, the surrounding phage genes between *w*Yak and *w*Pip are ~15% diverged (see Cooper *et al.* 2019, Fig. 6), supporting a common *cif*$_{[T4]}$ donor that is distinct from the phages that currently contain them. While higher quality genomes confirmed the placement of ISWpi1 5' of *cif*$_{[4]}$ loci in *w*SYT, ISWpi1 placement differed at 3' locations among *w*SYT (Baião *et al.* 2021). Variation in 3′ IS placement can plausibly be explained by subsequent *Wovirus* recombination and IS turnover that frequently reposition or delete IS elements following insertions (Cordaux *et al.* 2008). Supporting this interpretation, related *Wovirus* regions show high structural plasticity, numerous partial IS remnants, and strain-specific junctions consistent with secondary rearrangements (Wu *et al.* 2004; Klasson *et al.* 2009; Baião *et al.* 2021; Bordenstein and Bordenstein 2022; Vancaester and Blaxter 2023). Notably, IS elements flank many of the *cifs* in our analyses (Table S15), and among those with IS elements within 10kb, 13 of 41 have a both 5' and 3' transposases, 37 of 41 have a 5' transposase, and 17 of 41 have a 3' transposase (Table S15). More broadly, transposon involvement in *cif* transfers is supported in *Wolbachia* (*e.g.,* Amoros *et al.* 2025a) and other endosymbiont systems (Oswalt *et al.* 2025; Amoros *et al.* 2025b).

The presence of $cif_{[T4]}$ operons in *w*SYTZ, and their absence in all other *w*Mel-like *Wolbachia* we studied (Fig. 5B), including closely related *w*Mel and *w*Zts, implies IS-mediated transfer of $cif_{[T4]}$ loci into *Wovirus* observed in *w*SYTZ occurred after they diverged from (*w*Mel, *w*Zts). Loss of these loci by both *w*Mel and *w*Zts seems unlikely (cf. Baião *et al.* 2021). *Z. taronus* and *D. santomea* co-occur at high altitudes on Pico de São Tomé where we sample them, and it is plausible that species interactions mediated horizontal transfer of *w*Zta into the *D. yakuba* clade, followed by bouts of hybridization and introgression between *D. yakuba*-clade species that spread $cif_{[T4]}$-carrying *Wolbachia* among these hosts. Indeed, joint analysis of mtDNA supported horizontal *Wolbachia* acquisition by *D. yakuba*-clade hosts from an unknown donor (see our main text; Cooper *et al.* 2019), and *Z. taronus* is a plausible donor host.

### Estimating selection on *cifs*

In the main text we report that putatively intact copies of $cifA_{[T1]}$ are constrained by selection, with weaker stabilizing selection maintains intact $cifB_{[T1]}$. Our analyses support that the $cifB_{[T1]}$ Nuc2 domain is particularly constrained. Here we present detailed results for the SWAKK analyses that were summarized in the main text. We then provide additional details related to codeml analyses in the main text, including all results related to selection on $cifA_{[T1]}$ copies with open-reading frame (ORF) disrupting mutations.

**SWAKK analyses on intact $\mathbf{Cif_{[T1]}}$.** Pooled medians for $\omega$ across all windows were lower for $CifA_{[T1]}$ (0.29) than $CifB_{[T1]}$ (0.37), and the mean of the per-pair differences of medians on the log scale was significantly below 0 ($\Delta_p = -0.23$; BCa 95% CI: −0.38, −0.08; $P < 0.0002$). This difference in $\omega$ corresponded to a geometric-mean $CifA_{[T1]}/CifB_{[T1]}$ $\omega$ ratio of 0.80 (BCa 95% CI: 0.68–0.92), implying that the typical window-level $CifA_{[T1]}$ $\omega$ is 20% lower than for $CifB_{[T1]}$. We next compared $\omega$ for $CifB_{[T1]}$ domains to $\omega$ for the pooled non-domain $CifB_{[T1]}$ regions. While $\omega$ for Nuc1 ($\Delta_p = -0.20$; BCa 95% CI: −0.38, 0.05; $P = 0.08$) and Dub ($\Delta_p = -0.07$; BCa 95% CI: −0.30, 0.27; $P = 0.28$) domains trended lower than $\omega$ for the non-domain background, the differences were not statistically significant. In contrast, for the Nuc2 domain we observed that $\Delta_p$ was significantly below 0 ($\Delta_p = -0.38$; $P = 0.003$), despite its BCa interval overlapping with 0 (BCa 95% CI: −0.75, 0.15). Taken together, SWAKK results assessed in 3D sliding-window space broadly agreed with those based on maximum-likelihood models presented in the main text.

**codeml analyses: intralocus recombination.** Prior to implementing tree-based codeml analyses, we tested for intralocus recombination. GARD did not identify breakpoints for intact *cifB*$_{[T1]}$. For intact *cifA*$_{[T1]}$, GARD identified one breakpoint corresponding to codon 146 in the *w*Mel *cifA*$_{[T1]}$ copy. We label partition 1 as codons 1–146 and partition 2 as codons 147–475, with topologies ((*w*Zta, *w*Spa), (*w*Mel, *w*Dal)) (RAxML bootstrap support = 87%) and ((*w*Dal, *w*Zta), (*w*Mel, *w*Spa)) (RAxML bootstrap support = 97%), respectively. For disrupted *cifA*$_{[T1]}$, GARD identified one breakpoint corresponding to codon 87 in the *w*Mel *cifA*$_{[T1]}$ copy. We label partition 1 as codons 1–87 and partition 2 as codons 88 – 474, with topologies ((*w*Bic, *w*Ach),(*w*Stv, *w*NFa)) (RAxML bootstrap support = 100%) and ((*w*Bic, *w*Stv), (*w*NFa, *w*Ach)) (RAxML bootstrap support = 100%), respectively. In all these cases where GARD identified recombination, Phipack tests were statistically significant ($P < 0.01$). We separately analyzed each statistically supported partition in our tree-based analyses.

**codeml analyses: expanded presentation of main text results.** The following results for intact *cifA*$_{[T1]}$ are an expanded presentation of results provided in the main text. The one-ratio model estimated an average $\omega = 0.22$ for partition 1 (sites 1–146), with evidence for significantly heterogeneity in $\omega$ across sites (M1a vs. M0: LRT = 32.6, df = 1, $P < 0.0001$). However, models adding a positively selected site class did not improve model fits (M2a vs. M1a: LRT = 0.42, df = 2, $P = 0.81$; M8 vs. M7: LRT = 3.5, df = 2, $P = 0.17$). The one-ratio model estimated an average $\omega = 0.38$ for partition 2 (sites 147–475) that was more similar to $\omega$ for *cifB*$_{[T1]}$ than to partition 1 $\omega$. Strong purifying selection on the N-terminus region agrees with observations and interpretations provided in our main text. Again, the “nearly neutral” model fit significantly better with evidence of heterogeneity in $\omega$ across sites (M1a vs. M0: LRT = 38.3, df = 1, $P < 0.0001$). In contrast to partition 1, adding an $\omega > 1$ site class marginally improved the fit (M2a vs. M1a: LRT = 5.82, df = 2, $P = 0.053$), while the beta plus positive-selection model provided a significantly better fit and identified a small (3.2%) positively selected class (M8 vs. M7: LRT = 7.36, df = 2, $P = 0.025$). This included two codons with BEB posterior $P \geq 0.95$ (positions 171 and 390) identified only by the beta plus positive-selection model. Similar to *cifB*$_{[T1]}$, visual inspection of the codon alignment revealed MNM changes that violate the codon substitution model. In summary, these results indicate relatively stronger constraint on partition 1 of *cifA*$_{[T1]}$, substantial heterogeneity in $\omega$ across each partition, and little support for positively selected *cifA*$_{[T1]}$ sites in either partition.

**codeml analyses: selection on putatively disrupted *cifA*$_{[T1]}$.** To test the hypothesis that disrupted copies are under relatively relaxed selection, we compared $\omega$ for putatively intact *cifA*$_{[T1]}$ and putatively disrupted *cifA*$_{[T1]}$ copies using pairwise contrasts in codeml. While the HL median difference (*i.e.*, $\omega_{cifA\text{[T1]-intact}} - \omega_{cifA\text{[T1]-disrupted}}$) was negative (−0.03), the 95% CIs (−0.08, 0.03) overlapped with 0 so the difference was not statistically significant (Wilcoxon $W = 11$, $P = 0.15$). Bayesian bootstrap posteriors indicate with $P = 0.85$ that disrupted $\omega$ is lower, *i.e., P*($\Delta < 0$) where $\Delta = \omega_{cifA\text{[T1]-intact}} - \omega_{cifA\text{[T1]-disrupted}}$.

We conclude that while Wilcoxon test statistics were nonsignificant, our tests are limited by small sample sizes and future analyses may demonstrate relaxed selection on disrupted copies of *cifA*.

We next evaluated six models in codeml that differ in their treatment of variation in $\omega$ across lineages and across sites (see our Materials and Methods). For partition 1 of disrupted $cifA_{[T1]}$, the one-ratio model estimated an average $\omega = 0.265$, trending higher than for partition one of intact $cifA_{[T1]}$, and the branch model did not provide a significantly better fit (branch vs. M0: LRT = 6.24, df = 3, $P = 0.10$). In site-model analyses, the nearly neutral model significantly improved (M1a vs. M0: LRT = 14.32, df = 1, $P < 0.0001$), indicating heterogeneity in selective pressure across sites. However, neither positive-selection model provided significantly better fits (M2a vs. M1a: LRT = 0, df = 2, $P = 1$; M8 vs. M7: LRT = 0.42, df = 2, $P = 0.81$). For partition 2, the one-ratio model estimated an average $\omega = 0.39$, and the branch model did not significantly improve model fit (branch vs. M0: LRT = 0.98, df = 3, $P = 0.81$). Relatively strong purifying selection on the N-terminus region agrees with observations and interpretations provided in our main text related to its contributions to CI rescue. In site-model analyses, we observe significant heterogeneity in selective pressure among sites (M1a vs. M0: LRT = 80.12, df = 2, $P < 0.0001$). One of the positive selection models provided a marginally better fit (M8 vs. M7: LRT = 6.2, df = 2, $P = 0.045$) but the other did not (M2a vs. M1a: LRT = 4.5, df = 2, $P = 0.11$) with 1.6–2.2% of sites in the $\omega > 1$ class. However, neither model identified codons with BEB $P \geq 0.95$. In summary, there was significant heterogeneity in selective pressure across sites within both partitions of disrupted $cifA_{[T1]}$, with purifying selection constraining both partitions but with lower $\omega$ for the N-terminus region. Evidence for positive selection was restricted to partition two, but no sites with BEB $P \geq 0.95$ were found.

### Supplementary Figures

#### Figure S1

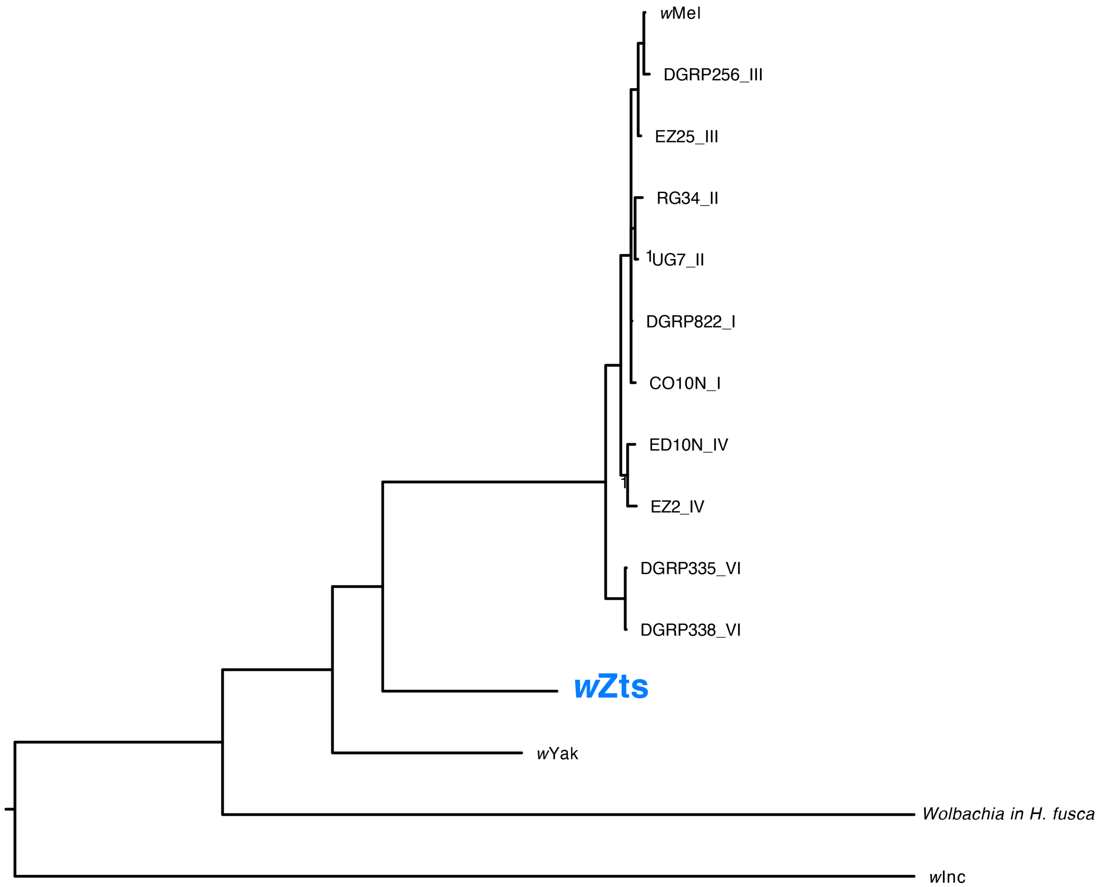


**Fig. S1. Phylogram of focal *w*Mel-like *Wolbachia*, focusing on the placement of *w*Zts relative to *w*Mel.** The phylogram includes the *w*Mel reference strain, *w*Mel variants from 10 *D. melanogaster* genotypes that differ in their mtDNA haplotypes, *w*Zts from *Z. tsacasi*, *w*Yak from *D. yakuba*, *Wolbachia* in *H. fusca*, and *w*Inc in *D. incompta*. Nodes with posterior support < 0.95 were collapsed into polytomies. All *w*Mel variants from the 11 *D. melanogaster* genotypes form a clade that is sister to *w*Zts, establishing that *w*Zts is much more closely related to *w*Mel than is the *Wolbachia* from *H. fusca* discussed in Scholz *et al*. (2020).

**Figure S2**

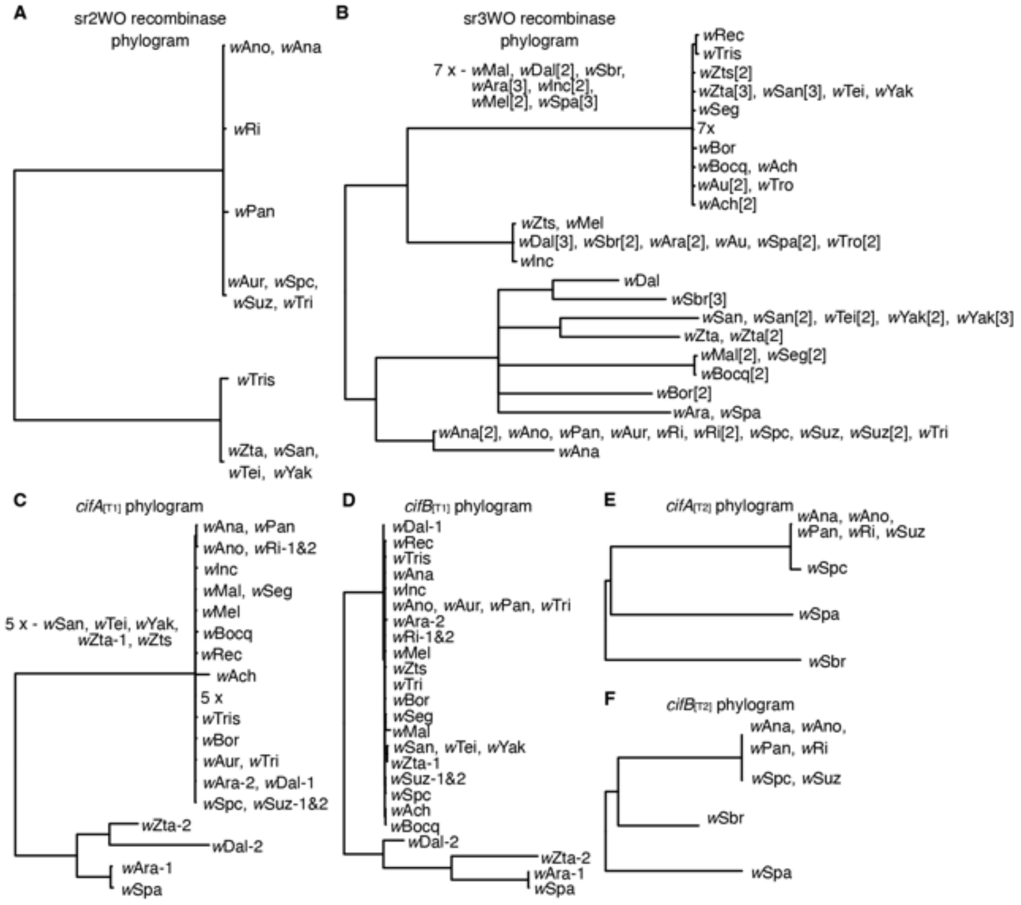


**Fig. S2. Phage and *cif* phylograms.** Phylograms for (**A**) serine recombinase homologs of the sr2WO phage WOVitA, and (**B**) serine recombinase for homologs of the sr3WO phage WOMelB. Brackets on tips denote different copies observed in a genome. Phylograms for (**C**) *cifA*$_{[T1]}$, (**D**) *cifB*$_{[T1]}$, (**E**) *cifA*$_{[T2]}$, and (**F**) *cifB*$_{[T2]}$. In panels C and D, dashes are used to denote specific copies observed in a genome and correspond to those presented in Figs. 5 and S4. Identical sequences were collapsed into single tips and nodes with posterior probability < 0.95 were collapsed into polytomies. Genes were not partitioned by codon position. The difficulty in visually distinguishing which nodes tiny branches subtend in panels C and D reflects the high similarity of many *cif*$_{[T1]}$ copies observed in *w*Mel-like and *w*Ri-like genomes (see main text).

**Figure S3**

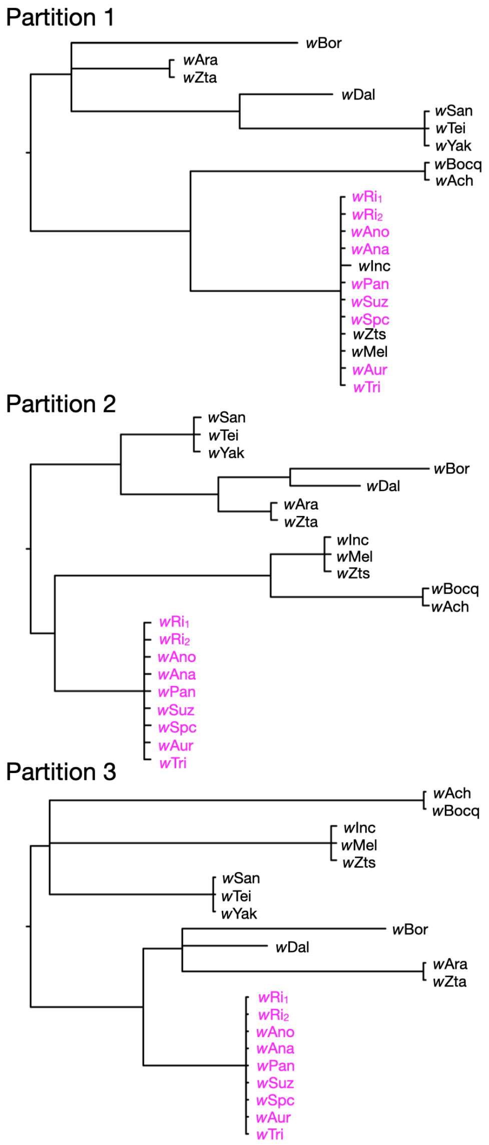


**Fig. S3. Phylograms for three sr3 partitions based on GARD analysis of intralocus recombination.** Data include only the sr3 copies plausibly associated with $cif_{[T1]}$ operons (see Fig. S5; Tables S16 & S17). GARD identified two breakpoints, corresponding to three sr3 partitions. A phylogram for each partition is presented. Nodes with posterior probability < 0.95 are collapsed into polytomies. Alleles from *w*Ri-like *Wolbachia* are colored in pink. Subscripts for *w*Ri represent different sr3 copies within the same *Wolbachia* genome. Table S16 presents additional details for sr3 copies and associated $cif_{[T1]}$ operons (entries in bold). Fig. S5 presents a phylogram for sr3 unpartitioned.

**Figure S4**

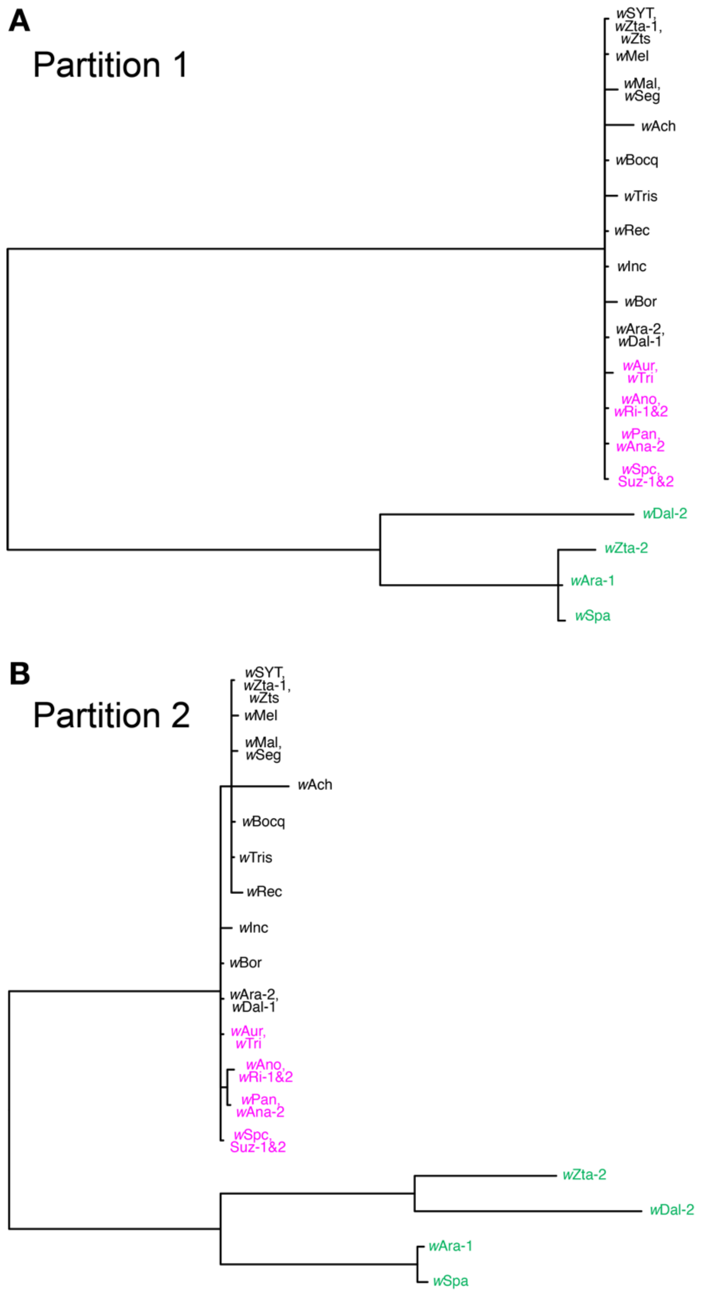


**Fig. S4. Phylograms for two *cifA*$_{[T1]}$ partitions based on GARD analysis of intralocus recombination.** All *w*Ri-like alleles (pink) and 16 *w*Mel-like alleles (black) comprise a clade of nearly identical copies in both partitions. In contrast, the *cifA*$_{[T1]}$ alleles from four other *w*Mel-like variants comprise a more distantly related clade in each partition (green). The relationships of these four distantly related alleles differ between partitions, with a copy in *w*Zta being more closely related to a copy in the *w*Dal genome in Partition 2, but more closely related to copies in *w*Ara and *w*Spa genomes in Partition 1. Nodes with posterior probability less than 0.95 were collapsed into polytomies. Dashes are used to denote specific copies observed in a genome and correspond to those presented in Figs. 5 and S2.

**Figure S5**

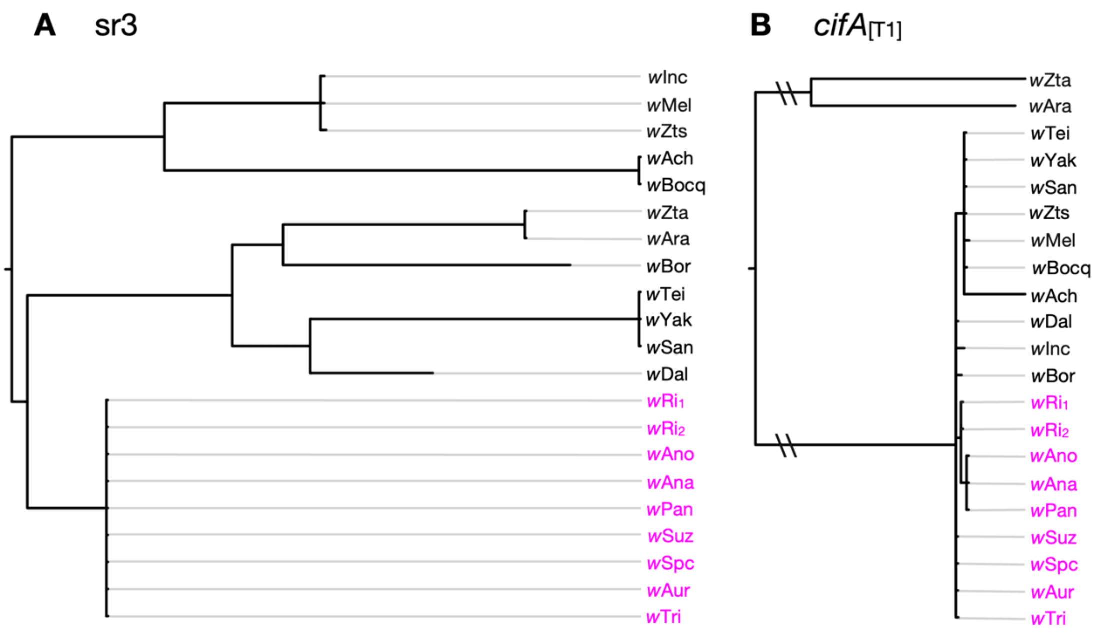


**Fig. S5. Nearly identical *cifA*$_{[T1]}$ alleles span relatively deep phylogenetic divergences among plausibly associated sr3 alleles.** Phylograms for **(A)** unpartitioned sr3 alleles and **(B)** *cifA*$_{[T1]}$ alleles plausibly associated with these sr3 alleles (Tables S16 & S17). Branches leading to the two sets of closely related *cifA*$_{[T1]}$ alleles are shortened (\\) to improve visualization. Excluding *w*Zta and *w*Ara, we observe at most 22 differences between the *cifA*$_{[T1]}$ alleles, or at most 5 differences excluding *w*Ach and *w*Bocq. In contrast, associated sr3 alleles span relatively deep phylogenetic divergences. Nodes with posterior probability $< 0.95$ are collapsed into polytomies. Alleles from *w*Ri-like *Wolbachia* are colored in pink. Subscripts for *w*Ri represent different sr3 and *cif*$_{[T1]}$ copies within the same *Wolbachia* genome. Table S16 presents additional details for sr3 copies and associated *cif*$_{[T1]}$ operons (entries in bold). Fig. S3 presents phylograms for GARD-based sr3 partitions.

**Figure S6**

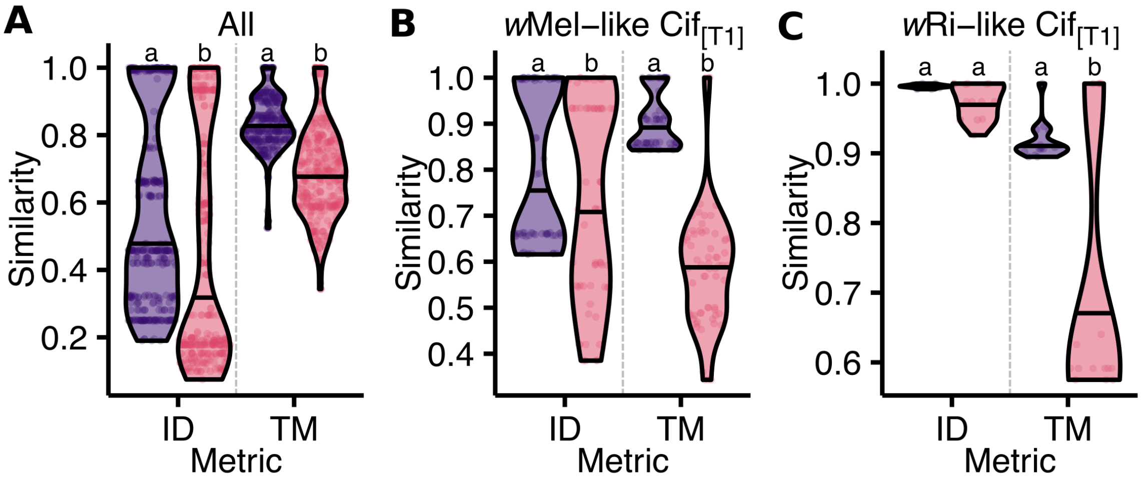


**Fig. S6. Cif protein sequence identity and structure similarity.** The relationship between CifA (purple, left) and CifB (pink, right) pairwise sequence identity (ID) and structure similarity (TM) compared across (**A**) intact *w*Mel-like and *w*Ri-like CifA$_{[T1]}$, CifA$_{[T2]}$, and CifA$_{[T4]}$ pairs (Table S10, $N = 28$); and (**B**) intact Cif$_{[T1]}$ pairs in *w*Mel-like *Wolbachia* genomes ($N = 12$); and (**C**) intact Cif$_{[T1]}$ pairs in *w*Ri-like *Wolbachia* genomes ($N = 6$). Shared letters within pairs represent statistically similar groups determined by a Mann-Whitney U test (2 groups) at $P < 0.05$.

### Figure S7

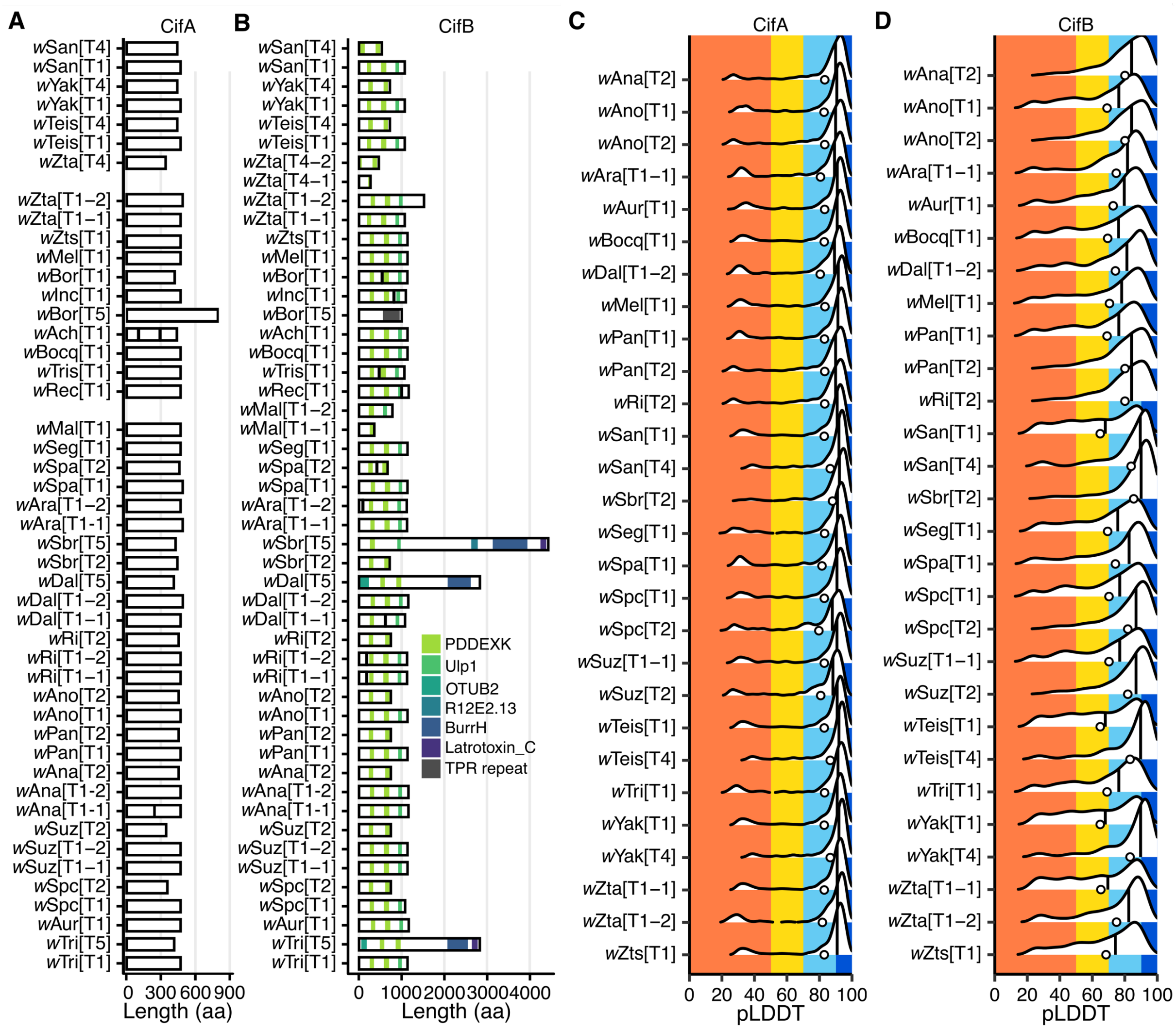


**Fig. S7. *w*Mel-like and *w*Ri-like Cif domain annotations and sequence/structure diversity.** (**A**) CifA protein schematics. HHPred revealed no annotations with > 80% probability (see main text). (**B**) CifB protein schematics. Predicted protein domains (HHpred, >80% probability) are shown for each CifB protein, with domain identities indicated by color as defined in the inset legend. Structural confidence for (**C**) CifA and (**D**) CifB Alphafold structures. pLDDT is calculated per residue. pLDDT of 100 to 90 (dark blue), 90 to 70 (light blue), 70 to 50 (yellow), and 50 to 0 (orange) indicate very high, high, low, and very low confidence, respectively. Ridgeline plots illustrate the range and density of pLDDT values per protein. Vertical lines and dots represent median and mean pLDDT, respectively.

**Figure S8**

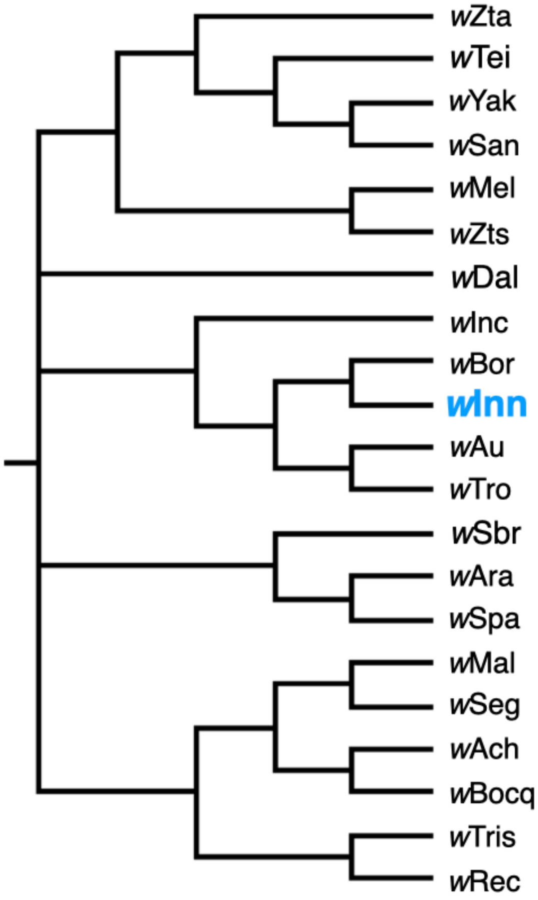


**Fig. S8. Phylogenetic placement of *w*Inn among *w*Mel-like *Wolbachia* based on concatenated single-copy genes.** A cladogram for the *w*Mel-like *Wolbachia* included in our study (Tables 1 & S1), plus *w*Inn from *D. innubila*. Nodes with posterior probability < 0.95 were collapsed into polytomies. Note that several of the basal nodes are not resolved, reflecting rapid diversification of these *w*Mel-like variants. The analysis confirms that *w*Inn is a member of the *w*Mel-like *Wolbachia* clade and is most closely related to *w*Bor. For the second most variable gene in the *w*Mel-like gene set, BLAST revealed the outlier *w*Bor allele shares 99.9% similarity with the homologous allele in *w*Inn.

### Supplementary Table Legends

**Table S1. The *Wolbachia* and hosts used in our study: 20 *w*Mel-like, 8 *w*Ri-like, 4 supergroup A *Nomada*-associated, and 3 supergroup C and 4 supergroup and D nematode-associated variants.** We present taxonomic information and genome accessions for both *Wolbachia* and the hosts that carry them. For each *Wolbachia* genome, we also provide the number of contigs that comprise our assemblies and the number of BUSCO genes found. This number can be used to assess fragmented genome quality by comparison with the complete genomes. Footnotes provide information related to the sources of our genomes and their use in other studies. To estimate the divergence times of some *w*Mel-like *Wolbachia* host species, we used genomic data for *Drosophila* species ($N$ =3 ) that do not carry *w*Mel-like *Wolbachia*. Information related to these species is listed in the final three rows of the table.

**Table S2. Gene-tree concordance factors (gCF) for internal branches of the species tree for seven supergroup A *Wolbachia* variants.** We summarize, for each internal branch (Node) of the reference topology C, the agreement of single-gene trees with that split. For each branch we report gCF (*i.e.,* the fraction of decisive gene trees supporting the reference split) and the two discordance factors (gDF1, gDF2) for the alternative nearest-neighbor resolutions around that branch, together with the number of decisive and uninformative genes. Results are presented for four bootstrap-support thresholds (50%, 60%, 80% and 95%) used to collapse gene-tree nodes prior to scoring decisiveness. Decisive genes are those that contain the required taxa and resolve one of the three alternatives for that split after collapsing low-support branches in the gene tree. Results are shown for $N$ = 442 genes.

**Table S3. ASTRAL quartet frequencies for the species tree of seven supergroup A *Wolbachia*.** For each internal branch, we report the ASTRAL quartet frequencies around that branch (q1, q2, and q3). q1 is the fraction of quartets supporting the species-tree resolution of the branch, while q2 and q3 are the fractions supporting the two alternative nearest-neighbor resolutions. Quartets were computed using weighted ASTRAL from the 442 gene trees for the seven supergroup A *Wolbachia* (Fig. 3).

**Table S4. The number of loci that support each of the 15 possible unrooted trees of the five entities *w*Ha, *w*Bic, *w*Bar, (*w*Mel, *w*Au) and (*w*Ri, *w*Tri).** Of the 52 phylogenetically informative loci described in the main text, 14 support C, the consensus phylogeny described in the main text and presented in Fig. 3. The 14 alternative unrooted topologies, denoted A1–A14, are each supported by 6 or fewer loci. While all 52 loci support (*w*Mel, *w*Au) and (*w*Ri, *w*Tri), 25 support ((*w*Ri, *w*Tri), *w*Ha), namely the loci supporting C and the two most common alternative topologies A1 and A2.

**Table S5. Positions of the 52 phylogenetically informative loci with respect to 4 reference genomes, *w*Mel, *w*Au, *w*Ri and *w*Ha.** We list the genes in their order of occurrence relative to the origin of the *w*Mel reference genome. For each alternative reference genome, we present their order of appearance in that reference genome in parentheses. Differences in gene order illustrate pervasive rearrangements, especially inversions. In the last two columns, we present two measures of gene-specific divergence among the seven genomes: the fraction of polymorphic sites within each gene and the total number of substitutions (estimated by the sum of branch lengths obtained under the GTR+Γ+I substitution model, without partitioning by codon position).

**Table S6. The 20 most polymorphic loci in comparisons of the seven *Wolbachia* variants that span the *w*Mel-like and *w*Ri-like clades, with consensus phylogeny: (((*w*Mel, *w*Au), (*w*Bic, *w*Bar)), ((*w*Ri, *w*Tri), *w*Ha)).** We ranked 438 loci by their fractions of variable sites. Only the most variable locus met our two-fold outlier criteria: RefSeq_WP_010962975.1, a membrane protein. This locus showed about 19% polymorphic sites, with 95 variable sites out of 498 bp.

**Table S7. No recent horizontal acquisition of *Wolbachia* between *D. simulans* and its parasitoid *Asobara tabida*.** Applying a "universal" rate of bacterial evolution and partial *Wolbachia ftsZ* sequences from *D. simulans* and its parasitoid *A. tabida*, Werren *et al*. (1995) inferred recent horizontal *Wolbachia* acquisition. Because *ftsZ* sequences seem to diverge particularly slowly, we assessed differences between these *Wolbachia* across 560 bp of *wsp* coding sequence. We obtained *A. tabida wsp* sequences from Rancès *et al.* (2008), identified the orthologous region in *w*Ri and *w*Mel with BLAST, and aligned them with MAFFT v. 7. We then counted the pairwise differences for each possible comparison. We present these differences here. As reported in the main text, *w*Ri differs from each of the three *Wolbachia* variants found in *Asobara tabida* by at least as much as *w*Ri differs from *w*Mel. *w*Ri and *w*Mel are diverged at least 1.4 MYA, which is comparable to the divergence times of *D. simulans* from *D. sechellia* and *D. mauritiana* (Suvorov *et al*. 2022), precluding recent horizontal acquisition of *Wolbachia* between *D. simulans* and its parasitoid A. *tabida*.

**Table S8. CI is common among *w*Mel-like and *w*Ri-like *Wolbachia*.** We report the CI phenotype observed for *w*Mel-like and *w*Ri-like *Wolbachia* in our study (Fig. 5A), including our discovery of relatively intense CI caused by *w*Mel-like *w*Seg and *w*Bocq variants. Relevant references for previous CI estimates are reported. CI phenotypes with an asterisk denote estimates reported from only transinfected *D. simulans* backgrounds. For the three *w*SYT variants that were initially characterized as not causing CI (Zabalou *et al.* 2004), we provide several references. *w*Tei causes CI that varies in intensity among *D. teissieri* backgrounds (Zabalou *et al.* 2004; Cooper *et al.* 2017) and relatively intense CI in *D. simulans* backgrounds (Zabalou *et al.* 2008; Martinez *et al.* 2015). While the two *w*SY variants can cause CI (Cooper et al. 2017), results have been inconsistent in natural backgrounds (Zabalou *et al.* 2004), including studies that have and have not observed CI in transinfected *D. simulans* backgrounds (Zabalou *et al.* 2008; Martinez *et al.* 2015).

**Table S9. *Woviruses* and *cifs* are common in *w*Mel-like and *w*Ri-like genomes.** We report the number of *Wovirus* copies, the Types of *Woviruses* observed, the number of *cif* copies, and the *cif* Types observed for each *Wolbachia* in our study. For *Wovirus* and *cif* Types observed in each genome, we report the number of copies of each Type in parentheses, following their respective entries. *Wovirus* identification and typing was based on the large serine recombinase (sr) described in the main text.

**Table S10. Cif sequence and structural similarity matrices. Pairwise percent identity of all (A) CifA and (B) CifB proteins reported in this study.** Pairwise percent identity of (C) CifA and (D) CifB proteins included in the structure and sequence similarity analysis. (A–D) Muscle5 was used to generate 100 multiple sequence alignments (MSAs), and the alignment with the highest column confidence was selected for downstream analyses. Pairwise identity was calculated as the percentage of identical sites between each pair in the MSA relative to the alignment length, including gaps. TM-scores from pairwise comparisons of (E) CifA and (F) CifB AlphaFold2 structures.

**Table S11. Codeml pairwise analyses indicate constraint on intact $cifA_{[T1]}$.** Across all pairwise contrasts in codeml, $\omega$ for intact $cifA_{[T1]}$ (range: 0.26–0.38) was consistently and significantly lower than $\omega$ for intact $cifB_{[T1]}$ (Fig. 6B; HL median difference = −0.06; Wilcoxon $V = 0$, $P = 0.02$). While the HL median difference between $\omega$ for intact $cifA_{[T1]}$ and disrupted $cifA_{[T1]}$ ($\omega$ range: 0.27–0.38) copies was negative (−0.03), the 95% CIs (−0.08, 0.03) overlapped with 0 and the difference was not statistically significant ($P = 0.15$).

**Table S12. Results of codeml model fitting for $cif_{[T1]}$ genes.** Models were run on intact $cifB_{[T1]}$ (entire sequence, no partitions), intact $cifA_{[T1]}$ (two codon partitions), and disrupted $cifA_{[T1]}$ (two codon partitions). Columns indicate the gene, partition, codon site range, evolutionary model (M0, Branch, M1a, M2a, M7, and M8), log-likelihood (lnL), and parameter estimates (*e.g.*, ω values and proportions).

**Table S13. Likelihood ratio test (LRT) results from codeml model comparisons for *cif* genes.** Analyses include intact *cifB*$_{[T1]}$ (no partitions), intact *cifA*$_{[T1]}$ (two codon partitions), and disrupted *cifA*$_{[T1]}$ (two codon partitions). Columns report the gene, partition, codon site range, model comparison (*e.g.,* M0 vs Branch, M1a vs M2a, M7 vs M8), log-likelihoods of null and alternative models, test statistic ($2\Delta \ln L$), degrees of freedom (df), and associated *P*-values.

**Table S14. Codon sites supported by Bayes Empirical Bayes (BEB, posterior probability ≥ 0.95) as positively selected in codeml analyses.** Significant sites were identified in intact *cifB*$_{[T1]}$ (entire sequence) and intact *cifA*$_{[T1]}$ (partition 2). Columns report the gene, partition, codon site range, site position, models supporting the site (M2a, M8, or both), observed codons, and whether codon variation was consistent with multinucleotide mutation (MNM)-like patterns.

**Table S15. IS elements flank the majority of *cifs* in our analysis. (Sheet A)** For each of the 50 *cifs* observed across our *w*Mel-like and *w*Ri-like *Wolbachia* genomes, we searched for genes annotated as transposases within 10K bp (5' and 3') on the same contig as a focal *cif.* Distances are calculated as the bp between focal transposase and *cifA* copies. We report the number and identity of genes between each transposase and *cifA* copy that meet our criteria. **(Sheet B)** Summary of the types of transposases found near *cifs* across our genomes and how often they occur.

**Table S16. Many *cif* operons occur on contigs that also contain a nearby *Wovirus* based on sr typing.** We report a complete list of the *cifA* copies (*N* = 50) observed in *w*Mel-like and *w*Ri-like *Wolbachia* genomes. Types (T1-T5) are observed across these *Wolbachia*, with the exception of T3. We report the distances between *cifA* and their nearest sr-typing allele on the same contig. Missing entries (N/A) occur when no sr is observed on the same contig as a focal *cifA*. The length of each contig is reported, with "complete" denoting cases where fully circularized genomes are available. Across all *cif* alleles, we identified 33 whose nearest sr is an sr3WO (sr3) allele, including 21 *cifA*$_{[T1]}$ alleles. Of these, 20 *cifA*$_{[T1]}$ alleles occur within 7,477–13,770 bp of their nearest sr3 allele, while the next closest is more than 200 kb away. *cif* and sr Type copies in each genome are distinguished with an underscore and a number denoting the copy. Plausibly associated sr3 and *cifA*$_{[T1]}$ alleles presented in Figs. S3 and S5 are in bold.

**Table S17. Plausibly associated *cif*$_{[T1]}$-sr3 alleles in *w*Mel-like and *w*Ri-like *Wolbachia* genomes.** Associations between *cif*$_{[T1]}$ and sr3 alleles are particularly common on contigs that encode both a *cif* operon and a *Wovirus*-typing allele. This table presents the 20 *cif*$_{[T1]}$ copies in which *cifA*$_{[T1]}$ is within 14 kb of the nearest sr allele (see Table S14). In all cases, *cifB*$_{[T1]}$ lies 5′ of *cifA*$_{[T1]}$; hence, we report the distances between *cifB*$_{[T1]}$ and the associated *sr3* allele, as well as the number and type of intervening genes based on NCBI annotations. Contig lengths are also provided. Phylograms for these *cif*$_{[T1]}$ and sr3 alleles are compared in Fig. S5.

**Movie S1. Omega mapped on a CifA$_{w\mathrm{Mel[T1]}}$ AlphaFold2 structure.** Colors indicate ln ($\omega$) values: dark blue (< -1), light blue ($-1 \le x < -0.2$), gray ($-0.2 \le x \le 0.2$), yellow ($0.2 < x \le 1$), and orange (> 1).

**Movie S2. Omega mapped on a CifB$_{w\mathrm{Mel[T1]}}$ AlphaFold2 structure.** Colors indicate ln ($\omega$) values: dark blue (< -1), light blue ($-1 \le x < -0.2$), gray ($-0.2 \le x \le 0.2$), yellow ($0.2 < x \le 1$), and orange (> 1).

**Movie S3. HHPred domains mapped on a CifB$_{w\mathrm{Mel[T1]}}$ AlphaFold2 structure.** Colors indicate PD-(D/E)XK nuclease domains in green and a deubiquitinase domain in pink.

**Movie S1-S3 Access:**

https://www.dropbox.com/scl/fo/o9go0dlgdvfwk73473c1n/AC_u4G8ivvmcE8nKrR-adPI?rlkey=hr8wuvpt4flpohzy1edptru9h&st=z93acrez&dl=0
